# MINT infers the latent single-cell spatial transcriptome from paired histology

**DOI:** 10.64898/2026.09.16.751717

**Authors:** Liying Chen, Zhongmin Zhang, Bing Wang, Wenle Ren, Hongke Peng, Shihong Lu, Mengyuan Xu, Yunhe Zhou, Xiongjian Luo, Tong Ye, Qi Zhu, Feng Yan, Luyi Tian

**Affiliations:** Guangzhou National Laboratory, Guangzhou, China; School of Biomedical Engineering, Shenzhen Campus of Sun Yat-sen University, Shenzhen, China; GMU-GIBH Joint School of Life Sciences, Guangzhou Medical University, Guangzhou, China; Lead Healthcare.AI (Guangzhou) Co., Ltd., Guangzhou, China; Department of Neurological Surgery, The Second Affiliated Hospital of Zhejiang University School of Medicine, Hangzhou, China

**Author notes:** Corresponding author(s). E-mail(s): chen.

**Keywords:** Spatial Transcriptomics, Single-cell Resolution, Histology, Optimal Transport, Morphological Features

## Abstract

Sequencing-based spatial transcriptomics captures transcriptome-wide molecular information, yet the corresponding cell-resolved tissue map remains incomplete. Histology records dense cellular morphology, tissue architecture and local context. Here, we present MINT (Morphological Integrative Network for Transcriptomics), which combines this latent cellular organisation with sample-specific spatial measurements to infer tissue-wide, cell-resolved transcriptomes without an external single-cell reference. MINT reconstructs expression from sparse Slide-tags profiles across unmatched adjacent sections; its shared nucleus-centred representation also resolves pooled Visium measurements into cell-level profiles. MINT improved Slide-tags alignment and expression recovery under controlled nuclear loss and cross-section perturbation. It also outperformed reference-free Visium disaggregation methods against Xenium ground truth and was validated with Visium HD sequencing. In human lung adenocarcinoma, MINT reconstructed the tumour immune microenvironment cell by cell and resolved tertiary lymphoid structures and their maturity. These results establish paired histology as a quantitative, sample-intrinsic complement to molecularly rich but cellularly incomplete spatial-transcriptomic assays.

## 1 Introduction

Tissues are organised into spatially defined cellular communities whose gene-expression programmes are shaped by local cell–cell interactions and microenvironmental cues [1–3]; a given cell type can adopt distinct transcriptional states across niches. Resolving expression at single-cell resolution within intact tissue is therefore essential, yet these state differences are lost when expression is averaged within a spot or cells are dissociated from their spatial context. Current spatial-transcriptomic platforms do not provide a whole-transcriptome profile for every individual cell in a tissue [4]. Imaging-based methods such as MER-FISH [5], seqFISH+ [6] and Xenium [7] reach subcellular resolution but measure predefined gene panels. Sequencing-based methods provide transcriptome-wide measurements but leave different components of the cellular map unresolved: spot- and bead-based platforms pool cells within each measurement, whereas fine-grained arrays resolve subcellular bins that must still be grouped into cells by image segmentation. Across sequencing-based assays, molecular breadth therefore does not ensure a complete cell-level map.

Slide-tags [8] presents a distinct form of incomplete cellular coverage. It spatially barcodes intact nuclei within a tissue section before recovery for single-nucleus RNA sequencing, so each recovered nucleus retains both a spatial barcode and a whole-transcriptome profile – a combination of single-cell resolution and unbiased transcriptome coverage that spot-based platforms pool away and imaging panels predefine. A major practical limitation is nuclear recovery: most barcoded nuclei are lost, leaving a sparse molecular map. We asked whether routine H&E histology from an adjacent serial section could extend this measurement without modifying the Slide-tags chemistry. The image supplies a dense census of nuclei and their tissue context from the same tissue block, but the two sections contain independent nuclei in unequal numbers, lack cell-level correspondence and differ through non-rigid deformation. Reconstructing the missing cellular map therefore requires joint cross-modal alignment and sparse-to-dense molecular inference. To our knowledge, this task is not addressed by spatial-alignment methods that assume transcriptomics on both sections, matched cell numbers or same-section correspondence [9–11], or by Slide-tags reconstruction methods that require an external single-cell reference or do not reconstruct expression from the section itself [12, 13].

Visium presents a complementary and widely used form of incomplete cellular assignment. Each 55-*µ*m spot pools transcripts from many cells [14], whereas fine-grained arrays such as Visium HD and Stereo-seq require image-based cell segmentation, often using segmented nuclei as anchors, to aggregate subcellular bins into cellular profiles [15, 16]. Reference-based deconvolution returns each spot’s cell-type composition from a matched scRNA-seq reference [17–22]; even where per-cell profiles are synthesised, they are drawn from that reference rather than read from the section itself [12], so cell states present in the tissue but absent from the reference cannot, by construction, be recovered. Reference-free methods that use paired H&E to predict expression at finer resolution [23–31] anchor predictions to geometric tiles, neighbouring-cell expression or nuclear appearance alone. Methods that directly predict single cells, such as GHIST [32], are supervised by an imaging-based subcellular measurement from the same tissue.

Here, we present MINT (Morphological Integrative Network for Transcriptomics), a framework that uses paired histology as a quantitative complementary measurement for cell-level spatial inference (Fig. 1). MINT combines the dense cellular organisation recorded in histology with sample-specific spatial measurements, without an external scRNA-seq reference. In Slide-tags, it aligns sparse spatially indexed profiles to an adjacent H&E section and infers expression for unsequenced histological nuclei with a morphology-conditioned generative model; we establish this reconstruction under controlled nuclear loss and cross-section perturbation, then apply it to mouse organs and clinical lung adenocarcinoma and glioblastoma. In Visium, the same nucleus-centred image representation resolves pooled spot measurements into constituent cellular profiles under spot-level supervision, tested against Xenium and Visium HD at cell level. Both modes use each segmented nucleus as the spatial and morphological anchor for one cell-level prediction and return cell-resolved spatial expression maps – one principle tested in two independent measurement settings.

**Fig. 1.**
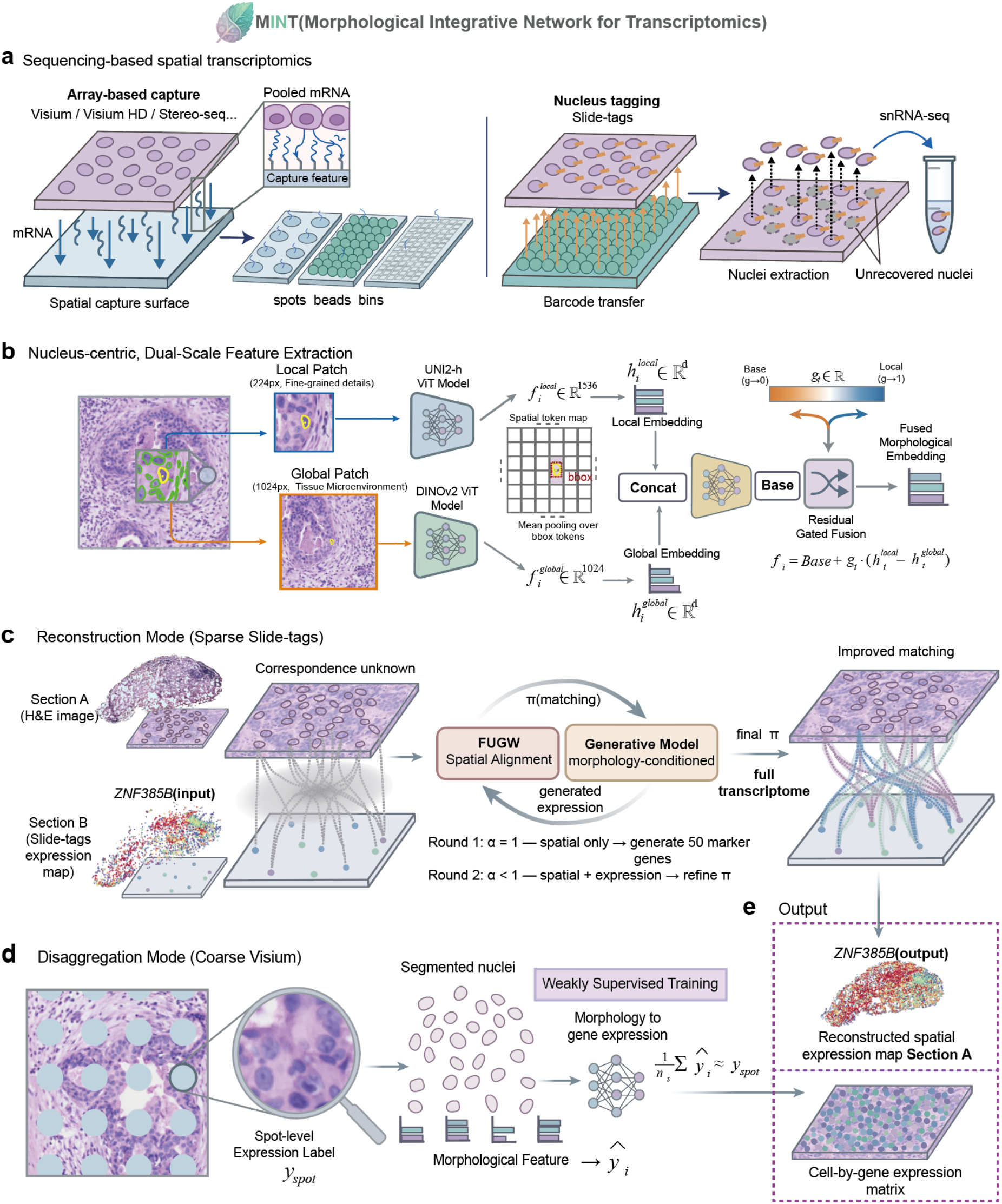
MINT couples paired histology to spatial sequencing to generate cell-resolved expression data. **a**, Two measurement configurations in sequencing-based spatial transcriptomics. Array-based capture measures transcripts aggregated in spatial spots, beads or bins rather than in individual cells. Visium, Visium HD and Stereo-seq are shown as examples of array-based capture formats; the disaggregation mode evaluated here uses pooled Visium measurements, with Visium HD providing a near-single-cell sequencing reference. In Slide-tags, barcodes transfer from the array to nuclei in situ; only extracted nuclei enter snRNA-seq, leaving a subset of tagged nuclei unrecovered. **b**, Nucleus-centric dual-scale feature extraction. For each segmented nucleus, a local patch (224 pixels) is encoded with UNI2-h and a larger contextual patch (1,024 pixels) with DINOv2. Spatial tokens overlapping the nuclear bounding box are mean-pooled at each scale; projected local and global features are combined by a residual-gated fusion module to form a fused morphological embedding. **c**, Reconstruction mode for sparse Slide-tags. An H&E section (section A) is paired with a sparse Slide-tags expression map from an adjacent section (section B). FUGW establishes and refines a soft cross-section correspondence, which provides supervision for a morphology-conditioned generative model. The example shows *ZNF385B* in the sparse input and after reconstruction on section A. **d**, Disaggregation mode for coarse Visium. A segmented nucleus anchors each cell-level prediction from the fused morphological features; profiles of cells assigned to the same spot are constrained in aggregate to recover the measured spot expression. **e**, Both modes yield cell-resolved expression data, represented as a spatial expression map and a cell-by-gene matrix with one profile and one spatial coordinate per nucleus-anchored cell.

## 2 Results

### 2.1 MINT uses paired histology to complete cellular maps from spatial sequencing

MINT addresses the two measurement settings in which sequencing-based assays leave cells unresolved (Fig. 1a). In both, it takes a routine H&E section from the same tissue block together with the measurement the assay does provide, and returns one whole-transcriptome profile and one spatial coordinate for every segmented nucleus (Fig. 1e). The segmented nucleus is the unit of inference throughout: it is encoded from the image at two spatial scales, and that representation is mapped to expression by a head matched to the molecular supervision the assay affords – a soft cross-section correspondence for Slide-tags, an aggregation constraint within each spot for Visium. Neither setting uses an external single-cell reference.

MINT encodes each segmented nucleus at two complementary scales, because a cell’s transcriptional state reflects both its intrinsic identity and the niche signals it receives from neighbouring cells. A local patch (~25 *µ*m) captures nuclear shape, size and chromatin texture, whereas a wider field (~ 110 *µ*m) captures glandular architecture, stromal organisation, inflammatory infiltrates and local cell density (Fig. 1b). Both are extracted from nucleus-centred image regions and fused by a residual gate (Methods; segmentation and stain-normalisation quality controls in Supplementary Fig. S1). On pseudo-Visium data, residual-gated fusion outperformed either scale alone and naïve concatenation, and nucleus-centred features produced more coherent tissue domains than fixed-grid features (Extended Data Fig. 1).

For Slide-tags, MINT aligns the sparse transcriptomic section to the dense H&E section (Fig. 1c). Fused unbalanced Gromov–Wasserstein optimal transport [33–35] accommodates the density imbalance and serial-section deformation, then iteratively incorporates an expression feature predicted from histology. The resulting soft correspondence supervises a morphology-conditioned generative model, which samples a distinct expression profile for every histological nucleus – preserving the cell-to-cell variability that each captured nucleus independently measures – to generate a dense, whole-transcriptome single-cell map (Fig. 1c,e).

For Visium, MINT maps the same fused image representation to cell-level expression profiles under spot-level supervision (Fig. 1d). Each segmented nucleus serves as the image anchor for one predicted cell, which is assigned to a spot by its nuclear centroid. The cell profiles within each spot are constrained in aggregate to recover the measured spot expression, while morphologically distinct cells can receive distinct profiles. The two modes therefore share one component and differ in one: the nucleus-centred dual-scale encoder is the same, whereas the supervision and the decoder it drives are matched to the assay – a cross-section transport plan supervising a generative decoder for Slide-tags, a within-spot aggregation constraint supervising a deterministic decoder for Visium. Both yield cell-resolved spatial expression data with one profile and one coordinate for each nucleus-anchored cell (Fig. 1e).

### 2.2 MINT recovers expression across sparse, adjacent sections under known ground truth

We tested MINT in a controlled pseudo-Slide-tags experiment with known single-cell truth (Methods). Starting from a Xenium mouse-brain section, we generated a sparse transcriptomic section by removing nuclei at graded rates and applying random and smooth spatial perturbations (Fig. 2a). The paired dense section supplied H&E-derived nuclear features. Alignment accuracy was evaluated against each nucleus’s known location, and expression recovery was evaluated on genes excluded from alignment.

**Fig. 2.**
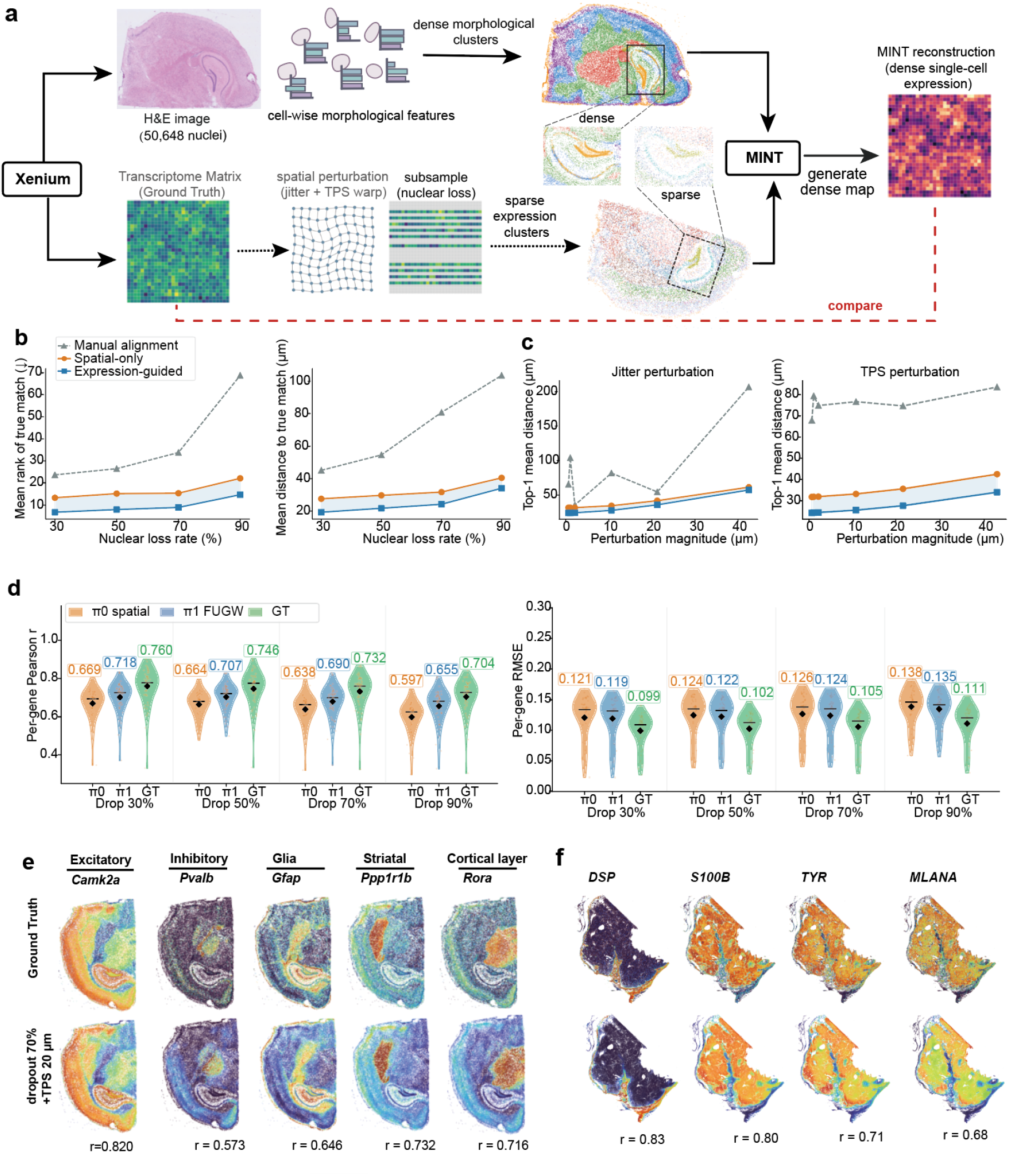
MINT recovers expression across sparse, adjacent sections under known ground truth. **a**, Validation schematic. MINT segments nuclei, extracts dual-scale morphological features and clusters the dense H&E reference section (section A). The sparse transcriptomic section (section B) was generated from the same tissue by nuclear loss (70%), Gaussian jitter and thin-plate-spline deformation, shown for a mouse hippocampal region. After alignment, MINT generated predictions for every nucleus in section A, which were evaluated against Xenium per-nucleus expression (Methods). **b**, Alignment accuracy versus nuclear loss rate (mouse brain), comparing manual landmark alignment, the spatial-only plan (*π*_0_) and the expression-guided plan (*π*_1_): mean rank of the true counterpart (left) and Top-1 mean distance (right) versus nuclear loss rate (Methods). **c**, Robustness to spatial perturbation at a fixed 70% nuclear loss, for the same three methods: Top-1 mean distance under graded Gaussian jitter (left) and under graded TPS deformation (right) (Methods). **d**, Per-gene Pearson correlation (left) and per-gene RMSE (right) of MINT predictions against Xenium ground truth on 50 spatially variable genes held out from alignment (ranks 51–100 by Moran’s *I*, disjoint from the top-50 genes used to guide the alignment), across four nuclear loss rates (30/50/70/90%) and the *π*_0_ (spatial-only), *π*_1_ (expression-guided) and GT (ground-truth-alignment) stages. **e**, Spatial expression of five cluster-defining mouse-brain marker genes (*Camk2a, Pvalb, Gfap, Ppp1r1b, Rora*) in the ground truth (top) and the MINT prediction with 70% nuclear loss and TPS ~20 *µ*m perturbation (bottom); per-gene Pearson *r* is shown beneath each prediction. Cells are coloured by expression level with the colour scale clipped at the 1st–99th percentiles. **f**, Human skin melanoma (Xenium, 80% nuclear loss): spatial expression of four marker genes (*DSP, S100B, TYR, MLANA*) in the ground truth (top) and the MINT prediction (bottom), with per-gene Pearson *r*.

MINT first estimates a spatial-only correspondence from tissue geometry. This initial plan captures global tissue architecture but lacks molecular information. MINT then predicts a compact panel of spatially variable marker genes from morphology and incorporates these predictions as an additional similarity signal, yielding a sharper expression-guided correspondence (Extended Data Fig. 2c). At 70% nuclear loss, this refinement reduced the median distance between each matched profile and its true nucleus from 27.9 *µ*m for the spatial-only plan (*π*_0_) to 21.5 *µ*m for the expression-guided plan (*π*_1_; Extended Data Fig. 2d), approximately two cell diameters. The improvement persisted in mean matching distance from 30% to 90% loss (Fig. 2b). To isolate the contribution of expression information, the figures show one guided round (*π*_1_); the default MINT correspondence uses a second refinement round (*π*_2_; Methods).

The expression-guided alignment remained more accurate than the spatial-only baseline under graded random and smooth perturbations (Fig. 2c). These smooth deformations approximate the gradual local distortions introduced by cryosectioning or paraffin embedding in serial sections [36]. Manual landmark alignment was less accurate and more variable across annotators (Fig. 2b,c; Methods).

Improved alignment increased cross-section expression recovery by the generative reconstruction model. Expression-guided alignment (*π*_1_) improved prediction accuracy for nearly all genes held out from alignment relative to the spatial-only baseline (*π*_0_) and recovered approximately half of the gap to the ground-truth-alignment ceiling (Fig. 2d). The recovered markers preserved their spatial organisation: the excitatory-neuron marker *Camk2a*, for example, retained its layer-specific cortical pattern (per-gene *r* beneath each gene; Fig. 2e). The reconstruction also recovered cell-type structure: predicted cluster maps matched ground truth at every loss level, including 90%, and cluster-defining markers separated cleanly between brain regions (Extended Data Fig. 2e,f).

A separate human skin melanoma dataset reproduced the alignment and expression-recovery gains (Supplementary Table S1). At 80% nuclear loss, expression-guided alignment reduced the mean matching distance from *π*_0_ to *π*_1_ by approximately one-third and recovered cell-type-specific spatial expression (Fig. 2f). Robustness to Gaussian jitter and thin-plate-spline deformation also reproduced the mouse-brain result (Supplementary Fig. S2; Supplementary Table S3).

### 2.3 MINT restores interpretable tissue architecture in real Slide-tags data

We applied reconstruction mode to Slide-tags data generated for this study from the olfactory bulb, kidney and brain. MINT predicted a profile for every segmented nucleus, increasing map density by 3- to 10-fold relative to the captured nuclei (Fig. 3).

**Fig. 3.**
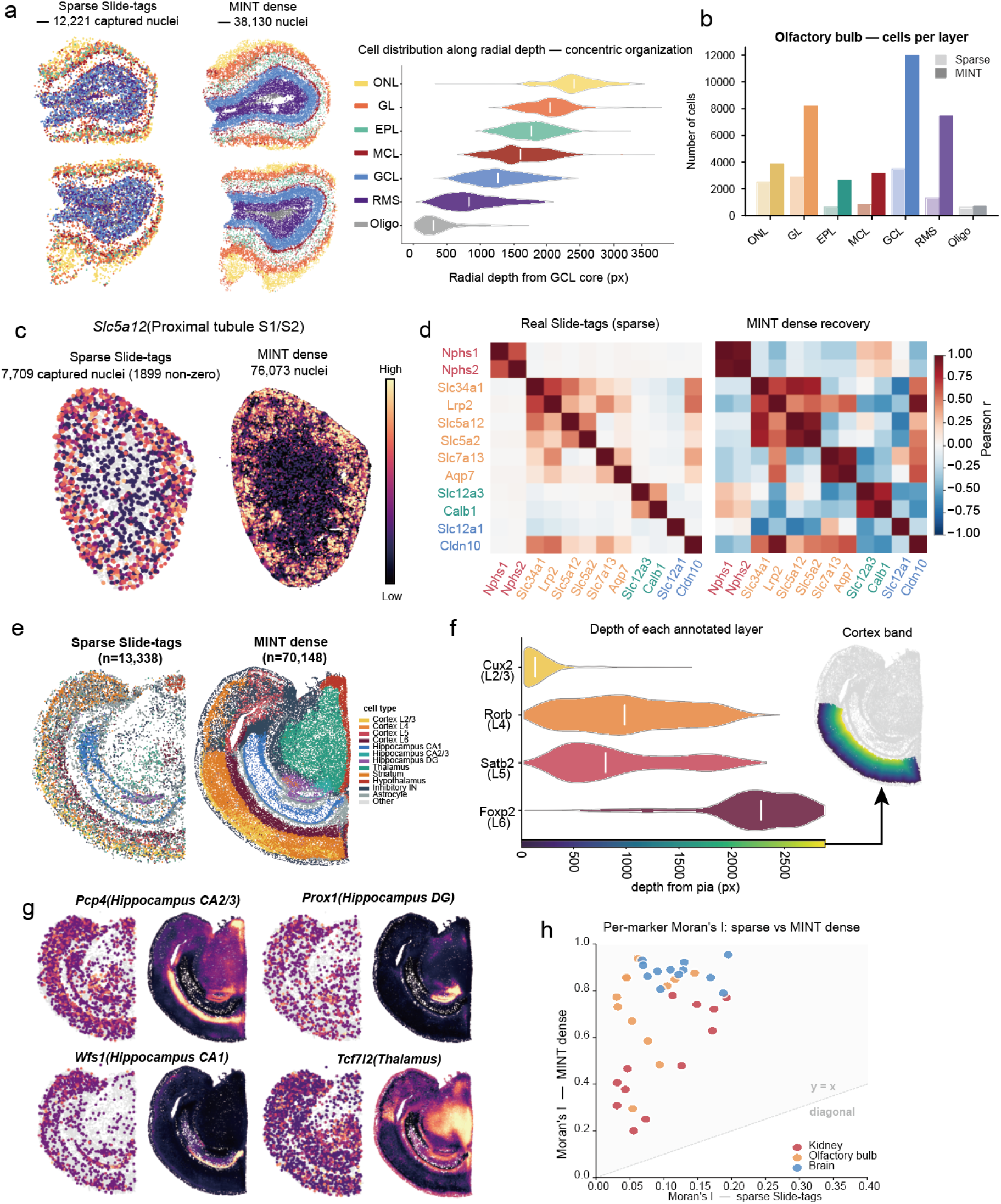
MINT restores interpretable tissue architecture in sparse Slide-tags data. **a**, Mouse olfactory bulb: sparse Slide-tags versus dense MINT reconstruction, with cell distribution along radial depth showing the concentric laminar organisation. **b**, Olfactory bulb cell counts per layer, sparse Slide-tags versus MINT dense reconstruction. **c**, Kidney single-gene recovery (*Slc5a12*): sparse Slide-tags (7,709 captured nuclei) versus MINT dense map (76,073 cells). **d**, Marker co-expression heatmap (Pearson *r*), real Slide-tags versus MINT dense reconstruction, for kidney nephron-segment markers. **e**, Mouse brain: sparse Slide-tags (13,338 captured nuclei) versus MINT dense (70,148 cells), coloured by cell type (spatially aware transcriptomic clusters of the reconstructed cells; Methods). **f**, Mouse brain cortical-depth distribution of laminar markers (*Cux2* layer 2/3, *Rorb* layer 4, *Satb2* layer 5, *Foxp2* layer 6) against depth from the pia. **g**, Mouse brain hippocampal subfields and thalamus, sparse Slide-tags versus MINT dense reconstruction (*Wfs1* CA1, *Pcp4* CA2/3, *Prox1* dentate gyrus, *Tcf7l2* thalamus). **h**, Cross-tissue spatial coherence: per-marker Moran’s *I* in the sparse data versus the MINT reconstruction for all three organs (points above the diagonal indicate increased spatial autocorrelation). Per-tissue detail in Supplementary Figs. S3–S6.

Per-marker Moran’s *I* increased in all three tissues (Fig. 3h), and markers not detected in an anatomical layer in the sparse data were not predicted in that layer after reconstruction – a specificity control against simple spatial smoothing. Because real Slide-tags data carry no dense single-cell reference, we assessed the reconstructions throughout by anatomical fidelity, marker co-expression, cross-layer specificity and spatial coherence (Methods).

In the olfactory bulb, clustering of sparse captured nuclei yielded a discontinuous map in which thin layers were not resolved, whereas MINT recovered continuous, correctly ordered concentric rings corresponding to the outer nerve, glomerular, mitral-cell and granule-cell layers (Fig. 3a,b; Supplementary Fig. S3). The mitral-cell layer was recovered as a distinct band. Marker co-expression between the sparse and dense maps was consistent (Pearson *r* = 0.64; Supplementary Fig. S3), supporting preservation of molecular relationships as spatial organisation became denser.

In the kidney, where captured nuclei were sparsest and MINT increased cell density about 10-fold, the reconstruction recovered segment-restricted expression across the nephron. The proximal-tubule marker *Slc5a12*, which was sparsely sampled in the captured nuclei, was recovered as a continuous cortical band (Fig. 3c), and nephron-segment markers occupied their expected radial positions along the cortico-medullary axis (Supplementary Fig. S4). Markers from the same nephron segment formed distinct on-diagonal co-expression blocks, whereas markers from different segments separated. This structure was resolved in the MINT dense map while remaining consistent with the sparse captured nuclei (Fig. 3d).

In the brain, captured nuclei were too sparse and scattered to reveal fine tissue architecture (Fig. 3e; Supplementary Figs. S5 and S6). We annotated the reconstruction by spatially aware clustering (BANKSY) of predicted transcriptomes and local spatial context, followed by canonical-marker labelling (Methods). MINT recovered cortical layers in their expected order from the pia and resolved the hippocampal subfields and thalamus. Region-defining markers occupied their expected territories: *Wfs1* in CA1, *Pcp4* in CA2/3, *Prox1* in the dentate gyrus and *Tcf7l2* in the thalamus (Fig. 3g).

### 2.4 MINT resolves tumour microenvironments from paired clinical histology

We applied reconstruction mode to Slide-tags data generated for this study from six human lung specimens: five early lung adenocarcinomas (P1–P5, including two driver-mutant cases) and one non-malignant pulmonary granuloma as a positive control for organised lymphoid tissue (Supplementary Table S4). Each Slide-tags section was paired with an adjacent H&E section. We used the dense reconstructions to examine tertiary lymphoid structures (TLS), organised lymphoid aggregates associated with prognosis and immunotherapy response [37].

A TLS is organised into molecularly distinct compartments – a B-cell follicle with follicular dendritic cells, surrounded by a T-cell zone – and its maturity is read from markers such as CR2 and FCER2 that only transcriptomic information provides. The lymphocyte aggregate is visible to a pathologist in routine H&E, yet sparse Slide-tags nuclei sample it too thinly to resolve these compartments. MINT recovered this structure by assigning a transcriptional identity to each nucleus in the H&E image. The resulting map resolved the follicle as an organised, cell-typed structure, whereas sparse capture showed only scattered nuclei (Fig. 4b; Extended Data Fig. 3d). The recovered spatial organisation followed histological morphology rather than a simple interpolation of sparse transcriptomic coordinates.

**Fig. 4.**
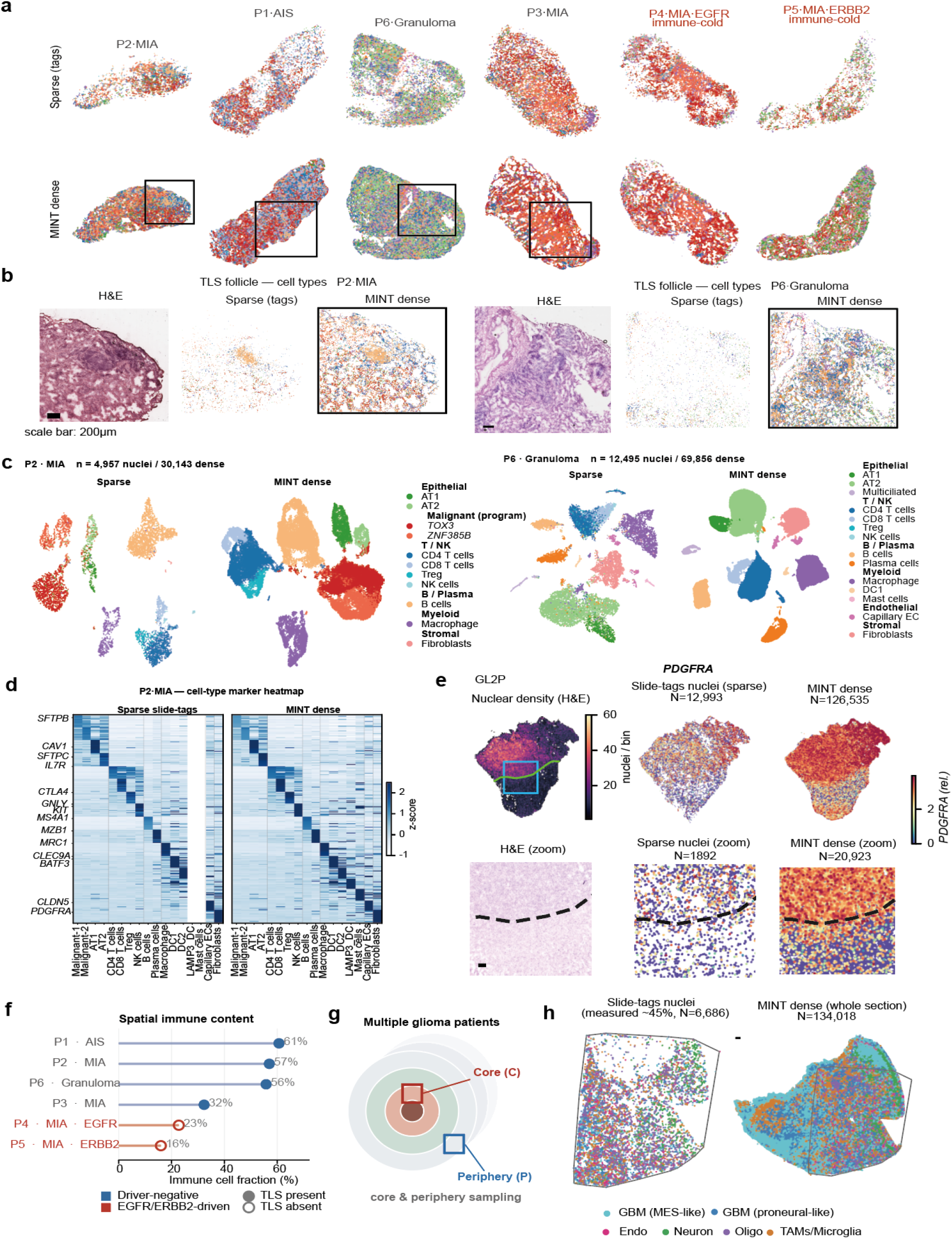
MINT resolves tumour microenvironments from paired clinical histology and sparse Slide-tags data. **a**, Sparse Slide-tags captured nuclei versus dense MINT reconstruction across the six lung samples (P1–P6), shown as spatial cell-type maps; captured-nucleus and dense cell counts are annotated, and the cell-type legend groups epithelial, T/NK, B/plasma, myeloid, endothelial and stromal lineages. **b**, Local follicle views for a TLS-bearing minimally invasive adenocarcinoma (P2) and the non-malignant granuloma (P6), showing adjacent-section H&E, sparse captured nuclei and the MINT dense cell-type reconstruction (scale bar, 200 *µ*m). **c**, UMAP embeddings of sparse captured nuclei and the MINT dense reconstruction for P2 and P6, with marker-defined cell-type annotations. **d**, Cell-type marker-signature heatmap for P2, comparing sparse captured nuclei and the MINT dense reconstruction (per-cell-type *z*-score). **e**, Glioblastoma peritumoural section GL2P: H&E-derived nuclear-density map, sparse captured nuclei and whole-section MINT dense reconstruction, with a local *PDGFRA*-coloured view (scale bar, 200 *µ*m). **f**, Spatial immune content per lung sample (immune cell fraction), annotating TLS-present versus TLS-absent and driver-mutation status as a descriptive summary of the reconstructed maps and clinical metadata. **g**, Glioblastoma sampling design: tumour core (C) and peritumoural margin (P) across patients. **h**, Glioblastoma core sample GL3C, in which Slide-tags covered about 40% of the section: sparse captured nuclei versus MINT dense reconstruction across the whole section, shown as spatial cell-type maps (scale bar, 200 *µ*m). Per-sample lung maps and TLS-maturity quantification in Extended Data Figs. 3 and 4 and Supplementary Figs. S7–S8; glioblastoma detail in Extended Data Figs. 5–6.

In a TLS-bearing minimally invasive adenocarcinoma (P2), captured nuclei from the immune compartment were too sparse and discontinuous to delineate a follicle. In UMAP space, the dense reconstruction retained the marker-defined cell-type structure of the captured nuclei, with reconstructed cells integrated within the measured manifold (Fig. 4c; Extended Data Fig. 4). On the tissue section, the reconstruction resolved a mature TLS comprising a B-cell follicle with an FDC network and a surrounding T-cell zone (Fig. 4a,b; Extended Data Fig. 3a,d). Follicular dendritic cells were 4.4-fold enriched within the B-cell follicle core (Extended Data Fig. 3a).

Marker expression and cohort-level composition supported the cell-type annotations used to define these structures. Marker-signature heatmaps in P2 showed the same lineage-specific marker blocks in the captured nuclei and the dense reconstruction, including rare annotated lineages represented by fewer than 15 captured nuclei that became interpretable only after reconstruction (Fig. 4d). Across the cohort, broad lineage composition remained similar before and after reconstruction, indicating that MINT densified the tissue without substantially altering the cellular composition of individual samples (Extended Data Fig. 4a). Per-sample marker maps provided gene-level spatial context for epithelial, lung-lineage and immune-associated programmes (Supplementary Figs. S7, S8).

Across the cohort, MINT recovered the maturity spectrum of these structures (Extended Data Fig. 3a). The adenocarcinoma in situ (P1) carried the highest immune content of the cohort and formed multiple discrete, concentric follicles (Fig. 4a,f; Extended Data Fig. 3). A third minimally invasive case (P3) contained a focal lymphoid aggregate with partial FDC evidence (*CR2* detected but *FCER2* below the captured-nucleus detection threshold). This pattern was consistent with a maturing follicle supported by partial FDC evidence, rather than a fully mature TLS. The non-malignant granuloma (P6) contained FDC-enriched, focally organised B-cell aggregates of comparable magnitude to the cancer-associated TLS (Fig. 4a,b; Extended Data Fig. 3), showing that MINT resolves organised lymphoid tissue in both cancer-associated and non-malignant settings.

The two driver-mutant adenocarcinomas lacked organised lymphoid architecture. P4 (*EGFR*) contained scattered lymphocytes forming at most an immature aggregate without follicular organisation, whereas P5 (*ERBB2*) was B-cell-poor and contained no follicular structure (Fig. 4a,f; Extended Data Figs. 3c,d and 4; Supplementary Figs. S7 and S8). In both cases, MINT preserved the absence of organised follicles observed in the tissue. These immune-cold phenotypes were consistent with less-inflamed microenvironments reported in EGFR- and ERBB2-mutant lung adenocarcinoma [38, 39].

Paired clinical H&E and sparse adjacent-section transcriptomics thus delineated TLS architecture and maturity at single-cell resolution.

### 2.5 MINT recovers single-cell spatial structure in glioblastoma

We then applied the reconstruction mode to Slide-tags data generated for this study from three glioblastoma patients, sampling the tumour core and, in two patients, the peritumoural margin (Fig. 4g; Supplementary Table S4). Each section was paired with an adjacent H&E image. MINT reconstructed dense single-cell maps at approximately 6- to 20-fold higher density, and the reconstructed cells remained integrated with the marker-defined manifold of captured nuclei (Extended Data Fig. 5a,b). Unlike the discrete lineages of the lung immune microenvironment, glioblastoma malignant cells occupy a continuous transcriptional spectrum, and several defining programmes are only partly represented in nuclear morphology [40].

The reconstruction was most informative for spatial cellular architecture. In a peritumoural section (GL2P), the H&E image contained an abrupt boundary in nuclear density that sparse captured nuclei were too few to trace. MINT reconstructed the tissue on both sides of this boundary at single-cell resolution, resolving the density transition and the local distribution of malignant and microenvironmental cells that the sparse map left as isolated points (Fig. 4e).

MINT also extended predictions into regions without recovered Slide-tags nuclei. In one core sample (GL3C), Slide-tags covered only about 40% of the sectioned tissue, whereas H&E covered the full section. MINT generated a cell-resolved map across the remaining tissue whose spatial organisation followed the visible histological structure (Fig. 4h). Molecular accuracy in this region could not be tested directly because no captured nuclei provided a reference.

The same densification turned the malignant and microenvironmental programmes from scattered points into spatial fields. In the two tumour-core sections shown, the perinecrotic hypoxia programme resolved into contiguous foci where the captured nuclei had shown only isolated high-scoring cells, and the mesenchymal-like (MES) programme was elevated over the same territory (Extended Data Fig. 6a) – the arrangement expected if the mesenchymal state is sustained in hypoxic tissue bordering necrosis [40]. The OPC-like/proneural programme [40, 41] and the principal microenvironmental compartments were likewise resolved as spatially continuous fields (Extended Data Figs. 5c, 6).

### 2.6 The MINT image representation resolves cells pooled by Visium

The same nucleus-centred representation that fills unsequenced nuclei in Slide-tags also resolves the opposite failure mode of array capture, in which transcripts from many cells pool within a spot. To measure single-cell accuracy directly in this setting, we benchmarked MINT against three reference-free comparators – iStar [27], Thor [28] and sCellST [29] – on pseudo-Visium data built from five Xenium sections across four tissue types by pooling measured single-cell expression into Visium-sized spots, which preserves the true cellular profile underlying every spot (Supplementary Table S1). All four methods received the same spot aggregates and the same paired H&E image, were applied to the same H&E-segmented nuclei and were compared in a common CP10k+log1p space, so every prediction corresponds directly to its Xenium reference and every method faced the same task: resolving measured spot profiles from paired histology without an external reference (Methods).

MINT achieved the highest single-cell prediction accuracy on all five tissue sections (Fig. 5e,f; Extended Data Fig. 7a,b; per-tissue, per-method values in Supplementary Table S2). It had the highest cell-wise profile correlation, the lowest per-gene RMSE and better performance than the next-best method for most genes tested (Fig. 5e; Extended Data Fig. 7a). MINT assigned distinct expression profiles to cells within the same spot, capturing within-spot heterogeneity against matched Xenium profiles; the spot measurements entered only as a training constraint, and neither cellular coordinates nor spot identity was available to the predictor.

**Fig. 5.**
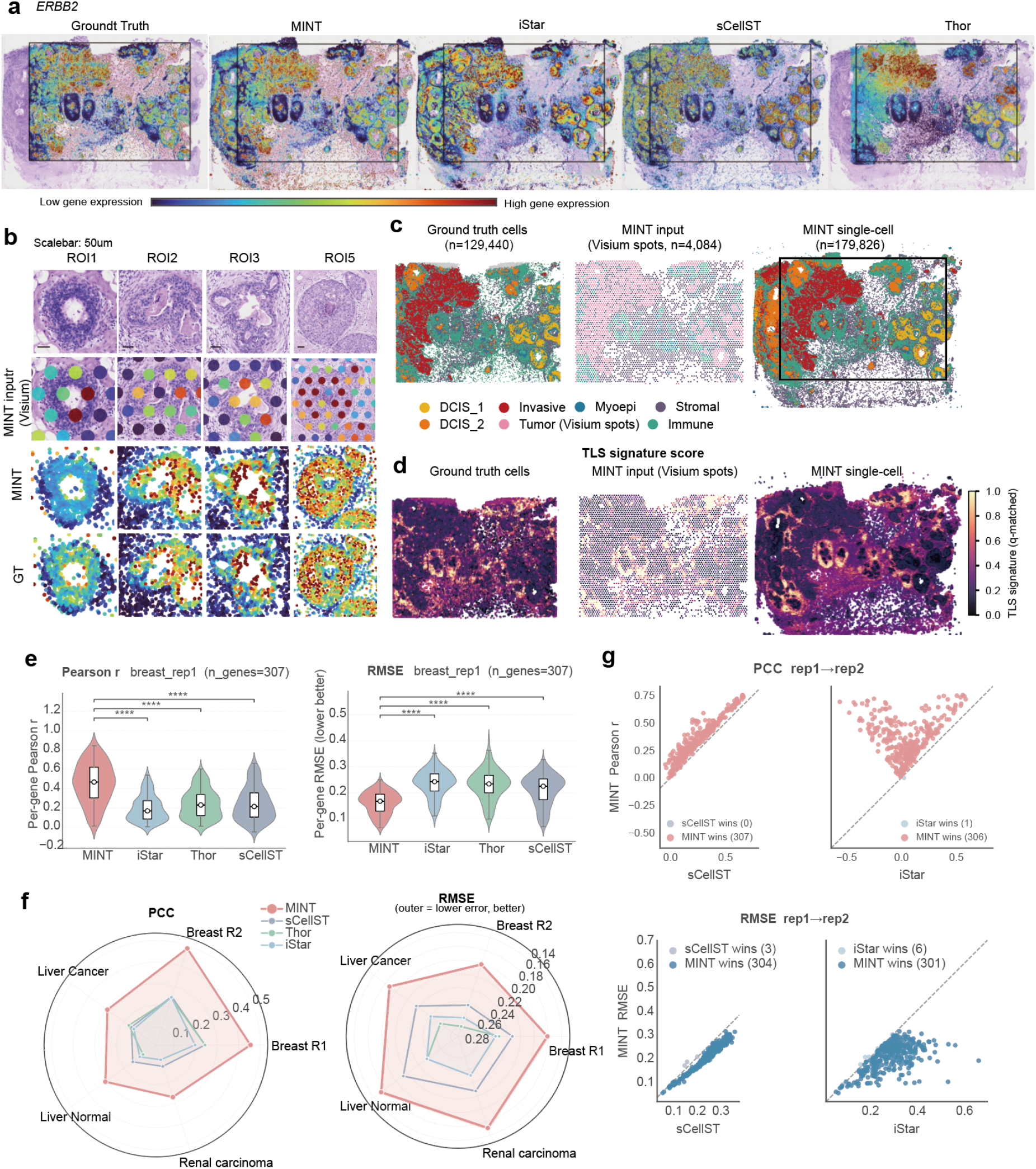
The MINT image representation resolves cells pooled by Visium across five tissues. All methods were trained on the same spot aggregates and applied to the same H&E-segmented nuclei. Predicted cell *i* therefore corresponds directly to Xenium cell *i*, without spatial matching; values are compared in a common CP10k+log1p space (Methods). **a**, Whole-section spatial expression maps of *ERBB2* (HER2) on the H&E for Xenium ground truth, MINT, iStar, sCellST and Thor (breast cancer Rep1). In **a** and **b**, cells are coloured by expression level with the colour scale clipped at the 1st–95th percentiles. **b**, Representative regions of interest (ROIs 1–3 and 5; locations outlined on the whole-section H&E in Extended Data Fig. 8a) comparing H&E, pseudo-Visium spot input, MINT single-cell prediction and Xenium ground truth for *ERBB2*. **c**, Cell-type clustering of the same section in Xenium ground-truth single cells, pooled Visium spot input and MINT single-cell predictions (cell counts annotated). Annotations include ductal carcinoma in situ subtypes (DCIS 1/DCIS 2), invasive carcinoma, myoepithelial, stromal and immune cells. MINT predicted every segmented nucleus across the full H&E section, extending beyond the Xenium-measured field; per-cell accuracy was evaluated only for nuclei matched to Xenium cells. **d**, Spatial maps of a tertiary-lymphoid-structure (TLS) signature score (multi-gene immune signature, Supplementary Table S7) for the same three views (Xenium ground truth, Visium spot input and MINT prediction). **e**, Per-gene Pearson correlation and RMSE distributions (violins) for MINT versus iStar, Thor and sCellST (breast Rep1, *n* = 307 genes); overlaid boxes mark the median (centre line) and interquartile range (Q1–Q3), whiskers extend to 1.5× the interquartile range, and brackets denote two-sided Wilcoxon signed-rank tests versus MINT. **f**, Multi-metric radar across all five tissue sections (per-gene Pearson and RMSE), MINT versus competitors. **g**, Cross-slice generalisation, shown for one of the two directions (train Rep1, predict Rep2): paired per-gene scatter of Pearson and RMSE, MINT versus sCellST and iStar; both directions are quantified in Supplementary Table S6.

At the tissue level, MINT recapitulated Xenium spatial expression patterns across genes (Fig. 5a,b, Extended Data Fig. 8 and Supplementary Figs. S9–S12). In breast cancer, MINT recovered the localised HER2 (*ERBB2*) overexpression pattern in HER2-high epithelial regions and the diffuse immune-cell distribution marked by CD45 (*PTPRC*). MINT preserved these spatial gradients, which competing methods blurred or displaced. This spatial fidelity extended to canonical lineage markers across renal carcinoma, liver cancer and normal liver. Unsupervised clustering of MINT predictions recovered cell-type structure at single-cell resolution, including ductal carcinoma in situ subtypes, invasive carcinoma, myoepithelial, stromal and immune cells; the spot data resolved only a coarse tumour compartment (Fig. 5c). The same predictions reconstructed the spatial distribution of a tertiary-lymphoid-structure signature in agreement with the Xenium ground truth, which the spot-level input could not resolve (Fig. 5d; Supplementary Fig. S13).

The fitted morphology-to-expression mapping can be applied to an additional H&E section of the same tissue without further spatial-transcriptomic profiling. We trained MINT on one breast cancer replicate and predicted the other in both directions (Rep1→Rep2 and Rep2→Rep1), evaluating predictions against Xenium single-cell ground truth in the held-out section (Fig. 5g; Supplementary Table S6). MINT attained the highest mean per-gene Pearson correlation and mean cell-wise profile correlation in both directions. Its mean per-gene RMSE differed from sCellST by less than 2% in both directions, and both nucleus-centred methods outperformed the grid-based iStar [27].

### 2.7 MINT generalises to real Visium tissue and Visium HD validation

To validate disaggregation on genuine Visium sequencing, we analysed a human colon adenocarcinoma section profiled by Visium HD. We aggregated its continuous 2-*µ*m grid into 55-*µ*m spots for training and independently assigned the original bins to cells using H&E-segmented nuclei as spatial anchors, constructing a near-single-cell reference containing ~400 unique molecular identifiers per cell (Supplementary Table S1; Methods). MINT was trained only on the aggregated spot profiles and evaluated against the quality-controlled reference cells using the same in-sample protocol as the pseudo-Visium benchmark.

Across reference cells, per-gene correlation between MINT predictions and measured expression increased with gene-detection rate and exceeded that of a morphology-only nearest-neighbour baseline throughout. The median correlation increased from ~0.28 for sparsely detected genes to ~0.52 for the most readily detected tier (Supplementary Fig. S14d), reaching 0.66–0.86 for well-detected markers, including *PIGR, COL3A1* and *CEACAM5* (Fig. 6a; Supplementary Fig. S15). Lower correlations for sparsely detected genes were consistent with their limited dynamic range and the shallow depth of the HD reference.

**Fig. 6.**
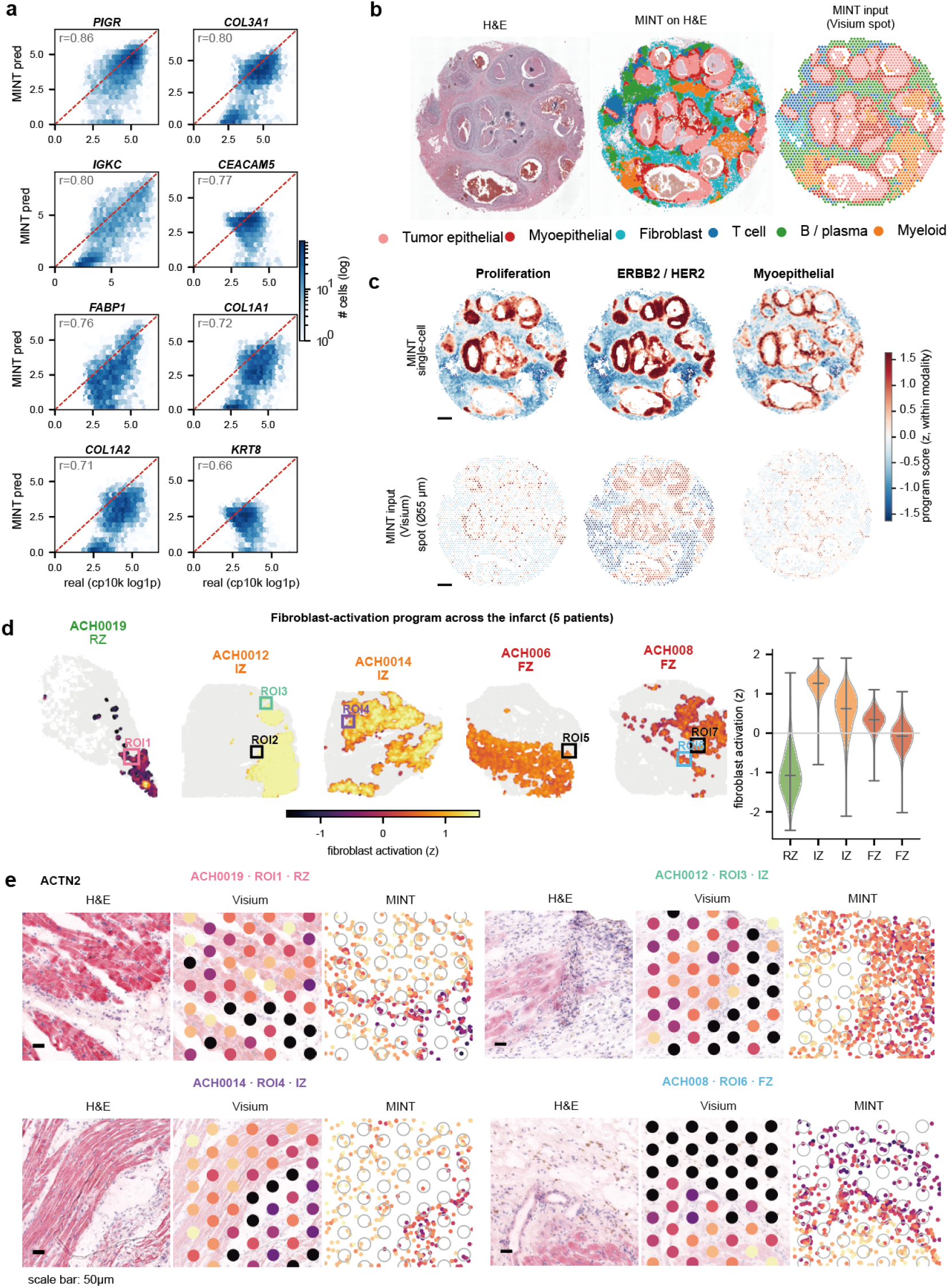
MINT generalises to real Visium tissue and is validated at cell level on Visium HD. **a**, Visium HD human colon cancer, single-cell validation: per-gene scatter of MINT predictions against the near-single-cell HD measurement (CP10k+log1p) for eight representative markers (*PIGR, COL3A1, CEACAM5* and others), across the quality-controlled reference cells, with the Pearson correlation annotated and points coloured by cell density. **b**, Human breast ductal carcinoma in situ (DCIS): H&E, the MINT single-cell cell-type map overlaid on the H&E, and the pooled Visium spot input for the same section (tumour epithelial, myoepithelial, fibroblast, T cell, B/plasma and myeloid). **c**, Breast DCIS cell-state programmes (proliferation, ERBB2/HER2, myoepithelial) at MINT single-cell versus Visium spot resolution (programme score *z*-scored within each modality). **d**, Human myocardial infarction: fibroblast-activation programme (*COL1A1* /*POSTN* /*COL3A1* /*FN1*) across the injury axis in five samples (remote, ischaemic and fibrotic zones), as single-cell spatial maps coloured by programme score with per-sample violins (central bar, median; vertical extent, full data range). **e**, Regions of interest at the infarct border, comparing H&E, Visium spot and MINT single-cell expression of the sarcomeric marker *ACTN2* (a marker of surviving cardiomyocytes). Detailed per-tissue results are in Extended Data Figs. 9 and 10 (olfactory bulb and coronal brain) and Supplementary Figs. S14–S20 (colon Visium HD, breast DCIS, and per-tissue marker maps).

MINT’s single-cell predictions also recovered the colon’s cell-type structure. Clustering the predictions resolved six cell types (epithelial, fibroblast, T cell, B/plasma, myeloid and endothelial), producing clearer separation than that supported by shallow HD counts alone (Supplementary Fig. S14a). In the HD counts, myeloid cells overlapped with the T-cell cluster, whereas MINT resolved a distinct myeloid population with nuclear morphology consistent with the H&E image (Supplementary Fig. S14b,c). The MINT partition subdivided the coarser HD structure. Marker-defined populations showed coherent nuclear-morphology galleries (Supplementary Fig. S16).

### 2.8 MINT resolves spatial structure that Visium spots average away

MINT reads laminar and regional architecture from real Visium tissue at cellular resolution beyond that of 55-*µ*m spots. We applied MINT to two mouse tissues with well-defined anatomical organisation, the olfactory bulb and a coronal brain section, whose expected layout is documented by the Allen Brain Atlas (Methods).

In the olfactory bulb, spatially aware clustering (BANKSY[42]) of MINT predictions recovered continuous, correctly ordered concentric rings corresponding to the outer nerve and glomerular layers, the mitral-cell layer and the inner granule-cell layer. These rings were finer than those recovered from spot data alone (Extended Data Fig. 9a–c). Layer-defining markers occupied their expected radial positions, matching the Allen Brain Atlas (Extended Data Fig. 9d).

In the coronal brain section, MINT resolved principal anatomical domains at cellular resolution, including cortical lamination from superficial layer 2/3 to deep layer 6, the hippocampal CA pyramidal arc and dentate gyrus, thalamic and striatal nuclei, and white-matter tracts (Extended Data Fig. 10a). At single-cell resolution, the hippocampal field resolved the characteristic V shape of the dentate gyrus (Extended Data Fig. 10b). Region-defining markers reproduced their spot-level spatial patterns, reaching a MINT-versus-spot per-gene *r* of up to 0.86 for the oligodendrocyte marker *Mbp* (Extended Data Fig. 10c).

We next examined disease tissues in which Visium spots averaged cellular architecture and state. The analyses used a breast ductal carcinoma in situ (DCIS) section and myocardial infarction tissue.

In the DCIS section, a single 55-*µ*m spot pooled the tumour-epithelial core, thin myoepithelial sheath, and peri-ductal stroma and immune infiltrate into one mixed profile. Overall, 27% of spots spanned at least two predicted cell types, with the greatest mixing at tumour–stroma interfaces. MINT resolved these components into single cells across the full section, including cells between spots, and recovered the concentric duct architecture that spot averages obscure. This architecture included the thin *KRT5* /*KRT14* myoepithelial shell surrounding each tumour-epithelial core (Fig. 6b; Supplementary Fig. S17). Programme maps further resolved this organisation (Fig. 6c). Proliferation and myoepithelial programmes showed little coherent structure in Visium spots but localised to ducts in MINT predictions. The more broadly distributed ERBB2/HER2 programme was already spatially patterned at spot resolution and was sharpened by MINT. Predictions on held-out spots remained accurate for hallmark genes (*ERBB2, r* = 0.81; *EPCAM, r* = 0.77; Supplementary Figs. S17 and S18; Methods).

Kuppe et al.[43] mapped cellular reorganisation in infarcted human heart by integrating single-nucleus RNA-seq, single-nucleus ATAC-seq and Visium. Using Visium and H&E alone, MINT recovered single-cell remodelling programmes in this tissue. Across five sections spanning remote, ischaemic and fibrotic zones, MINT resolved the major cardiac lineages at single-cell resolution (cardiomyocyte, fibroblast, endothelial, immune and pericyte/smooth-muscle cells; Supplementary Fig. S19a). It also distinguished two spatially resolved remodelling programmes. The fibroblast-activation programme (*COL1A1, POSTN, COL3A1, FN1*) was lowest in remote myocardium, highest in the actively remodelling ischaemic zone and lower in the mature fibrotic scar (Fig. 6d). By contrast, the cardiomyocyte-stress programme (*NPPA, NPPB, ANKRD1*) concentrated in myocardium bordering the scar (Supplementary Fig. S19b). These zonal trends were visible in Visium spots. MINT localised each programme to the cells carrying it within a spot and sharpened boundaries that spot-level measurements average over (Supplementary Figs. S19c and S20). At an infarct border selected from the H&E image, *ACTN2* expression declined abruptly from surviving myocardium to the necrotic core. A straddling spot collapsed this transition into one value, whereas MINT resolved a sharp cell-level boundary (Fig. 6e; Supplementary Fig. S19d).

## 3 Discussion

Routine H&E histology has traditionally served as a qualitative diagnostic readout. MINT repositions it as a quantitative, sample-intrinsic measurement that complements molecularly rich but cellularly incomplete spatial-transcriptomic assays. We developed this strategy around the missing-coverage problem in Slide-tags, which directly profiles the whole transcriptome of individual recovered nuclei but loses most barcoded nuclei during recovery. Reconstruction from an adjacent H&E section requires joint cross-modal alignment and sparse-to-dense inference because the two sections contain independent nuclei in unequal numbers, lack cell-level pairs and differ through sectioning deformation. MINT combines sample-specific Slide-tags profiles with the dense cellular scaffold in histology to align the sections and infer expression for unsequenced histological nuclei. Its shared nucleus-centred representation also resolves pooled Visium measurements into cell-level profiles, extending the same complementary-measurement principle from missing nuclei to mixed capture features.

Existing reference-free approaches anchor predictions to fixed image tiles, isolated nuclei or expression propagated over a similarity graph [27–29]. MINT instead represents each nucleus together with its local niche. By reading expression from the section itself rather than an external reference, it can resolve cell states that the tissue contains but a reference atlas does not – as illustrated by rare lung lineages represented by fewer than 15 captured nuclei that became interpretable only after reconstruction (Fig. 4d). The image representation is shared, whereas task-specific heads match the molecular supervision: measured spot aggregates for Visium and spatially indexed nuclei linked by a soft cross-section transport plan for Slide-tags. This design resolves a mixed Visium spot into multiple cellular lineages and preserves informative cross-section correspondence when nuclear morphology alone is ambiguous.

The supporting evidence spans controlled and real settings. In pseudo-Slide-tags data with known ground truth, expression-guided alignment improved correspondence after nuclear loss and serial-section perturbation and increased recovery of genes excluded from alignment, and a separate human melanoma dataset reproduced both gains. Real Slide-tags reconstructions recovered organ architecture, marker organisation and cross-layer specificity across the olfactory bulb, kidney and brain; in clinical tissue, the lung maps resolved tertiary lymphoid structures and their maturity and the glioblastoma maps delineated cellular architecture and transcriptional programmes across sparse and partially uncovered sections. For Visium, Xenium-derived profiles withheld from spot-level training and a real Visium HD section supplied cell-level references, against which MINT resolved each measured spot profile across its morphologically distinct constituent cells.

MINT’s inference boundary is set by the information visible in histology. The set of cells represented in the output is defined by segmented nuclei, and predictions are most reliable for identities and programmes with morphological or microenvironmental correlates. Genomic alterations and transcriptional states that leave little histological trace still require direct sequencing or orthogonal assays. Accuracy also varies among genes and lineages; a single global confidence value would obscure this variation. Prospective serial-section studies with an independent, dense molecular reference will be needed to quantify reconstruction accuracy in routine clinical tissue.

The broader contribution of MINT is a sample-intrinsic formulation of cross-modal spatial inference: a dense, broadly available modality can align and constrain a sparse, information-rich one through optimal transport and conditional modelling, without one-to-one correspondence. Routine pathology thereby becomes a quantitative complementary measurement anchored by direct molecular data, and molecular breadth and cellular completeness are supplied by complementary readouts from the same tissue rather than optimised within a single assay. For Slide-tags, this adds only an adjacent serial H&E section and preserves the assay chemistry. The same scaffold extends to other paired measurements in which structural morphology complements a sparse molecular readout, such as spatial proteomics or chromatin accessibility, and provides a route for incorporating improved segmentation and pathology image models.

## 4 Methods

### 4.1 Image preprocessing

For Xenium data, the H&E image and spatial-transcriptomic coordinates are defined in different coordinate systems. We aligned the H&E image to the morphology coordinate system with the platform-provided 3 × 3 affine transformation matrix and bilinear interpolation. In Visium data, the H&E image and spot coordinates are intrinsically co-registered and required no additional alignment.

### 4.2 Nucleus segmentation

We segmented nuclei in the preprocessed H&E image with Cellpose [44], a generalisable cell-segmentation model. Cellpose inference yielded nuclear centroids and bounding boxes. For each nucleus *i*, we recorded its centroid (*x*_*i*_, *y*_*i*_), bounding box and area. CellViT++ [45] and StarDist are also supported as alternative segmentation backends.

We inspected segmentation overlays in the densest tissue regions. Segmentation recall and precision relative to Xenium DAPI-based nuclei are reported in Supplementary Fig. S1a–c.

### 4.3 Dual-scale morphological feature extraction

MINT extracts morphological features for each segmented nucleus at two spatial scales, capturing both cell-intrinsic and niche-dependent information.

#### Local features

For each nucleus *i* with centroid (*x*_*i*_, *y*_*i*_), we extracted a 224×224-pixel patch (~ 25 *µ*m at typical pathology resolution) centred on the nucleus from the stain-normalised H&E image. We encoded this patch with UNI2-h [46], the successor checkpoint to the UNI pathology foundation model [47], a pathology-specialised Vision Transformer (ViT-H/14) with 1,536-dimensional output. UNI2-h produces a 16×16 spatial-token map. We applied *spatial token aggregation*, averaging only tokens whose positions overlapped the nuclear bounding box within the patch:

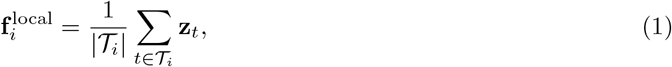

where *T*_*i*_ denotes the spatial tokens overlapping the bounding box of nucleus *i*, and **z**_*t*_ ∈ ℝ^1536^ denotes the token at position *t*. This procedure yields a nucleus-specific feature vector.

#### Global microenvironment features

We extracted a wider 1024 × 1024-pixel patch (~ 110 *µ*m) centred on the same nucleus to represent the surrounding tissue microenvironment. After resizing the patch to 512 × 512 pixels, we encoded it with DINOv2 [48] (ViT-L/14; 1,024-dimensional output). We applied the same spatial-token aggregation to the 36 × 36 token map after rescaling the nuclear bounding-box coordinates to the resized patch:

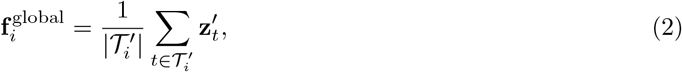

where 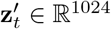.

#### Residual-gated fusion

We projected the local and global features to a common dimensionality *d* through learned linear layers: 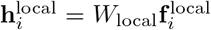 and 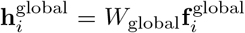. We then derived a joint backbone representation and a cell-specific gate from their concatenation:

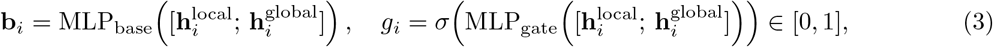

where *σ* is the sigmoid function and [·;·] denotes concatenation. The fused feature is then computed as a residual correction:

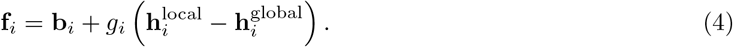

In the default implementation, *g*_*i*_ is a cell-specific scalar gate on the residual between the local and global projections and is learned end-to-end with the remaining network parameters.

Both foundation models are used in inference mode with frozen parameters. Features are standardised (zero mean, unit variance) per dimension across all nuclei in the sample before downstream prediction.

### 4.4 Disaggregation mode: resolving Visium spots into single-cell profiles

Visium measures transcripts captured across the tissue area under each spot; it does not isolate nuclei. In MINT disaggregation, each H&E-segmented nucleus serves as the spatial and morphological proxy for one cell-level output.

#### Spot–cell assignment

We assigned each nucleus-anchored cell to its nearest Visium spot when the nuclear centroid lay within the spot radius (*r* = 27.5 *µ*m). Cells whose nuclear centroids lay outside all spot boundaries were designated gap-region cells; they were excluded from training but retained during inference.

#### Model architecture

An hourglass multi-layer perceptron (MLP) maps the fused feature vector **f**_*i*_ ∈ ℝ^*d*^ to a predicted expression vector **ŷ**_*i*_ ∈ ℝ^*G*^, where *G* is the number of genes. Its layer widths are determined from the input and output dimensions. The encoder compresses the fused feature to a 512-dimensional bottleneck, and a decoder whose depth increases with *G* expands this representation before the output layer. Each hidden layer comprises a linear transformation, LayerNorm, LeakyReLU activation (negative slope 0.2) and depth-scaled dropout. The output layer uses softplus activation for log1p-normalised targets.

#### Spot aggregation loss

Because single-cell ground truth is unavailable, training uses a spot-level supervision strategy. For each Visium spot *s* containing *n*_*s*_ assigned cells, the predicted spot expression is the mean of its constituent cell predictions:

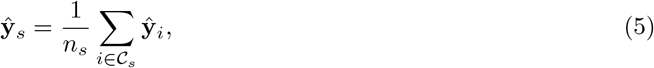

where *C*_*s*_ is the set of nucleus-anchored cells assigned to spot *s*. The spot-level loss is:

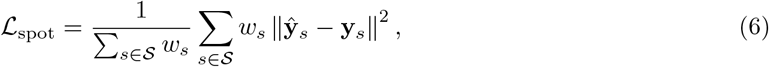

where **y**_*s*_ is the observed spot expression, S is the set of spots and *w*_*s*_ ∝ 1*/n*_*s*_ down-weights crowded spots. Thus, spots contribute comparably irrespective of the number of assigned cells; *w*_*s*_ = 1 gives the unweighted mean.

#### Per-gene correlation

To encourage each gene’s predicted spatial profile to track its measured profile beyond matching the spot mean, we add a per-gene Pearson correlation term at the spot level. For each gene *g*, the Pearson correlation *r*_*g*_ between the predicted and observed spot-level expression is computed across all spots, and the term penalises its complement:

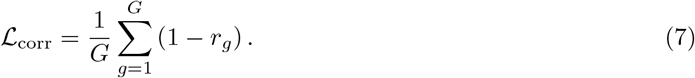

The total loss is ℒ = ℒ_spot_ + *λ*_corr_ ℒ_corr_. Because each prediction depends on the morphology of the corresponding cell, cells sharing a spot can receive distinct profiles despite spot-level supervision.

#### Training

We trained with spot-based mini-batches, AdamW optimisation, learning-rate warm-up and cosine annealing. All reported results used a single model per section.

### 4.5 Reconstruction mode: recovering dense expression from sparse Slide-tags data

#### Problem formulation

Let 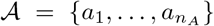 denote the nuclei segmented from the H&E section (section A, dense) and 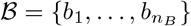 denote the nuclei with measured expression from the adjacent Slide-tags section (section B, sparse, *n*_*B*_ ≪ *n*_*A*_). The goal is to predict expression for all nuclei in *A* by jointly solving two coupled problems: establishing a soft spatial correspondence 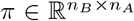 between sections, and learning a morphology-to-expression mapping *h*_*θ*_ : ℝ^*d*^ → ℝ^*G*^.

#### Fused Unbalanced Gromov–Wasserstein (FUGW) matching

The correspondence matrix *π* is obtained by solving a fused unbalanced Gromov–Wasserstein optimal transport problem [33–35]:

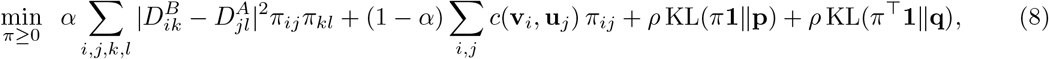

where *D*^*A*^ and *D*^*B*^ are intra-section pairwise distance matrices capturing spatial geometry, *c*(**v**_*i*_, **u**_*j*_) is a feature-space cost between the measured expression **v**_*i*_ of nucleus *b*_*i*_ and the predicted expression **u**_*j*_ = *h*_*θ*_(**f**_*j*_) of nucleus *a*_*j*_, *α* ∈ [0, 1] balances structural and feature terms, *ρ >* 0 controls marginal relaxation, and **p, q** are prior marginal distributions.

The Gromov–Wasserstein term (first sum) preserves pairwise distance structure without requiring the two point sets to share a common coordinate system. The unbalanced formulation (KL divergence penalties) accommodates the severe density imbalance caused by nuclear loss in the Slide-tags protocol.

#### FUGW configuration

We obtained the transport plan with an in-house FUGW solver in dense mode. The coarse-to-fine approximation was not used. Before optimisation, we isotropically normalised the coordinates of each section by subtracting the per-axis minimum and dividing by max(*x*-range, *y*-range), thereby preserving the *x*– *y* aspect ratio and mapping coordinates to the unit interval. The intra-section costs *D*^*A*^ and *D*^*B*^ were Euclidean distances in this normalised space. In fused rounds, *c*(**v**_*i*_, **u**_*j*_) was the cosine cost defined below (“Alignment feature from the generative model”); each cost matrix was scaled by its own maximum before combination through *α*. We used uniform marginal priors **p** and **q** and initialised the plan as their outer product. The marginal-relaxation parameter was asymmetric, *ρ* = (*ρ*_source_, *ρ*_target_) = (10.0, 0.1), anchoring sparse captured nuclei while permitting dense histological nuclei to remain unassigned. We used entropic regularisation *ε* = 1 × 10^−5^ and block-coordinate descent with 10 outer iterations of 500 Sinkhorn iterations each. For each transport-plan row, we retained the smallest set of highest-weight histological correspondences whose cumulative weight reached 99% of the row mass and renormalised the retained weights. The support size therefore adapted to the uncertainty of each captured nucleus.

Dense histological sections can contain up to ~ 1.8 × 10^5^ nuclei, for which a full target-side float32 distance matrix *D*^*A*^ would require ~ 135 GB. When *D*^*A*^ exceeded a fixed memory budget, we computed its rows from the normalised coordinates in blocks, using ∥*a* − *b*∥^2^ = ∥*a*∥^2^ +∥*b*∥^2^ −2 ⟨*a, b*⟩. We accumulated the (*D*^*A*^)^2^ and *D*^*B*^*π*(*D*^*A*^)^⊤^ terms blockwise and recomputed *D*^*A*^ as required; the smaller source matrix *D*^*B*^ (captured nuclei, ~ 10^4^ points) was retained in memory. The blockwise calculation reproduced the in-memory solution to floating-point precision and kept peak GPU memory near 20 GB, allowing the largest section to run on one 80 GB GPU.

#### Iterative Predict-then-Align strategy

MINT estimates correspondence by iterative predict-then-align refinement, which alternates optimal transport with representation learning [49]. At each round, the predictor is re-fitted to measured expression through the current correspondence, and the transport plan is re-solved against fixed tissue geometry. The refinement proceeds as follows:

- **Step 1 (spatial-only alignment):** We set *α* = 1 (pure spatial geometry) and solved equation (8) to obtain the spatial-only plan *π*_0_. This correspondence provided pseudo-labels for fine-tuning the morphology-conditioned generative model (“Alignment feature from the generative model”, below) on a compact spatially variable gene panel. We ranked the top 100 spatially variable genes in the Slide-tags data by Moran’s *I*, used the top 50 for alignment and reserved ranks 51–100 as an independent evaluation panel (Fig. 2d; “Pseudo-Slide-tags construction and evaluation”). Expression recovery was therefore never evaluated on a gene used for matching. The model’s conditional mean on the top-50 panel supplied the expression feature for the next alignment step.
- **Step 2 (expression-informed refinement**, *π*_0_ → *π*_1_ → *π*_2_): We set *α* = 0.8 to combine predicted-expression similarity with spatial geometry and solved equation (8) using conditional-mean predictions on the SVG panel as feature vectors **u**_*j*_, yielding *π*_1_. We re-fitted the model on this updated plan and repeated the refinement once to obtain *π*_2_, the final correspondence. Each predictor fit used the adaptive 99% cumulative-mass support defined above.

The 50-gene guidance panel comprises the most spatially structured genes in the data. In our experiments, enlarging this panel introduced weakly or noisily patterned genes into the optimal-transport feature cost and did not improve alignment. Real-data reconstructions used *π*_2_ as the final correspondence. The first refinement (*π*_0_ → *π*_1_) contributed most of the improvement, whereas the second (*π*_1_ → *π*_2_) added a smaller increment (Supplementary Fig. S2; Supplementary Table S3).

The guidance panel constrains the alignment but does not restrict reconstruction output. After fixing *π*_2_, we fitted a model of the same architecture with an output layer sized to the target panel, ranging from focused marker sets to the full measured transcriptome. The decoder therefore predicts every gene measured by the assay; whole-transcriptome Slide-tags data yield whole-transcriptome predictions, although alignment in every case uses the same 50-gene guidance panel.

#### Alignment feature from the generative model

The morphology-conditioned generative model supplies the expression feature for the fused alignment step (equation (8)). During alignment iterations the model used only the local (224-pixel) morphological feature, which is cheap to recompute each round and suffices to predict the compact guidance panel used for matching; the contextual scale is added only for the final reconstruction (“Generative reconstruction model”, below). At each round, we re-fitted the model on the current plan’s *π*-pooled captured counts using the compact spatially variable gene panel. Its conditional mean, **u**_*j*_ = E[**x** | **f**_*j*_], was estimated by averaging a small number of flow samples and entered the feature cost as *c*(**v**_*i*_, **u**_*j*_) = 1 − cos(**v**_*i*_, **u**_*j*_) in log(1+*x*), *L*_2_-normalised space. Because the features were *L*_2_-normalised before optimisation, the squared-Euclidean feature distance used by transport was equivalent to this cosine cost. Averaging flow samples approximated the conditional mean, stabilising the feature cost across rounds while keeping transport optimisation tractable. Where morphology carried little expression signal, spatial geometry contributed most of the fused correspondence.

#### Generative reconstruction model

After fixing *π*_2_, the final reconstruction model — now conditioned on the fused (dual-scale) morphological feature rather than the local feature used during alignment — generates the dense single-cell map by sampling from the conditional expression distribution rather than using its mean. Each Slide-tags profile is an independently measured whole-transcriptome profile of one nucleus, so the data directly observe cell-to-cell expression variability. MINT therefore models *p*(counts | morphology) and samples a distinct profile for each nucleus, preserving this variability rather than collapsing morphologically similar nuclei onto a shared mean. The cross-section correspondence *π* is a soft, uncertain many-to-one assignment, and a sampled profile represents each nucleus as one plausible draw from its conditional distribution. By contrast, Visium pooling destroys single-cell information – no individual cell profile is observed, and supervision can constrain only the aggregate mean within each spot – so disaggregation uses deterministic prediction of the conditional mean (Results).

We optimised the model end-to-end on *π*-pooled soft pairs. A morphology encoder mapped the fused feature **f**_*j*_ of nucleus *a*_*j*_ to a conditioning embedding. A variational encoder with a negative-binomial (NB) decoder defined a cell-state latent space, and a conditional flow-matching (rectified-flow) network learned to transport a standard Gaussian **z**_0_ ~ N(**0, I**) to this distribution. The NB decoder mapped the latent and morphology to a per-gene rate **r**_*j*_ (a softmax distribution over genes) with per-gene dispersion. Each captured nucleus *b*_*i*_ (integer counts **y**_*i*_, library size *L*_*i*_) was paired with the row-specific set of histological nuclei *a*_*j*_ retained by the 99% cumulative-mass rule. Pair (*b*_*i*_, *a*_*j*_) contributed the renormalised transport-weighted negative-binomial likelihood *π*_*ij*_ NB-NLL(**y**_*i*_|*L*_*i*_ **r**_*j*_). The full objective additionally included the flow-matching velocity-regression loss and a *β*-VAE KL term. The same transport-weighted supervision was used during alignment and final reconstruction; final outputs were sampled rather than point estimates.

At inference, each nucleus *a*_*j*_ in section A is represented by its morphology. We sampled a latent from the prior, integrated it through the conditioned flow, converted it to a per-gene rate with the NB decoder, scaled the rate by a per-nucleus library size and sampled integer counts from the NB distribution. This yielded a dense AnnData object of single-cell counts, subsequently CP10k+log1p-normalised for downstream analysis. The output panel matched the measured gene set and was independent of the compact guidance panel. The final reconstruction flow used AdamW optimisation with linear warm-up, cosine annealing, gradient clipping and an exponential moving average of parameters (decay 0.999); shorter alignment-round fits used AdamW with cosine annealing. Reconstruction fidelity and diversity relative to the reference distribution are reported in Supplementary Table S5.

### 4.6 Gene expression normalisation

Unless otherwise stated, we normalised expression values to 10,000 total counts per cell or spot (CP10k) and applied a log(1+*x*) transform (CP10k+log1p). For disaggregation, we computed the spot-aggregation loss (equation (6)) in CP10k+log1p space. Some MINT configurations predict per-gene min–max-scaled values for numerical stability. Before comparison with measured data, we denormalised these outputs to raw counts with the stored per-gene scaling parameters, clipped values at zero and re-normalised them to CP10k+log1p space. For downstream clustering, we re-normalised MINT predictions with `sc.pp.normalize_total` followed by log1p before highly variable gene selection. Cross-sample analyses used absolute CP10k+log1p units rather than per-sample *z*-scores.

### 4.7 Benchmark datasets and evaluation metrics

#### Pseudo-Visium construction

To evaluate disaggregation mode in a controlled setting where single-cell ground truth is available, we simulated pseudo-Visium data from Xenium datasets. Xenium provides H&E images and spatially resolved transcript coordinates at subcellular resolution, enabling construction of a ground truth that can be directly compared to MINT’s cell-level predictions. For each section, every H&E-segmented nucleus served as the anchor for one evaluation cell. Xenium transcripts were assigned to their nearest cell anchor, equivalent to a Voronoi partition of the tissue by nuclear centroid, and aggregated to form a Xenium-derived expression profile for each cell. For each Xenium section, we imposed a regular hexagonal grid with 55-*µ*m spot diameter (matching Visium geometry) and aggregated these cell-level profiles within each spot to generate simulated Visium spot measurements. MINT was then trained on these pseudo-Visium spot measurements using the paired H&E image, and its single-cell predictions were scored against the Xenium-derived ground truth for all segmented cells (Cell-level evaluation, below). For the breast cancer sections, of the 541-probe Xenium panel listed in Supplementary Table S1, 307 (Rep1) and 305 (Rep2) gene targets were retained after per-replicate quality-control filtering and used for training and evaluation; the other tissues used their full panels. The two breast cancer replicates were Reinhard colour-normalised [50] to a shared reference before feature extraction, harmonising the staining between the two sections (Supplementary Fig. S1d).

#### Spot-level evaluation

For secondary spot-level evaluation, we averaged MINT’s per-cell predictions within each spot boundary and compared the aggregates with pseudo-Visium spot ground truth. For each gene *g*, we report:

- **Per-gene Pearson correlation coefficient (PCC):** Pearson *r* between predicted and observed spot expression vectors across all spots.
- **Per-gene RMSE:** 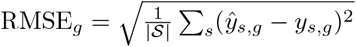, computed on [0, 1]-normalised expression.
- **Per-gene SSIM:** Structural similarity index measure between the predicted and observed spatial expression maps, treating each gene’s spot-level expression as a spatial image (consistent with iStar [27] and Thor [28]).

Metrics are summarised as mean ± standard deviation across all genes, and as hit rate (fraction of genes with PCC *>* 0.3 and PCC *>* 0.5).

#### Cell-level evaluation

The primary comparison was cell-for-cell on the shared evaluation cells represented by H&E-segmented nuclei. This is an in-sample, transductive evaluation because cells contributing to pooled training spots were also evaluated; cross-slice transfer to an independent breast-cancer replicate assessed out-of-sample generalisation separately. The predictor receives only each cell’s nucleus-centred morphological features, without spatial coordinates. Because all methods were evaluated on the same nucleus-anchored cells, predicted cell *i* corresponded directly to its Xenium-derived ground-truth profile, without spatial matching; values were compared in a common CP10k+log1p space. Segmentation recall relative to Xenium DAPI nuclei is reported separately below. The comparator set comprised reference-free single-cell predictors. The Visium HD analysis additionally included a non-learning morphology-only nearest-neighbour baseline. Reference-based deconvolution methods, including cell2location[18] and Tangram[19], estimate per-spot cell-type proportions from matched scRNA-seq references and do not produce cell-level expression profiles for direct comparison here.

Four metrics are computed at the cell level: (1) **Per-gene Pearson** *r* **and Spearman** *ρ*: correlation between predicted and Xenium-measured expression across all cells for each gene; (2) **Per-gene normalised RMSE:** RMSE computed after min–max normalising each gene’s expression to [0, 1] using the Xenium ground-truth range, removing the influence of gene expression magnitude; (3) **Per-gene spatial SSIM:** the predicted and measured single-cell expression values are rasterised onto a 96 × 96 regular spatial grid by averaging the cells that fall within each bin (empty bins set to the neutral background value), and SSIM is computed between the resulting spatial maps, capturing whether the predicted spatial expression pattern preserves fine-grained tissue structure; (4) **Cell-wise profile Pearson** *r*: for each individual cell, the Pearson correlation between its predicted and Xenium-measured expression vectors across all panel genes, reflecting single-cell transcriptional profile reconstruction quality. This cell-level evaluation framework is analogous to the approach used by sCellST [29] and Thor [28].

#### Pseudo-Slide-tags construction and evaluation

To validate reconstruction mode with known spatial correspondence, we constructed pseudo-Slide-tags data from Xenium Prime Mouse Brain Coronal (50,648 nuclei, 5,006 genes) and an independent Xenium Human Skin Melanoma section (82,696 nuclei, 282 genes after removing negative-control probes). The H&E-segmented nuclei passing quality control formed the complete reconstruction target set *A*_model_. Its subset *A*_ref_ with a uniquely assigned Xenium profile formed the reference set: each Xenium transcript was assigned to its nearest H&E nucleus, and retained nuclei inherited that aggregated profile and a fixed true A id. Pseudo-Slide-tags input *B* was drawn uniformly without replacement from *A*_ref_; its nuclei retained their profiles and identities while only their coordinates were perturbed. Nuclear loss therefore denotes deletion from *A*_ref_, not loss from the reconstruction target set. Within each realisation, a single random permutation defined nested retained sets across loss rates.

The mouse-brain primary benchmark used 30, 50, 70 and 90% nuclear loss. All perturbations were defined in physical coordinates: Gaussian jitter independently added *N* (0, *σ*^2^) to each axis, with *σ* = 0, 2, 5, 10, 20 or 40 *µ*m. For thin-plate-spline (TPS) deformation, a 5 × 5 control-point grid extended 10% beyond the unperturbed *B* bounding box. Mean-centred, two-dimensional Gaussian control-point displacements were interpolated separately by coordinate with a strictly interpolating TPS and rescaled so that the median displacement across *B* nuclei was 0, 2, 5, 10, 20 or 40 *µ*m. We recorded the realized displacement distribution and rejected fields with a non-positive Jacobian determinant. The skin melanoma validation used the same construction and analysis with its separate numerical conditions, reported in Supplementary Fig. S2 and Supplementary Table S3.

For each dropout realisation, we removed control probes and genes detected in fewer than max(50, 0.05*n*_*B*_) nuclei, then ranked the remaining genes by Moran’s *I* in the observed *B* coordinates. Moran’s *I* used CP10k+log1p expression, a symmetric *k* = 15 nearest-neighbour graph, and row-standardised binary weights. The top 100 genes were selected before perturbation and reused across its jitter and TPS conditions. Ranks 1–50 provided the expression feature for FUGW alignment; ranks 51– 100 were held out from alignment to evaluate whether the transport plan generalised to spatial patterns it had not used. They were not held out from final reconstruction training: after the transport plan was fixed, the generative model was trained to output the complete non-control gene panel, while the alignment evaluation reported the ranks 51–100 panel. We used five independent data realisations and three training seeds within each realisation; training-seed results were averaged within realisation before summary across the five realisations.

Each retained *B* nucleus was evaluated against its fixed true A id. Top-1 distance was the Euclidean distance, in the original *A* physical coordinate system, between the argmax entry in its transport-plan row and its true counterpart. Mean rank was the 1-based descending rank of the true counterpart in that row, with a fixed stable ordering for ties. Top-*k* matching accuracy was retained as a supplementary metric. The GT reference used the identity transport plan and therefore isolates reconstruction error from alignment error. For the manual baseline, three independent annotators, blind to the true nuclear identities and method results, each placed the same 15 pre-specified anatomical landmarks spanning the tissue edge and interior. Each affine transform mapped *B* to *A* before nearest-nucleus hard matching; the reported manual-baseline metrics are the mean across annotators.

#### Segmentation coverage of the evaluation cell set

H&E-based and Xenium DAPI-based segmentation overlapped only partially. At a 10-*µ*m matching threshold, Cellpose achieved 74.7% recall against Xenium nuclei (precision, 96.4%; F1, 0.84), leaving approximately 25% of Xenium-detected nuclei without a corresponding H&E-segmented nucleus. These unmatched nuclei contributed no cell-level prediction and were excluded from the shared evaluation cell set. All compared H&E-to-spatial-transcriptomics methods used this same set of segmented nuclei [28, 29]. The reported cell-level metrics therefore apply to the matched cell set and do not quantify performance across all Xenium-detected nuclei.

#### Spatial clustering comparison

To qualitatively compare nucleus-centric and grid-based feature extraction strategies, we performed unsupervised spatial clustering on morphological features extracted by three approaches: MINT local (UNI2-h, 224-pixel nucleus-centric patches, 1536 dimensions), MINT global (DINOv2, 1024-pixel nucleus-centric patches, 1024 dimensions), and grid-based tiling (ViT, fixed-size tiles, 576 dimensions). For the grid-based approach, features were extracted at the tile level and each cell was assigned the feature vector of the tile containing its centroid, following the tiling strategy commonly used in computational pathology pipelines. All three feature sets were processed through an identical unsupervised clustering pipeline: PCA reduction to 50 components, construction of a *k*-nearest-neighbours graph (*k* = 15), and Leiden clustering. This comparison was performed on the Xenium FFPE Human Breast Cancer dataset (129,440 cells; Leiden resolution 0.5) and an in-house human lung cancer tissue section (S16; 49,499 cells; Leiden resolution 1.0).

### 4.8 Real Visium datasets and disaggregation on real data

We applied MINT disaggregation to five real Visium datasets: mouse olfactory bulb and coronal brain (10x Genomics), human breast ductal carcinoma in situ (10x Genomics), human myocardial infarction tissue spanning remote, ischaemic and fibrotic zones (Kuppe et al.[43]), and human colon adenocarcinoma profiled by Visium HD (10x Genomics). We used five of the six available myocardial-infarction sections; one was excluded because of poor H&E quality. For every dataset, we segmented nuclei from the full-resolution H&E image and extracted dual-scale morphological features as in Sections 4.2–4.3. For standard-resolution Visium, the morphology-to-expression MLP was trained with measured spots under spot-level supervision (equations (5)–(7)). As these data lacked cell-level ground truth, we evaluated predictions using marker-gene spatial patterns, the Allen Brain Atlas for neural tissues and pathological features for tumour tissues. The colon Visium HD section provided a near-single-cell reference for direct scoring (Section 4.9).

#### Training configuration

We trained spot-supervised predictors for 600 epochs at a learning rate of 3 × 10^−4^ without early stopping. An initial 200-epoch schedule at 1 × 10^−4^ did not recover sharply localised genes, including myelin genes; held-out spot validation supported the longer schedule. We disabled the gate-entropy regulariser for all real-data runs because it drove the fusion gate towards a uniform value and over-smoothed predictions. For held-out validation, we withheld a subset of spots entirely from supervision and compared spot-aggregated predictions with their measured expression.

#### Clustering and annotation

We clustered single-cell predictions with BANKSY, which augments each cell’s expression with a neighbourhood-averaged feature before graph construction and Leiden clustering. We annotated clusters by marker-group dominance: a background-corrected module score was computed for each candidate cell type, and clusters were assigned to the type with the dominant score. We annotated anatomical layers and regions in olfactory bulb and brain, and cell types in breast and colon tumours. MINT and spot annotations were generated at their respective resolutions and compared by biological interpretation rather than forced label concordance. We reported cell-type reliability as the per-gene correlation between MINT and spot expression for lineage markers. Nuclear-density and nuclear-count features were not used for biological claims because segmentation under-detects nuclei in dense immune and fibrotic regions. A DCIS Visium spot was considered to span multiple cell types when MINT-predicted cells within its 55-*µ*m footprint contained at least two labels.

### 4.9 Visium HD single-cell construction and cell-level evaluation

To obtain a near-single-cell reference on a real sequencing platform, we used the continuous 2-*µ*m bin grid of Visium HD and treated each H&E-segmented nucleus as the anchor for one cell. Bin centres were converted to the physical coordinate system of the nuclear masks. A bin centre within a mask was assigned to the corresponding cell; otherwise, it was assigned to the cell with the nearest mask boundary only when the distance was ≤8 *µ*m. When masks overlapped or distances tied, the bin was assigned to the cell with the nearest nuclear centroid. Each bin was assigned once or discarded. Cell anchors were retained when they received at least five uniquely assigned bins and 20 aggregated UMIs, lay within the tissue mask without image or tissue-crop truncation, and passed segmentation-artifact and confidence quality control. For the colon adenocarcinoma section this yielded 59,964 reference cell profiles from 108,506 segmented nuclear anchors (median 41 bins and ~ 400 unique molecular identifiers per cell).

To provide the spot-level input, we placed 55-*µ*m-diameter circular capture areas on a non-rotated hexagonal lattice with a 100-*µ*m centre-to-centre pitch. The lattice origin was fixed in the uncropped slide coordinate system; the four prespecified offsets were (0, 0), (50, 0), (25, 43.301) and (75, 43.301) *µ*m. We retained spots whose capture circle intersected the tissue and assigned a bin to a spot only when its centre lay within the 27.5-*µ*m capture radius. Each offset therefore defined an independent pseudo-Visium input, with raw UMI counts aggregated within spots after removing control probes and normalised per spot to CP10k+log1p.

MINT was trained independently for each grid under the same spot-level disaggregation supervision used for standard Visium (spot-mean MSE and per-gene Pearson, equations (5)–(7)). The morphology-only nearest-neighbour baseline used the same frozen dual-scale H&E features: each spot was represented by the mean feature of its constituent cell anchors, and each target cell received the linear CP10k expression profile of its nearest spot prototype. The global-median baseline assigned every target cell the per-gene median of the training spots’ linear CP10k expression. All three methods predicted the same nucleus-anchored cells, and the HD-derived cell profiles were used only for scoring. Before comparison, predictions and reference profiles were converted to a common per-cell CP10k+log1p space. We evaluated the shared reference cells and the common non-control gene set detected in at least 5% of reference cells; genes with zero reference variance were excluded from correlation calculations. Single-cell accuracy was quantified by per-gene Pearson correlation between predicted and measured expression, stratified by gene detection rate. Each grid was scored independently, and reported summaries are the mean, standard deviation and 95% confidence interval across the four grids.

### 4.10 Real Slide-tags datasets and reconstruction on real data

We applied reconstruction mode to Slide-tags datasets generated for this study from mouse olfactory bulb, kidney and brain. Slide-tags libraries were generated with the published protocol [8]. For each dataset, we acquired H&E images from the immediately preceding and following serial sections of the same tissue block. We included one H&E section per block that met prespecified quality criteria: tissue integrity, staining and focus quality, absence of sectioning artefacts, and overlap with the Slide-tags tissue area. Captured and segmented nuclei were independent populations and therefore had no one-to-one correspondence. The selected H&E section provided the dense cellular scaffold for the sparse whole-transcriptome measurement. We segmented nuclei with CellViT++[45] and extracted morphological features as described above. We established correspondence with the iterative FUGW Predict-then-Align procedure. Because FUGW matches internal pairwise-distance geometry, no landmark-based registration was used; maps were displayed in a common orientation. After removing off-tissue captured nuclei, we obtained a spatial-only correspondence and refined it with expression information. Alignment used the generative model’s conditional mean on the 50-gene SVG panel, and the generative model sampled the dense whole-transcriptome reconstruction using captured counts as the *π*-pooled supervision target. Pseudo-Slide-tags benchmarks provided direct tests against unsequenced nuclei with known correspondence. We assessed real reconstructions by anatomical structure, marker co-expression and the change in marker-gene Moran’s *I* from sparse to dense maps. As a spatial-smearing control, we tested whether markers absent from an anatomical layer in sparse data remained absent after reconstruction.

### 4.11 Clinical lung adenocarcinoma analysis

We analysed five human lung adenocarcinoma resections (adenocarcinoma in situ and minimally invasive adenocarcinoma) and one non-malignant granuloma, each profiled by Slide-tags with an adjacent-section H&E image. We segmented nuclei with CellViT++ followed by manual curation and extracted morphological features as described above. The generative model predicted every measured gene, yielding whole-transcriptome reconstructions. Cross-sample quantities were calculated in absolute CP10k+log1p units. Markers undetected in a sample were predicted near zero and classified as not measured rather than biologically absent. We established correspondence with the iterative FUGW procedure used for the mouse Slide-tags data and reconstructed dense single-cell maps with the morphology-conditioned generative model. We annotated cells by absolute marker dominance. For each cell type, the mean expression of its markers (Supplementary Table S7) in absolute CP10k+log1p units defined an independent per-type score. We applied an immune gate, resolved immune cells into B-follicle, follicular dendritic cell, T-cell, myeloid and plasma identities, and assigned non-immune cells to epithelial, stromal or vascular compartments. Genes detected in fewer than 1% of captured nuclei were excluded from downstream analysis. Each follicle call was accompanied by the captured-nucleus detection rate of its defining FDC markers (*CR2, FCER2*); undetected markers were treated as not measured. Two spatial statistics summarised each B-cell follicle. Focality was the fraction of the top-decile B-lineage cells, defined by the compact B-lineage score (*MS4A1, CD79A, PAX5*; Supplementary Table S7), lying within twice the follicle radius *R* (6% of the section span) of the single densest B focus. FDC fold-enrichment was the mean FDC-lineage signal (*CR2, FCER2*) among the top-decile B-lineage cells divided by the tissue-wide mean FDC signal (4.4-fold in the exemplar TLS, P2). A cell was counted positive for a lineage when its score exceeded 0.5 in absolute CP10k+log1p units, and the pan-immune programme map (Extended Data Fig. 3c) is the *PTPRC* field. The tertiary-lymphoid-structure signature scores were computed from the gene sets in Supplementary Table S7: the breast-panel signature (Fig. 5d) with score genes, and the lung B-follicle/germinal-centre signature (Extended Data Fig. 3b) by per-gene min–max scaling to [0, 1] followed by a cross-gene mean. TLS maturity was assessed from spatial organisation, including focal FDC networks and concentric follicle architecture, together with captured-nucleus detection of FDC markers. All clinical analyses were observational and used one section per patient. The two immune-cold cases carried oncogenic driver mutations (P4, *EGFR*; P5, *ERBB2*). Less-inflamed microenvironments have been reported in EGFR- and ERBB2-mutant lung adenocarcinoma [38, 39], but the association is not uniform and no statistical inference was drawn across the six patients.

### 4.12 Clinical glioblastoma analysis

We analysed five Slide-tags samples from three glioblastoma patients: two patients contributed tumour-core and peritumoural-margin samples (GL1C, GL1P, GL2C and GL2P), and one contributed a tumour-core sample (GL3C). Each sample had an adjacent-section H&E image. In GL3C, Slide-tags covered approximately 40% of the section, and MINT reconstructed the full section from adjacent histology. We segmented nuclei with CellViT++ and extracted morphological features as described above. The generative model predicted every measured gene, and cross-sample quantities were calculated in absolute CP10k+log1p units. We established correspondence with iterative FUGW alignment and generated dense maps with the morphology-conditioned generative model, as for the lung cohort. Malignant transcriptional programmes were scored with background-corrected module scores over the Neftel cell-state signatures (MES-like, OPC-like, NPC-like, AC-like) [40] and the Verhaak subtype signatures [41]; microenvironmental compartments (myeloid/macrophage, vascular) and a perinecrotic hypoxia programme were scored from curated marker sets. Because glioblastoma forms a continuous transcriptional spectrum, programme scores are reported as regional axes rather than discrete cell-type calls; the mesenchymal programme in particular lacks cell-type-specific markers. All glioblastoma analyses were observational and used one section per patient.

### 4.13 Statistical analysis

We assessed paired per-gene accuracy differences with two-sided Wilcoxon signed-rank tests across genes. Because genes are strongly co-expressed and are not statistically independent, these tests were interpreted descriptively; the fraction of genes for which MINT performed better was treated as the primary comparison. Extended Data Fig. 7b additionally reports paired two-sided Wilcoxon signed-rank tests across cells for per-cell profile correlations; these tests were descriptive because cells from the same section were not independent. We report Pearson *r* or Spearman *ρ* as indicated. Per-gene Pearson *r* was the primary accuracy metric; cell-wise profile correlation, Spearman *ρ*, RMSE and SSIM were reported as complementary metrics. We controlled gene-level *p*-values by the Benjamini–Hochberg procedure when multiple genes were tested individually. No correction was applied to a single global test. Sample sizes are reported with each analysis. The clinical cohorts were observational convenience case series, with sample size determined by specimen availability; no power calculation was performed. Clinical analyses used one section per patient and did not support across-patient statistical inference.

### 4.14 Training details

The Visium disaggregation predictor and Slide-tags reconstruction model were trained separately for each section using that section’s spatial measurement, without an external reference. Both used AdamW with random seed 42. The disaggregation predictor was an MLP over fused morphological features with an automatically sized hourglass architecture: a 512-dimensional bottleneck, a decoder whose depth increased with target-gene number and depth-scaled dropout (~0.2–0.25). We trained it for 600 epochs at a learning rate of 3 × 10^−4^, batch size 1,024 and weight decay 10^−3^, with a 10-epoch cosine warm-up and cosine schedule and without early stopping. The loss combined a spot-mean term (*λ*_mse_ = 1.0) and per-gene Pearson term (*λ*_pearson_ = 0.2). We set the residual-gate entropy regulariser to zero. The reconstruction model used conditional flow matching with a negative-binomial count decoder. We re-fitted the model at each alignment round on the SVG panel and once on frozen *π*_2_ over the full gene panel, using *π*-pooled captured-nucleus supervision, a learning rate of 10^−3^ and weight decay 10^−5^. Its objective comprised negative-binomial negative log-likelihood, flow-matching velocity loss and a *β*-VAE KL term (*β* = 0.5). Default hyperparameters were shared across tissues and not tuned per sample.

### 4.15 Software and hardware

MINT was implemented in Python with PyTorch for model training and Scanpy/AnnData for single-cell data handling. The paper-associated release pins PyTorch 2.2.0, Scanpy 1.10.3 and AnnData 0.9.2. We extracted morphological features with UNI2-h[46, 47] and DINOv2[48], segmented nuclei with Cellpose[44] or CellViT++[45], performed optimal-transport alignment with a fused unbalanced Gromov–Wasserstein solver and used BANKSY for spatially aware clustering[42]. Foundation-model inference and training were performed on NVIDIA A800 GPUs (80 GB). Each section-level predictor used one GPU, whereas the two feature-extraction scales ran in parallel on separate GPUs. Training required ~ 30–40 min for a moderate-sized Visium section (for example, 37 min for breast DCIS with 51,680 cells and 2,045 genes); inference on a new H&E section from a trained tissue completed within minutes. We ran iStar, Thor and sCellST from their public repositories with default settings, modifying only input-format handling, and used identical hardware across methods. Thor completed on the four smaller sections but exceeded a 200 GB host-memory budget on the largest section (human renal carcinoma, ~ 388,000 cells) during cell-similarity-graph diffusion.

## Acknowledgements

This work was supported by the Prevention and Control of Emerging and Major Infectious Diseases–National Science and Technology Major Project (project no. 2027ZD01999500; subproject no. 2027ZD01999501), the Major Project of Guangzhou National Laboratory (grant nos. GZNL2024A01015 and GZNL2024A03001), and the National Natural Science Foundation of China (grant no. 32401247).

## Author contributions

L.C. and L.T. conceived the study. L.C. developed the methodology and software, performed the analyses, prepared the figures and wrote the original draft. L.T. contributed to methodology, supervision, funding acquisition and manuscript revision. Z.Z. provided the in-house lung cancer Slide-tags data and contributed to its biological interpretation. W.R. and S.L. established and optimised the Slide-tags workflow and generated the lung cancer data. F.Y. and Q.Z. provided the glioblastoma specimens, and H.P. coordinated Slide-tags sequencing and curation of the glioblastoma data. B.W., M.X., Y.Z. and X.L. generated the mouse and glioblastoma Slide-tags data and performed upstream processing. T.Y. contributed to algorithm development and software organisation. All authors reviewed and approved the manuscript.

## Data availability

The Xenium datasets used for the pseudo-Visium and pseudo-Slide-tags benchmarks (FFPE Human Breast Cancer, Human Renal Carcinoma, Human Liver tumour and non-diseased sections, Prime Mouse Brain Coronal and Human Skin Melanoma) are publicly available from 10x Genomics; accession URLs for every dataset are listed in Supplementary Table S1. The HER2ST dataset used for colour normalisation validation was obtained from Andersson et al. [51] and is available at Zenodo (https://doi.org/10.5281/zenodo.4751624). The real Visium datasets (mouse olfactory bulb, mouse coronal brain, human breast ductal carcinoma in situ and human colon cancer Visium HD) are publicly available from 10x Genomics (accession URLs in Supplementary Table S1), and the human myocardial infarction Visium data are from Kuppe et al. [43] (Zenodo, https://doi.org/10.5281/zenodo.6580069). Inhouse mouse Slide-tags data (olfactory bulb, kidney and brain), each paired with an adjacent-section H&E image, have been deposited in the National Genomics Data Center (NGDC), China National Center for Bioinformation, under BioProject PRJCA069611 and OMIX accession OMIX018761, and will be released publicly upon publication. The human lung Slide-tags data analysed here are part of a companion study and have been deposited in the National Genomics Data Center (NGDC), China National Center for Bioinformation, under BioProject PRJCA068416 and OMIX accession OMIX018434, comprising Slide-tags spatial transcriptomic data from 25 tissue sections together with matched chromatin-accessibility data from four samples. From this deposit, the present study analysed six lung sections that passed quality control and were each paired with an adjacent-section H&E image: sections S16 (P1), S7 (P2), S13 (P3), S10 (P4), S14 (P5) and S12 (P6) in Supplementary Table S4. The human glioblastoma Slide-tags data are part of a separate study in preparation and are available from the corresponding author on reasonable request; they will be deposited under controlled access upon publication of that study. All data supporting the findings are available as described above.

## Code availability

During peer review, a versioned archive of the MINT source code, documentation and demonstration materials is provided confidentially to the editors and referees under the PolyForm Noncommercial License 1.0.0. The paper-associated release will be made publicly available in a dedicated GitHub repository and archived in a DOI-minting repository no later than publication. The repository URL and archive DOI will be added when available.

## Use of large language models

During the preparation of this work the authors used large language models (Anthropic Claude and OpenAI GPT) to assist with code development, data analysis and manuscript editing. The authors reviewed and edited all generated content and take full responsibility for the integrity and accuracy of the work.

## Ethics statement

The lung adenocarcinoma resections and the non-malignant pulmonary granuloma analysed in this study were collected under a protocol approved by the Ethics Committee of the First Affiliated Hospital of Nanjing Medical University (Jiangsu Province Hospital) (approval no. 2025-SR-458), with written informed consent obtained from all participants. The glioblastoma specimens were collected under a protocol approved by the Human Research Ethics Committee of the Second Affiliated Hospital of Zhejiang University School of Medicine (approval no. IRB-2026-2161), with written informed consent obtained from all participants. All animal procedures were approved by the Animal Care and Use Committee of Westlake University (license no. AP#23-111-LXD). Olfactory bulb, kidney and brain tissues were dissected from the same batch of 8-week-old male C57BL/6J mice, group-housed on a 12-h light–dark cycle, for Slide-tags profiling. Public datasets analysed here (10x Genomics Xenium and Visium, Kuppe et al. human myocardial infarction Visium) were released under the ethics approvals of their original studies.

---

**Extended Data Fig. 1.**
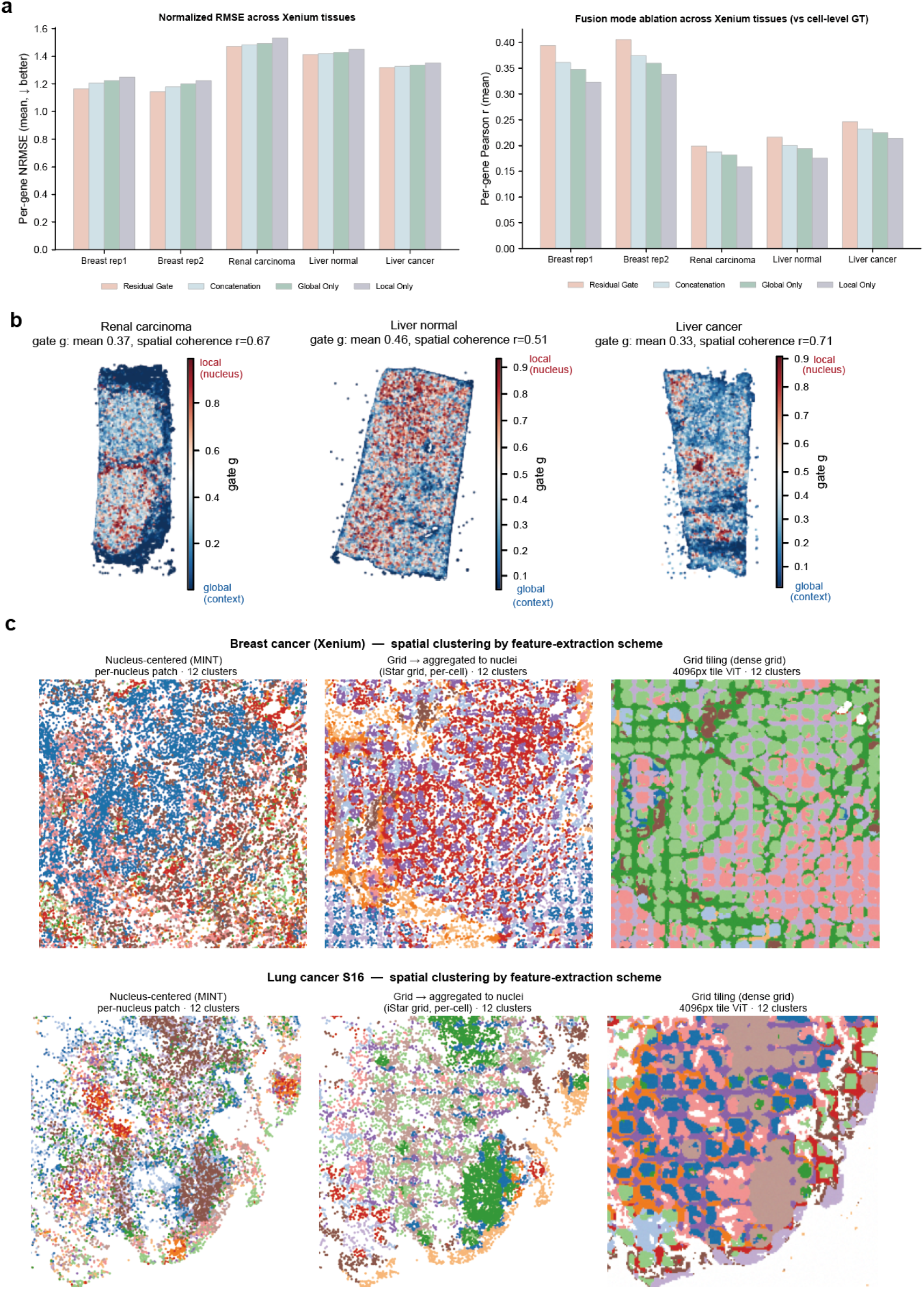
Ablation of dual-scale feature extraction: residual-gated fusion and nucleus-centred versus grid-based features. **a**, Prediction accuracy of residual-gated fusion, concatenation, global-only and local-only feature extraction across five Xenium tissues (breast Rep1, breast Rep2, renal carcinoma, normal liver and liver cancer): mean per-gene Pearson correlation (left) and mean per-gene normalised RMSE (right), evaluated against cell-level ground truth. Residual-gated fusion ranked first or tied first in each tissue. **b**, Spatial maps of the learned residual gate *g* in representative tissues. A value of 0 leaves the joint local–global backbone unchanged; increasing *g* adds a larger local-minus-global residual. Per-tissue spatial coherence is the correlation between each cell’s gate and the mean gate of its 15 nearest spatial neighbours (*r* = 0.51–0.73). **c**, Spatial clustering with nucleus-centred features, grid features aggregated to nuclei and dense grid tiling in Xenium breast cancer and lung cancer section S16. Rectangular tile-boundary patterns are visible in the dense-grid maps.

**Extended Data Fig. 2.**
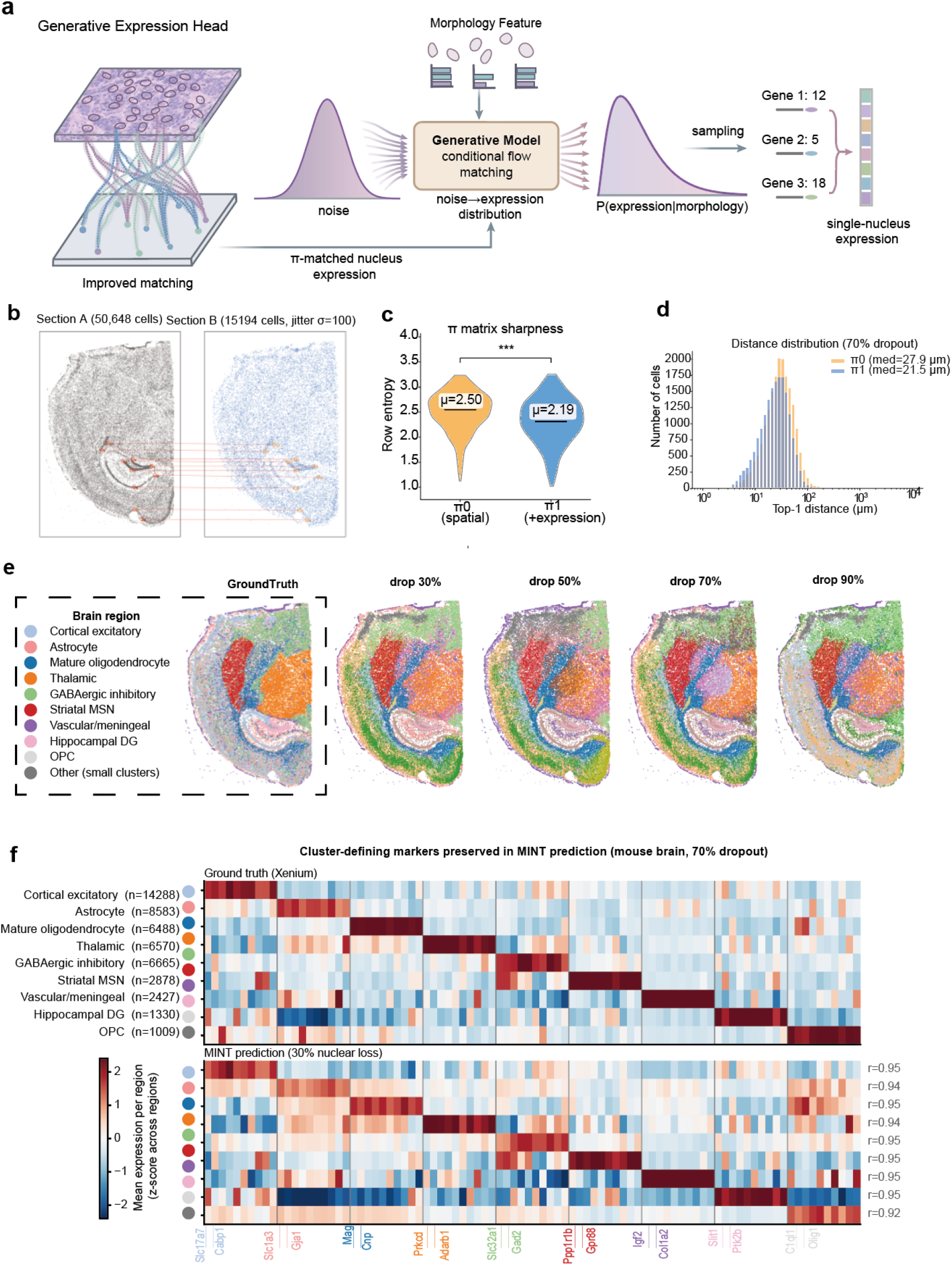
Generative reconstruction and alignment validation on Xenium mouse-brain pseudo-Slide-tags data. **a**, Generative expression head. The fused morphological feature of each nucleus and expression from its *π*-matched nuclei condition a flow-matching model that transports noise to a conditional expression distribution; sampling from this distribution yields a discrete single-nucleus profile. **b**, Dense reference section A (50,648 nuclei) and sparse section B (15,194 nuclei after 70% nuclear loss and Gaussian jitter, *σ* = 100 *µ*m). Example transport links illustrate the cross-section correspondence. **c**, Sharpness of the FUGW transport plan, quantified as row entropy. The expression-guided plan (*π*_1_, mean entropy 2.19) is sharper than the spatial-only plan (*π*_0_, mean entropy 2.50; ∗∗∗, *P <* 0.001, two-sided Wilcoxon). **d**, Distribution of Top-1 matching distances at 70% nuclear loss for *π*_0_ (median 27.9 *µ*m) and *π*_1_ (median 21.5 *µ*m). **e**, Spatial cluster maps comparing Xenium ground truth with MINT predictions at 30%, 50%, 70% and 90% nuclear loss. **f**, Cluster-defining marker expression averaged by brain region in Xenium ground truth (top) and MINT prediction at 70% nuclear loss (bottom); right-hand labels report the region-wise correlations.

**Extended Data Fig. 3.**
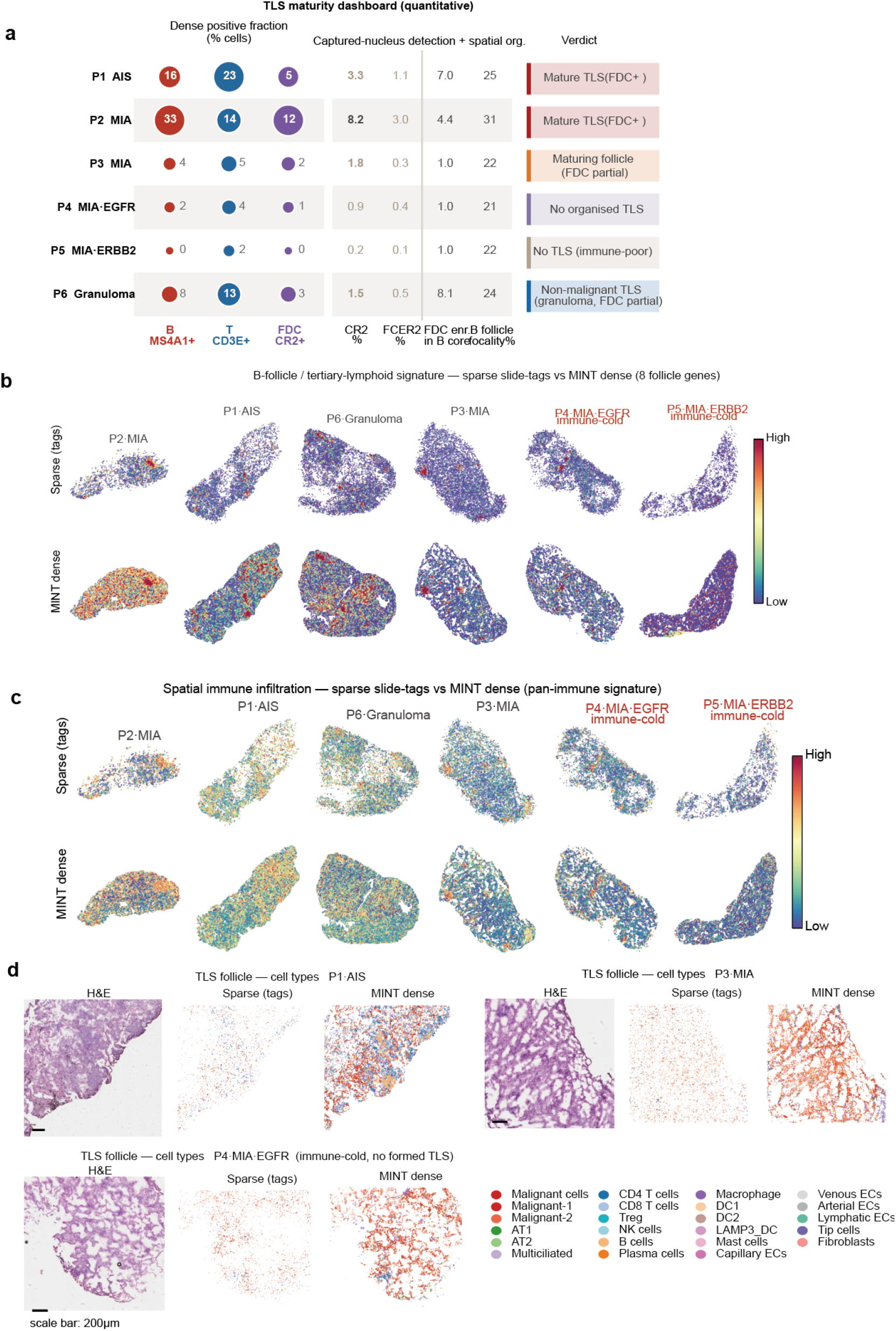
Lung adenocarcinoma: TLS-maturity summary, immune signatures and follicle views. **a**, Tertiary-lymphoid-structure presence and maturity summary across the cohort, including dense positive fractions for B-cell, T-cell and FDC markers, captured-nucleus detection rates for *CR2* and *FCER2*, spatial-organisation scores and the assigned maturity category. **b**, B-follicle/tertiary-lymphoid signature maps in sparse Slide-tags captured nuclei and MINT dense reconstructions across the cohort. **c**, Pan-immune signature maps in sparse captured nuclei and MINT dense reconstructions across the cohort, including driver-mutant cases P4 (*EGFR*) and P5 (*ERBB2*). **d**, Local follicle views showing adjacent-section H&E, sparse captured nuclei and MINT dense cell-type reconstructions for representative samples. FDC-marker detection rates are reported in **a**.

**Extended Data Fig. 4.**
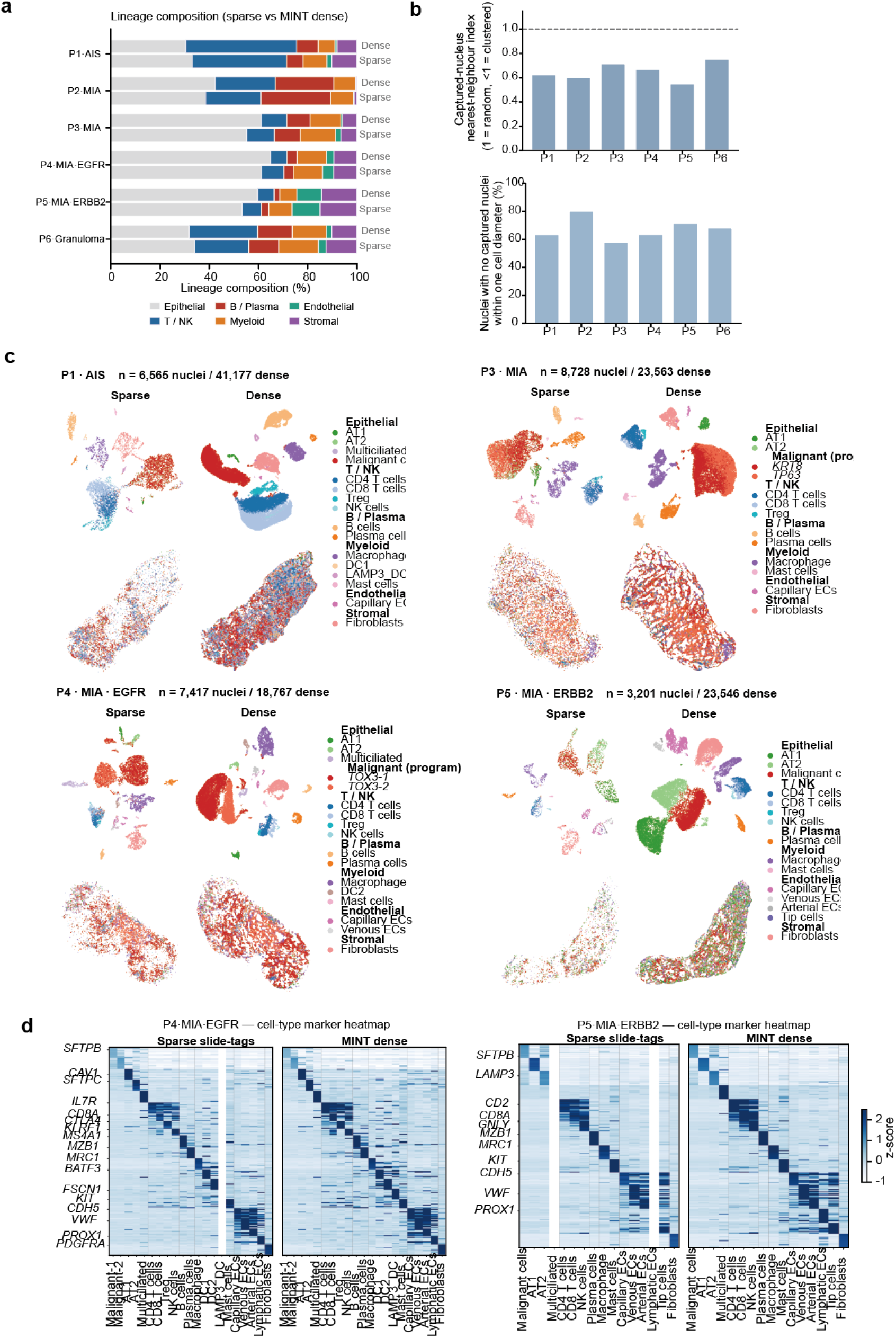
Lung adenocarcinoma cohort: capture coverage, composition and cell-type maps. **a**, Broad lineage composition in sparse Slide-tags captured nuclei and MINT dense reconstructions for samples P1–P6. **b**, Sparse-capture coverage per sample: the fraction of tissue nuclei with no captured nucleus within about one cell diameter, and the nearest-neighbour index of the captured nuclei (below one indicates spatially clustered capture). **c**, UMAP embeddings of sparse captured nuclei and MINT dense reconstructions for four samples, with marker-defined lineage annotations. **d**, Cell-type marker-signature heatmaps for the two immune-cold driver-mutation cases (P4, *EGFR*; P5, *ERBB2*), sparse versus MINT dense.

**Extended Data Fig. 5.**
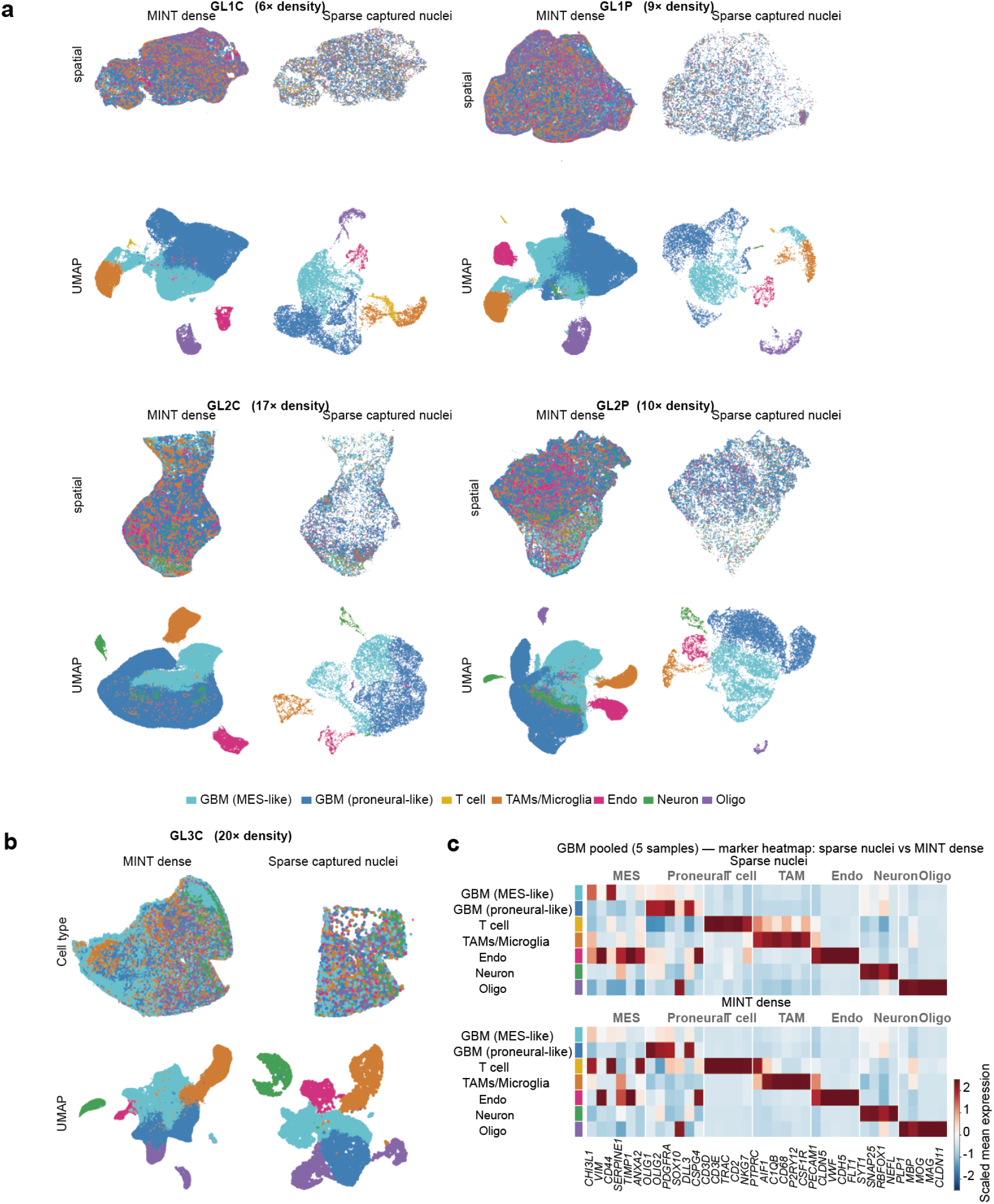
Slide-tags human glioblastoma: dense single-cell reconstruction across five samples. **a**, Sparse Slide-tags captured nuclei versus MINT dense single-cell reconstruction for four samples (GL1C, GL1P, GL2C, GL2P; C, tumour core; P, peritumour; 6- to 17-fold density increase), shown as spatial cell-type maps and UMAP embeddings. Labels group malignant programmes (GBM mesenchymal-like, GBM proneural/OPC-like) and microenvironmental and normal-CNS cell types (T cell, TAM/microglia, endothelial, neuron, oligodendrocyte). **b**, The fifth sample (GL3C, tumour core; ~20-fold density increase), sparse captured nuclei versus MINT dense, shown as spatial cell-type map and UMAP embedding. **c**, Cross-cell-type marker-signature heatmap pooled across five samples, comparing sparse captured nuclei with MINT dense reconstructions (per-cell-type *z*-score). The annotations include malignant programmes, microenvironmental cell types and the non-specific mesenchymal programme.

**Extended Data Fig. 6.**
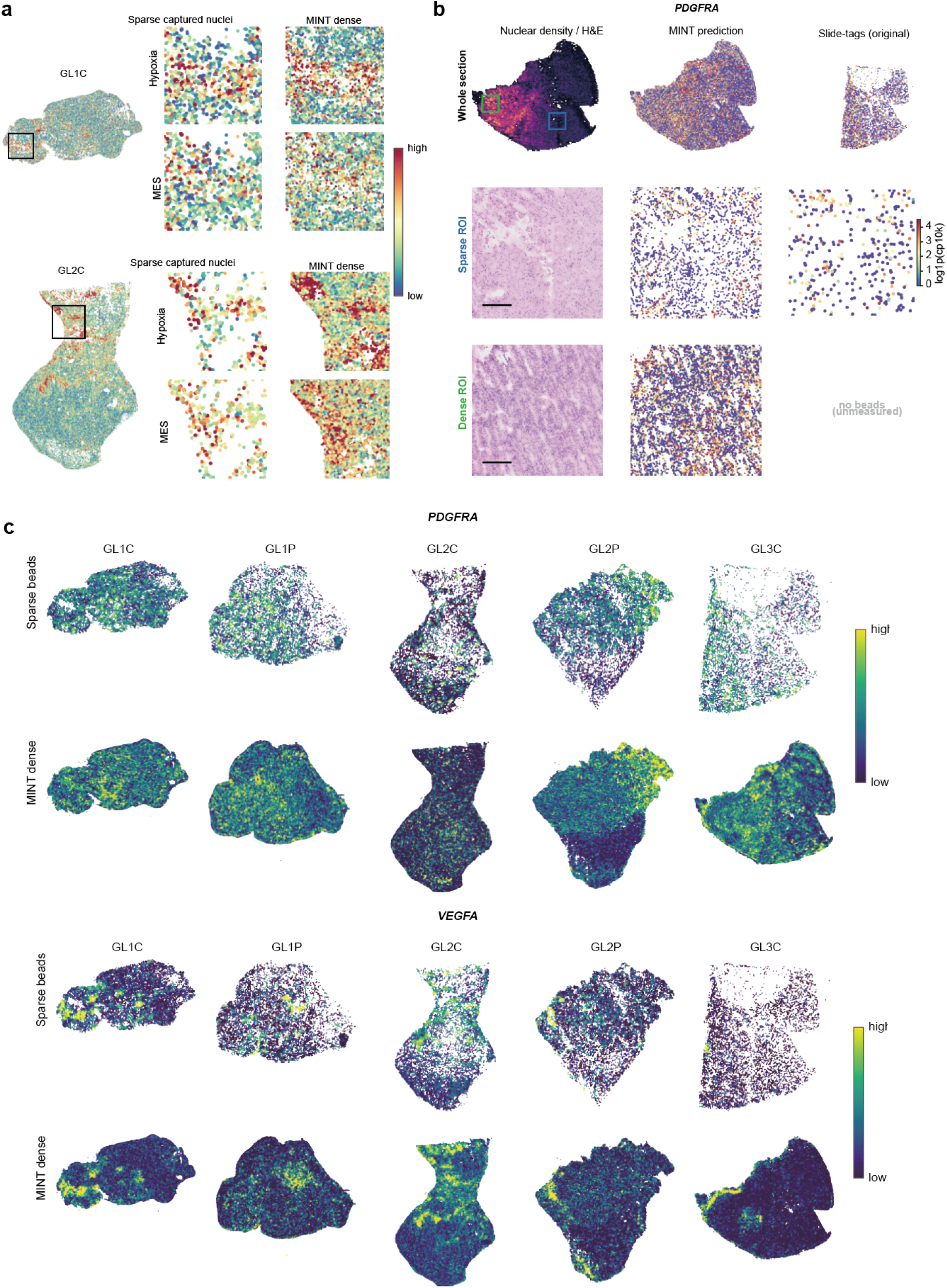
Slide-tags human glioblastoma: transcriptional-programme and marker maps. **a**, Perinecrotic hypoxia and mesenchymal (MES) programme scores for two tumour-core samples (GL1C, GL2C), sparse Slide-tags captured nuclei versus MINT dense reconstruction, with local zooms. **b**, Nuclear-density and *PDGFRA* maps for tumour-core sample GL3C, in which Slide-tags covered about 40% of the section: whole-section nuclear density from H&E, MINT dense prediction and the original Slide-tags capture, with zooms of a sparsely covered and a densely covered region. **c**, Spatial maps of *PDGFRA* and *VEGFA* across the five samples (GL1C, GL1P, GL2C, GL2P, GL3C), sparse captured nuclei versus MINT dense reconstruction. Each sample is independently scaled to compare spatial pattern rather than absolute magnitude (Methods).

**Extended Data Fig. 7.**
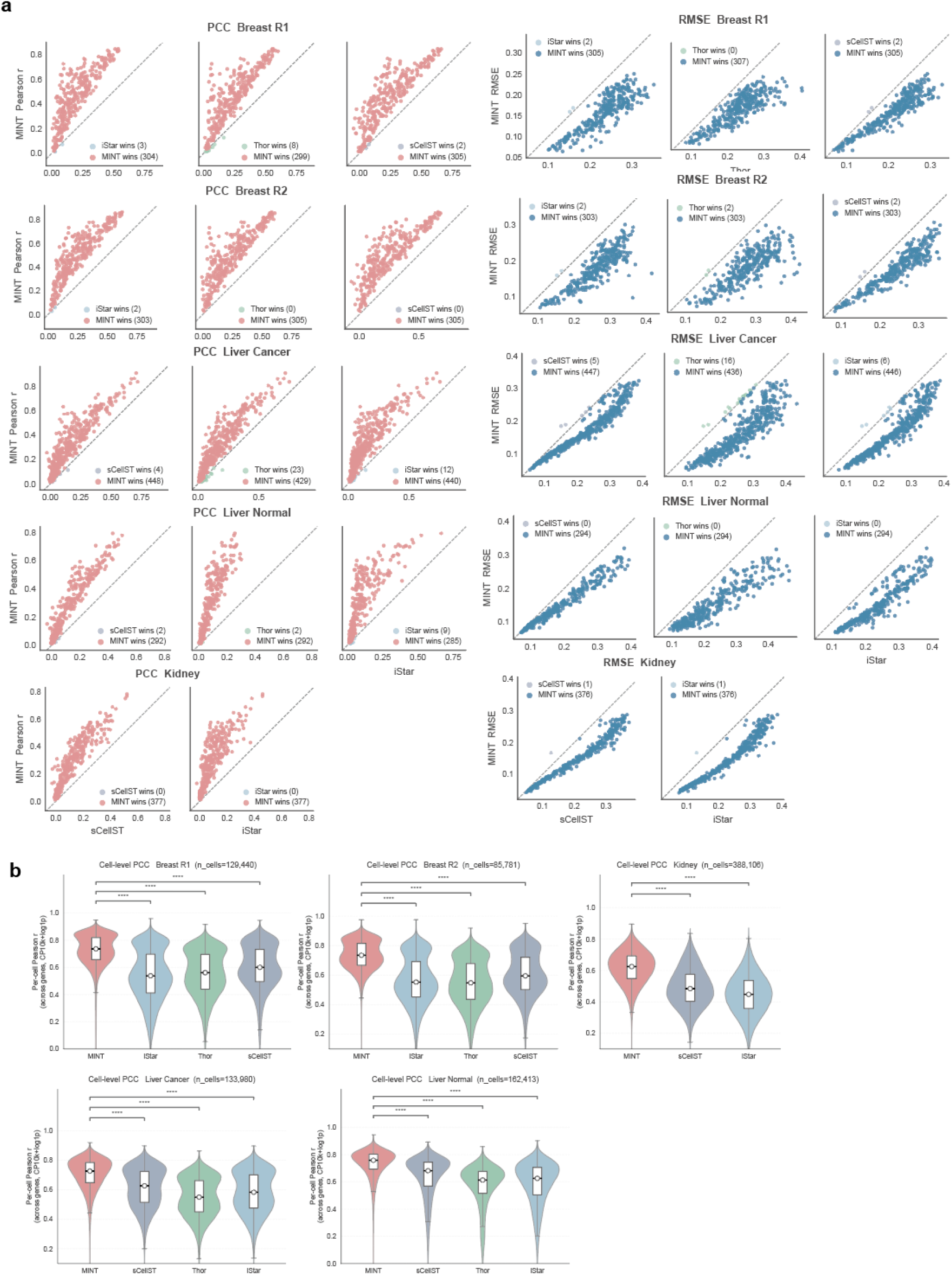
Cross-tissue single-cell benchmark on pseudo-Visium. **a**, Per-gene accuracy of MINT against each competing method across the five Xenium-derived pseudo-Visium sections (breast Rep1, breast Rep2, liver cancer, normal liver and renal carcinoma). For every tissue, paired scatter plots show MINT (*y*-axis) versus iStar, Thor and sCellST (*x*-axis), each point a single gene; the left block reports per-gene Pearson *r* (points above the dashed identity line favour MINT) and the right block per-gene RMSE (points below the line favour MINT), with insets counting the genes each method wins. **b**, Cell-level accuracy quantified as the per-cell Pearson correlation across genes (one value per cell), shown as violins with an inset box plot (centre line, median; box, interquartile range Q1–Q3; whiskers, 1.5× the interquartile range); brackets denote descriptive paired two-sided Wilcoxon signed-rank tests of each competitor against MINT (^∗∗∗∗^*P <* 10^−4^), and the number of evaluated cells is annotated per tissue. Cells within a section were not treated as independent biological replicates (Methods). Thor did not complete on the renal carcinoma section (~388,000 cells) and is therefore absent from the renal carcinoma panels in **a** and **b**. All values are in CP10k+log1p space.

**Extended Data Fig. 8.**
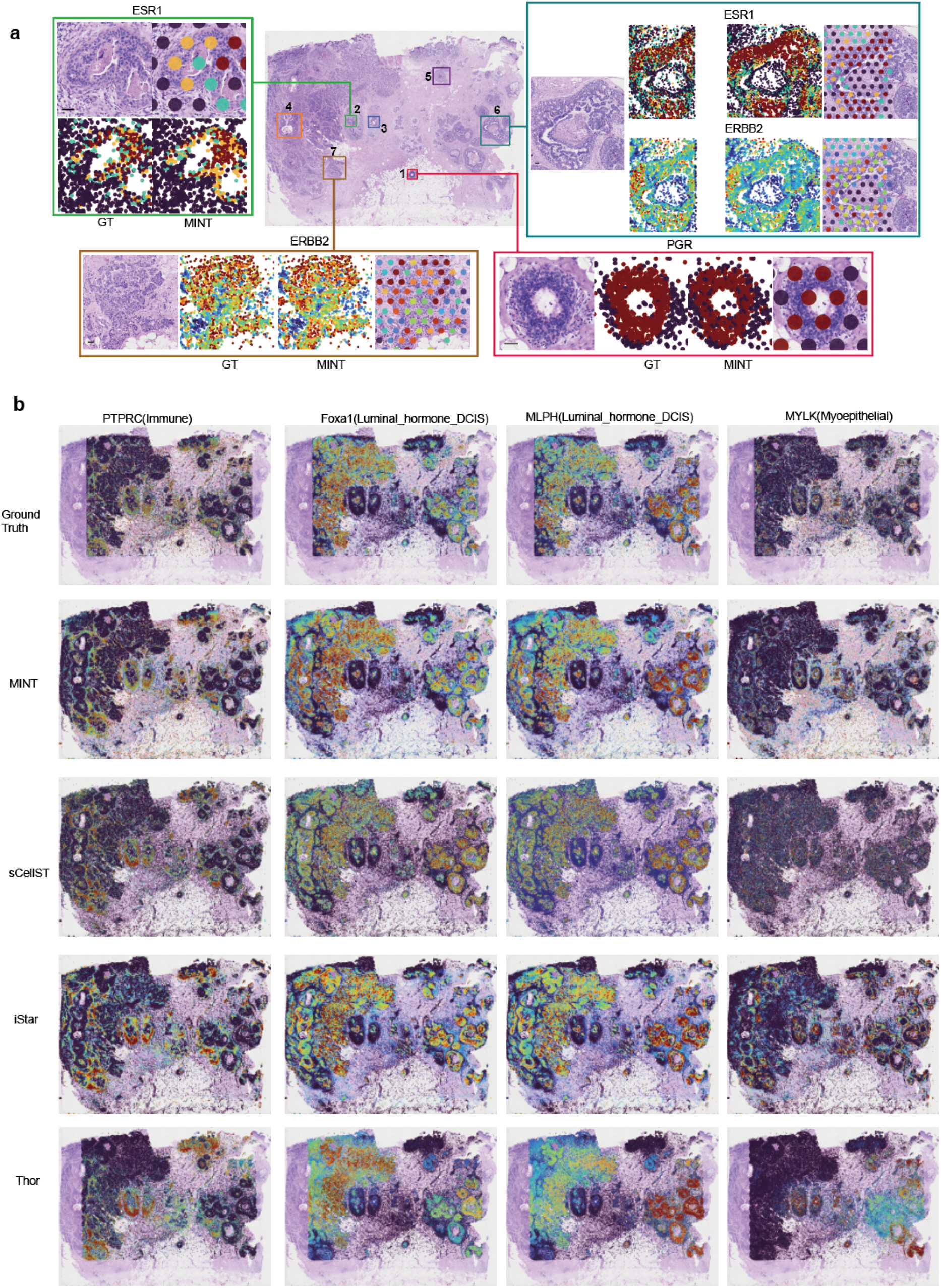
Marker-gene spatial expression maps: breast cancer (Rep1). **a**, Whole-section H&E of the Xenium FFPE human breast cancer slide (Rep1) with regions of interest outlined; for three regions, paired close-ups compare Xenium ground truth with MINT single-cell predictions for the hormone-receptor and HER2 markers *ESR1, ERBB2* and *PGR*. **b**, Tissue-wide spatial maps for four lineage markers (*PTPRC*, immune; *FOXA1* and *MLPH*, luminal; *MYLK*, myoepithelial). Rows show Xenium ground truth, MINT, sCellST, iStar and Thor. Per-cell expression is overlaid on the downsampled H&E and coloured by expression level, with the colour scale clipped at the 1st–95th percentiles (Methods).

**Extended Data Fig. 9.**
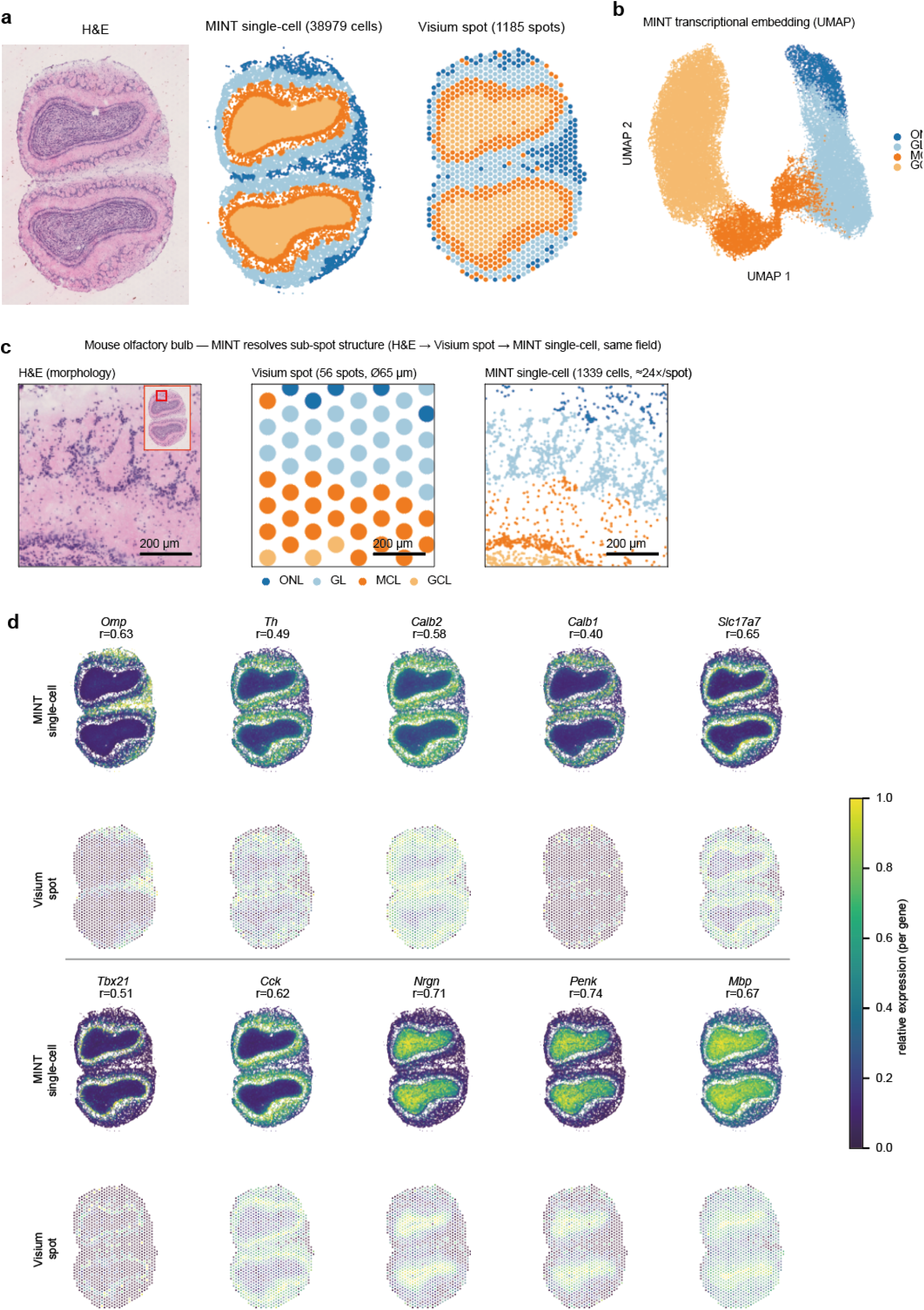
Real Visium mouse olfactory bulb. **a**, H&E and spatially aware clustering maps for MINT single cells and Visium spots from the same section. **b**, UMAP of MINT transcriptional profiles. **c**, A sub-spot view resolving 56 spots into 1,339 single cells (~ 24 cells per spot) across the ONL, GL, MCL and GCL layers. **d**, Layer-marker spatial maps for MINT single cells and Visium spots, with per-gene correlations.

**Extended Data Fig. 10.**
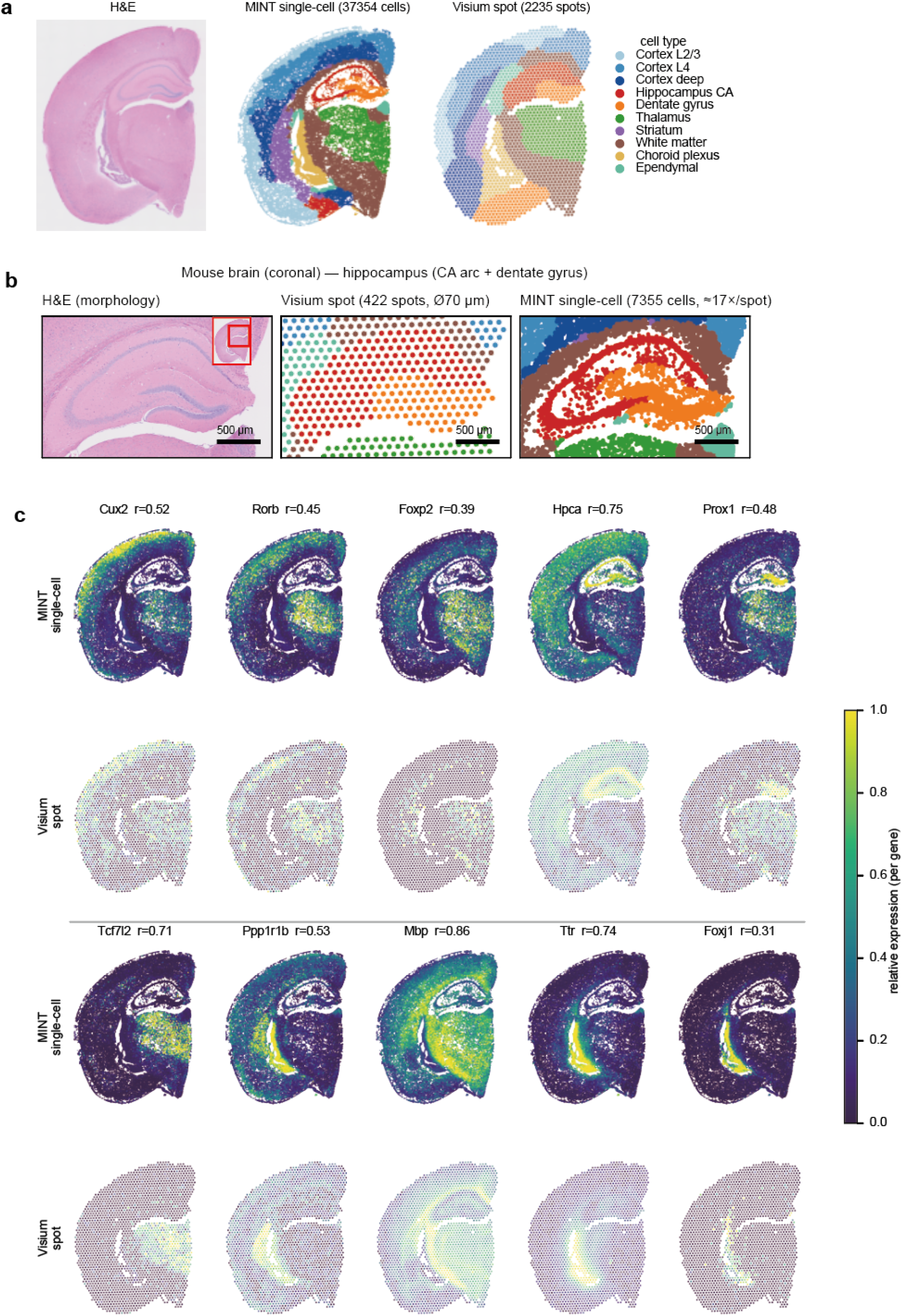
Real Visium mouse coronal brain. **a**, H&E and anatomical-domain maps for MINT single cells and Visium spots, including cortical layers, hippocampal CA arc and dentate gyrus, thalamic and striatal nuclei, and white matter. **b**, Hippocampal view resolving 422 spots into 7,355 cells. **c**, Region-marker spatial maps with per-gene correlations.

