## Supplementary Information for "MINT infers the latent single-cell spatial transcriptome from paired histology"

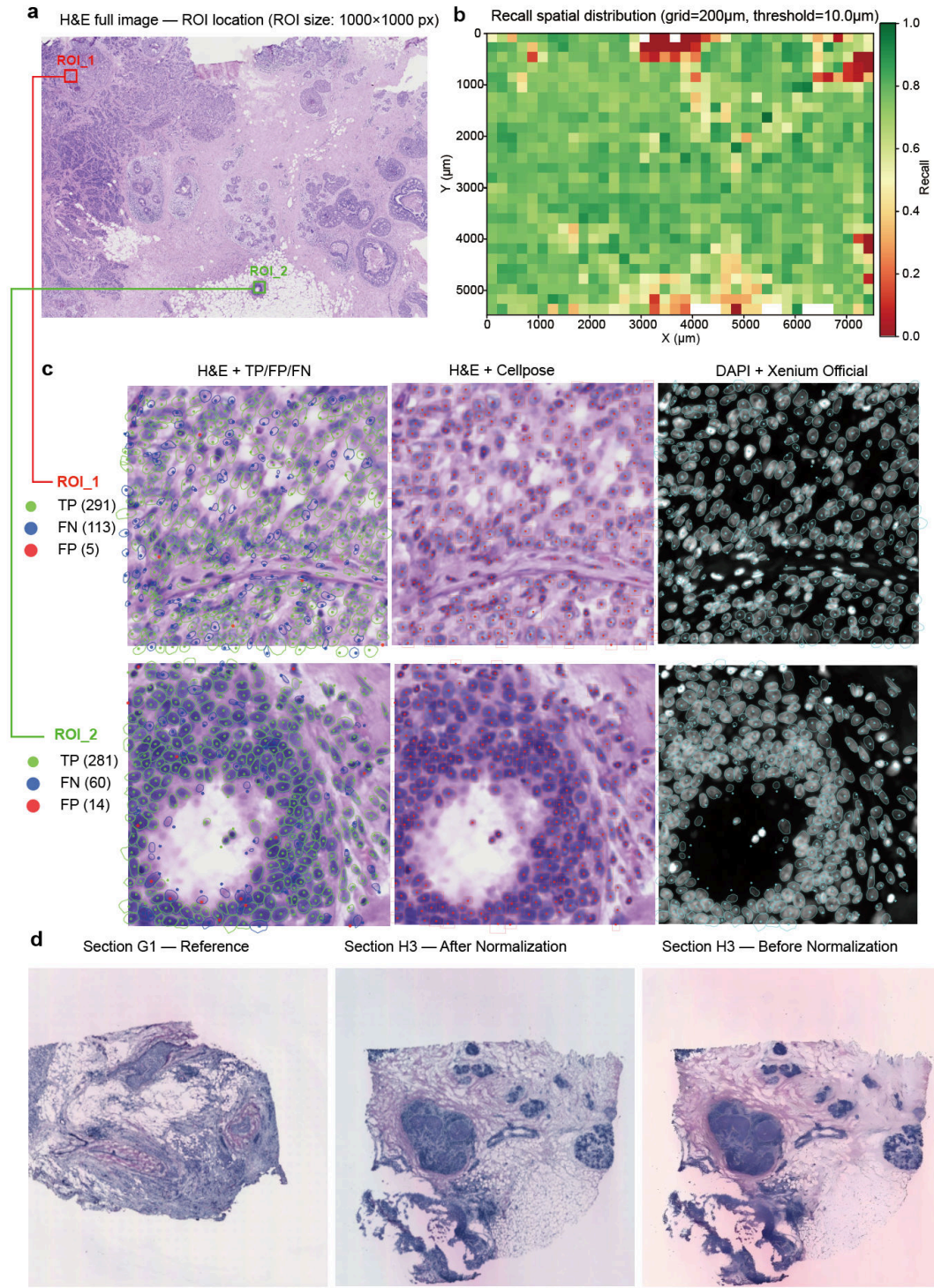

**Supplementary Fig. S1 Preprocessing quality assessment: nucleus segmentation and stain normalisation.** **a**, Full H&E image of the Xenium FFPE Human Breast Cancer tissue section with two regions of interest (ROIs) marked for detailed inspection. **b**, Spatial heatmap of segmentation recall across the tissue, computed on a 200-μm grid. Recall is defined as the fraction of Xenium DAPI-segmented nuclei matched to a Cellpose-detected nucleus by mutual nearest-neighbour (MNN) matching at a 10-μm distance threshold. Overall segmentation achieved 96.4% precision and 74.7% recall ($F1 = 0.84$) at this threshold. **c**, Detailed views of the two ROIs. For each ROI, three panels show: H&E with true positive (TP), false positive (FP), and false negative (FN) annotations; H&E with Cellpose segmentation overlays; and DAPI fluorescence with Xenium official segmentation boundaries. Representative H&E images from the HER2ST dataset: reference section (G1, left), target section after normalisation (centre), and target section before normalisation (right). Reinhard normalisation aligns the colour distribution of each section to the reference in CIE Lab colour space (Methods).

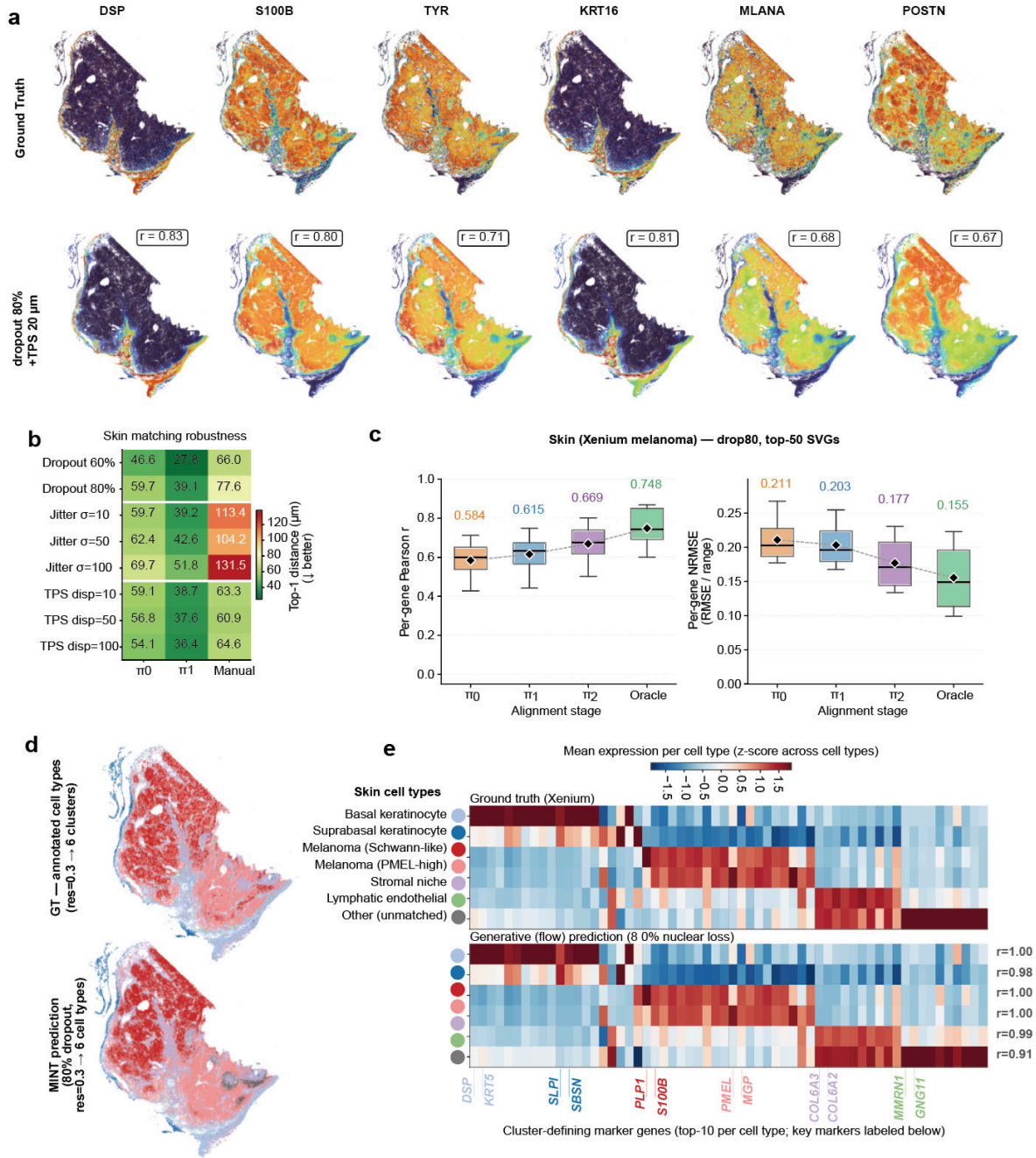

**Supplementary Fig. S2 FUGW alignment validation on Xenium human skin melanoma pseudo-Slide-tags data (80% nuclear loss).** **a**, Spatial expression of six top-performing marker genes (*DSP*, *S100B*, *TYR*, *KRT16*, *MLANA*, *POSTN*) at ground truth (top) and MINT prediction under 80% nuclear loss + TPS $\sim 20 \mu\text{m}$ perturbation (bottom). Per-gene Pearson $r$ shown for each gene. **b**, Skin matching robustness across dropout (60%, 80%), Gaussian jitter ($\sigma \in \{10, 50, 100\} \mu\text{m}$), and TPS median-displacement magnitudes ($\{10, 50, 100\} \mu\text{m}$). Each row shows Top-1 mean distance ($\mu\text{m}$) for $\pi_0$, $\pi_1$, and the manual landmark baseline. **c**, Per-gene Pearson correlation (left) and per-gene normalised RMSE (right) on the 50 spatially variable genes held out from alignment (ranks 51–100 by Moran's $I$, disjoint from the top-50 alignment panel) across the four alignment stages ($\pi_0$, $\pi_1$, $\pi_2$, GT). Normalised RMSE uses the 1–99 percentile range of log<sub>1p</sub> ground-truth expression as the per-gene normalisation factor. **d**, Ground-truth cell-type annotation (top) and MINT-predicted cell-type clusters (bottom; Leiden $\text{res}=0.3$, six clusters) at 80% nuclear loss. At a 50% overlap threshold all six predicted clusters mapped to ground-truth cell types. **e**, Heatmap of cluster-defining marker gene expression averaged per cell type (top: ground truth; bottom: MINT prediction at 80% nuclear loss; per-cell-type $r$ shown on the right).

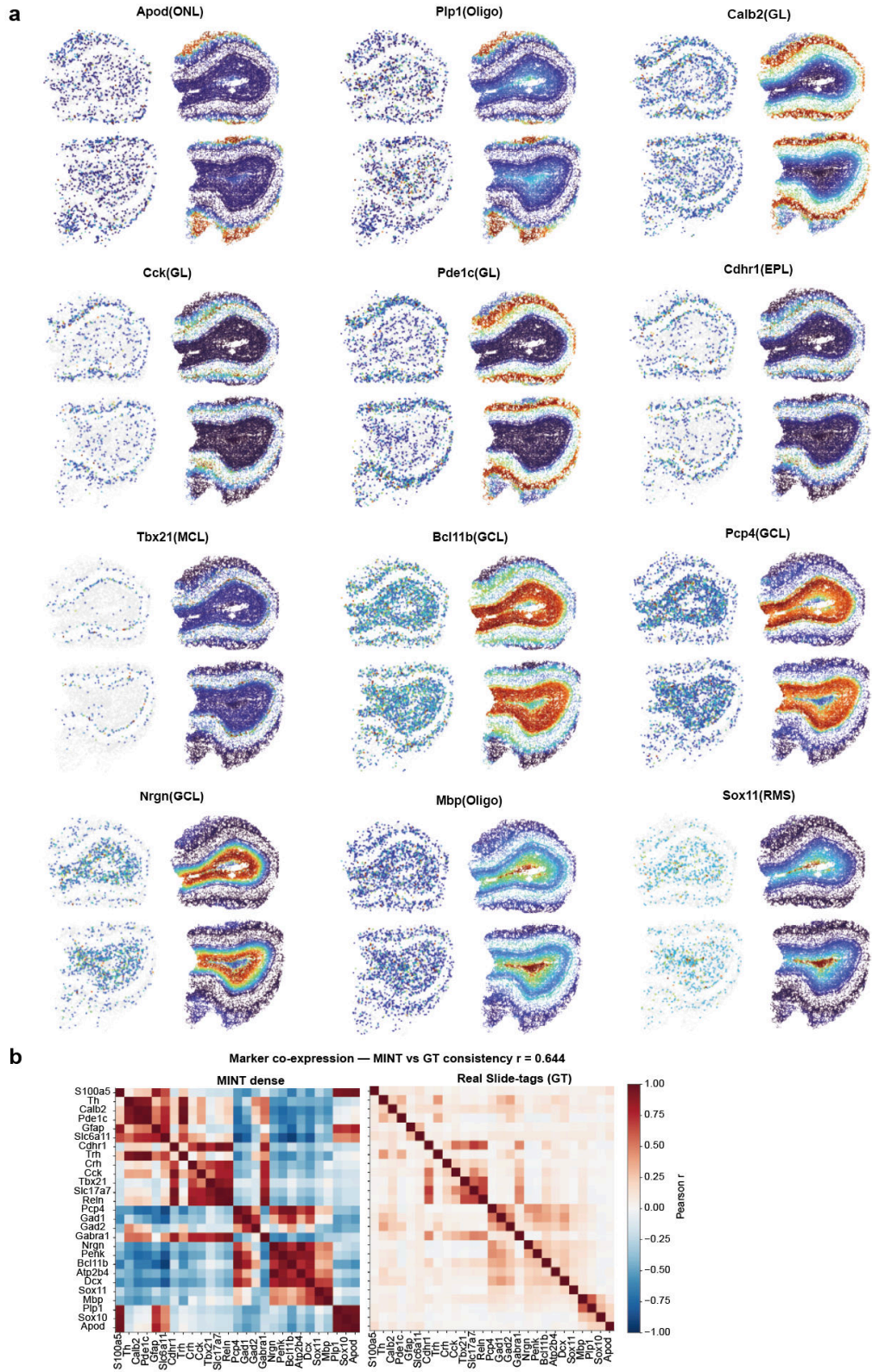

**Supplementary Fig. S3 Slide-tags mouse olfactory bulb: per-marker reconstruction and marker co-expression.** **a**, Spatial reconstruction of 12 layer-marker genes (*Apod*, *Plp1*, *Calb2*, *Cck*, *Pde1c*, *Cdhr1*, *Tbx21*, *Bcl11b*, *Pcp4*, *Nrgn*, *Mbp* and *Sox11*) in the two olfactory-bulb lobes. For each marker, sparse Slide-tags captured nuclei are shown at left and the MINT dense reconstruction at right. Sparse maps show captured nuclei with non-zero counts over a grey tissue outline. **b**, Marker-marker co-expression matrices (Pearson correlation across the marker panel) for the MINT dense reconstruction and real Slide-tags data; their off-diagonal structures have a Pearson correlation of $r = 0.64$.

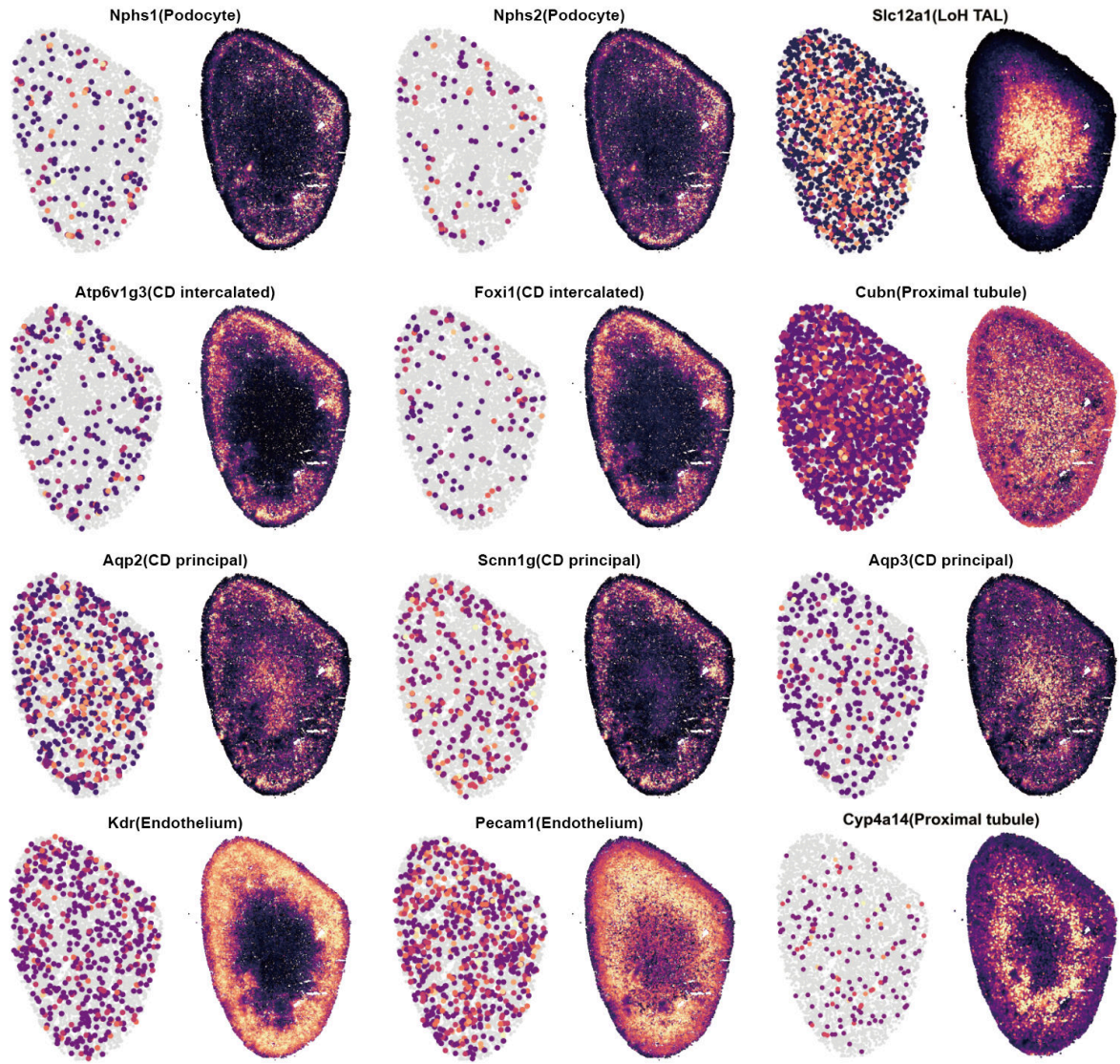

**Supplementary Fig. S4 Slide-tags mouse kidney: nephron-segment marker reconstruction.** Spatial maps of 12 nephron and vascular marker genes in sparse Slide-tags captured nuclei (left of each pair) and the MINT dense reconstruction (right; 76,073 nuclei). Markers represent podocyte (*Nphs1*, *Nphs2*), proximal tubule (*Cubn*, *Cyp4a14*), loop-of-Henle thick ascending limb (*Slc12a1*), collecting-duct intercalated cells (*Atp6v1g3*, *Foxi1*), collecting-duct principal cells (*Aqp2*, *Aqp3*, *Scnn1g*) and endothelium (*Kdr*, *Pecam1*). Sparse maps show captured nuclei with non-zero counts over a grey tissue outline.

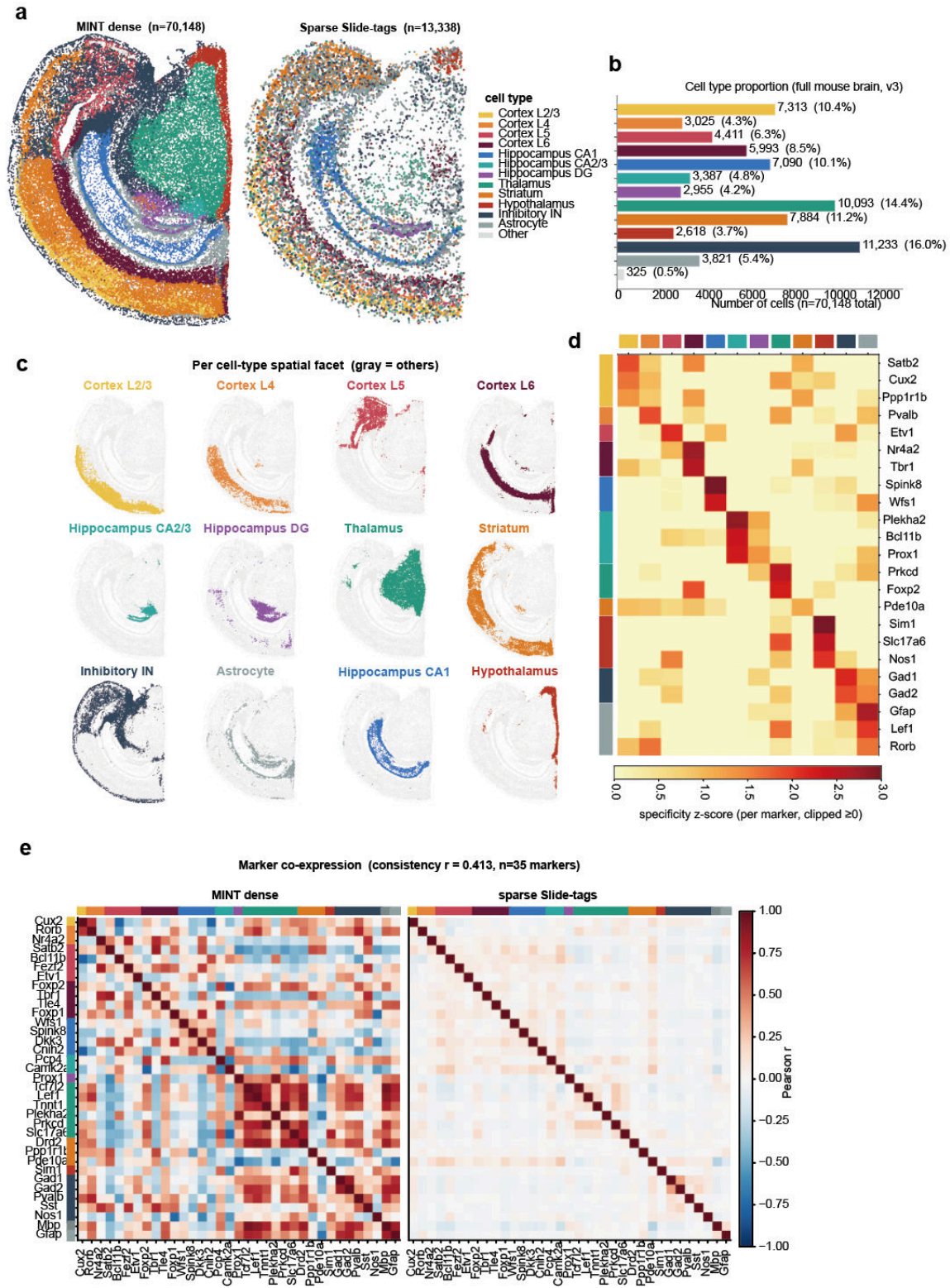

**Supplementary Fig. S5 Slide-tags mouse brain: downstream clustering, cell-type composition and marker fidelity.** **a**, Spatial cell-type maps for the MINT dense reconstruction (70,148 nuclei) and the sparse Slide-tags captured nuclei (13,338 nuclei), coloured by twelve annotated cell types: cortical layers L2/3, L4, L5 and L6; hippocampal subfields CA1, CA2/3 and the dentate gyrus; thalamus; striatum; hypothalamus; inhibitory interneurons; and astrocytes (with an unassigned class). **b**, Cell-type proportions across the reconstructed brain (number of nuclei per type). **c**, Per-cell-type spatial layout: each panel highlights one cell type against all remaining nuclei in grey, showing that laminar and regional types occupy spatially coherent territories. **d**, Marker-specificity heatmap showing per-marker, per-cell-type specificity z-scores for the curated regional marker panel. **e**, Marker-marker co-expression matrices (Pearson correlation) for the MINT dense reconstruction and the sparse Slide-tags data; the off-diagonal structure is preserved between them (consistency $r = 0.41$ across the marker panel).

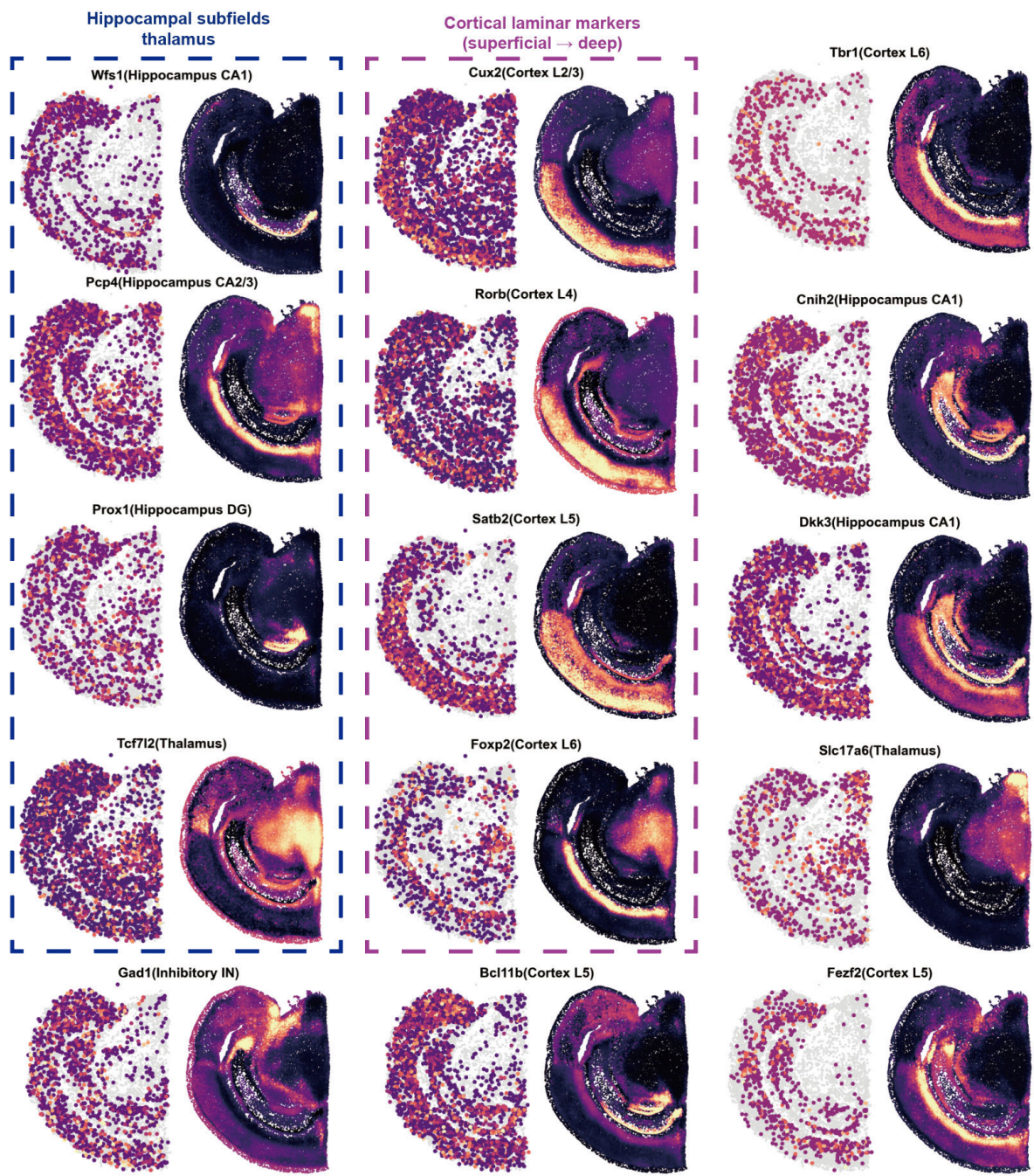

**Supplementary Fig. S6 Slide-tags mouse brain: marker-gene spatial maps.** Sparse Slide-tags versus MINT dense reconstruction for cortical laminar markers (*Cux2/Rorb/Satb2/Foxp2*), hippocampal subfields (*Wfs1/Pcp4/Prox1*) and thalamus (*Tcf7l2*).

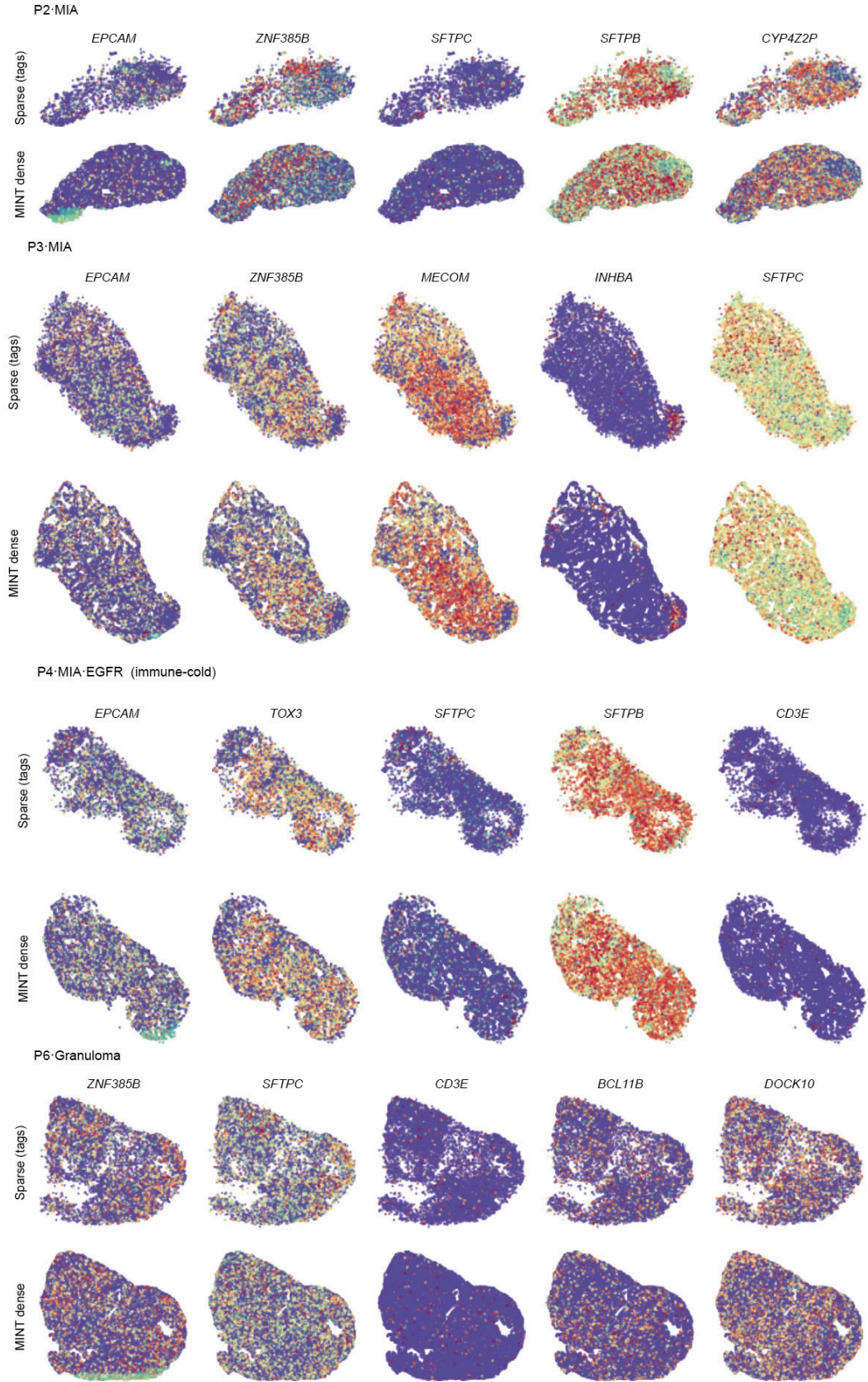

**Supplementary Fig. S7 Lung adenocarcinoma: per-sample spatial marker maps (P2, P3, P4, P6).** Sparse Slide-tags captured nuclei versus MINT dense reconstruction for representative epithelial, lung-lineage and immune-associated marker genes in four samples. These maps provide gene-level spatial context for the cell-type and immune-programme summaries in Fig. 4.

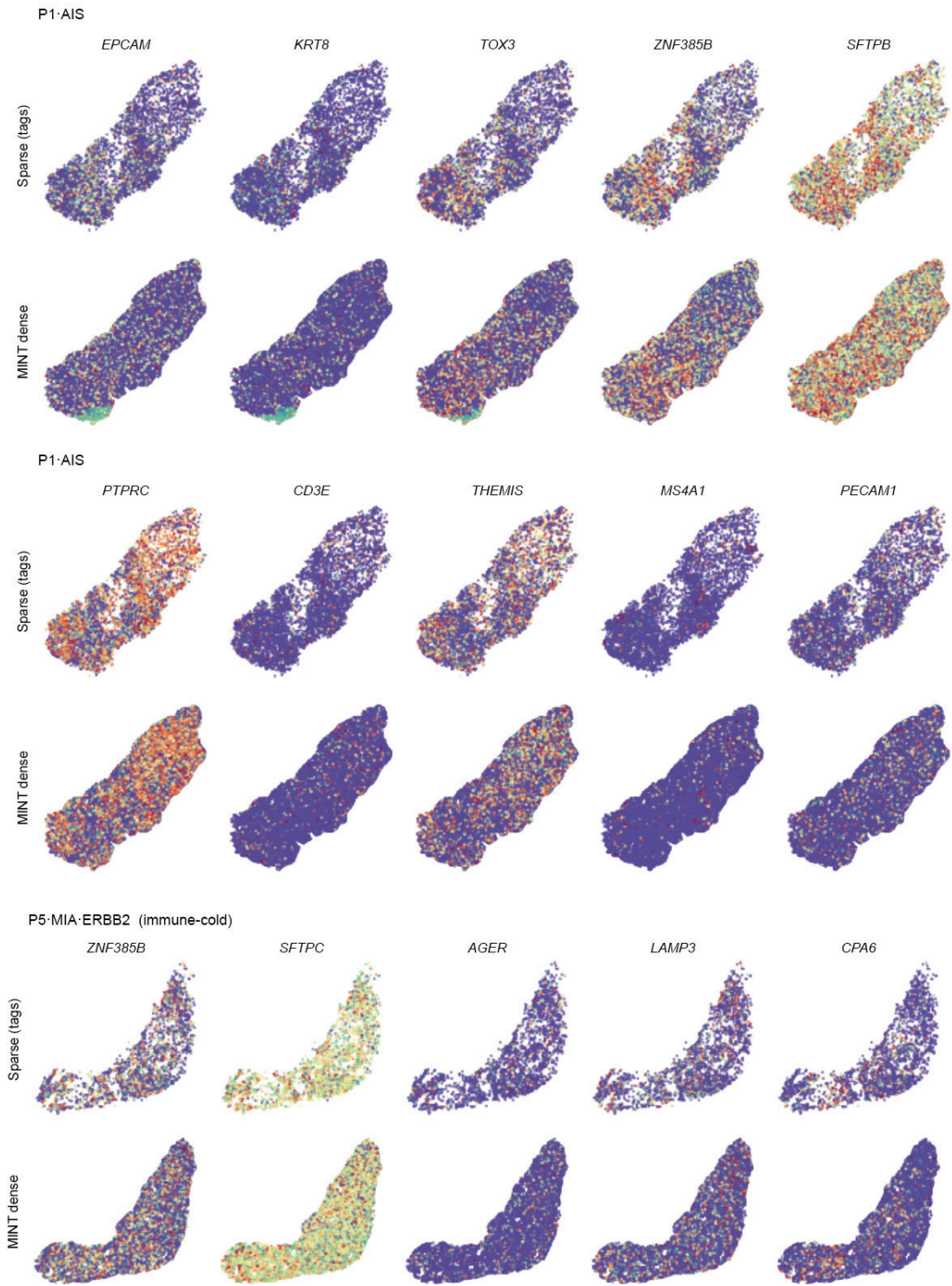

**Supplementary Fig. S8 Lung adenocarcinoma: per-sample spatial marker maps (P1, P5).** Sparse Slide-tags captured nuclei versus MINT dense reconstruction for representative epithelial, immune and vascular or dendritic-cell marker genes in the adenocarcinoma in situ (P1) and the immune-cold *ERBB2*-mutant case (P5).

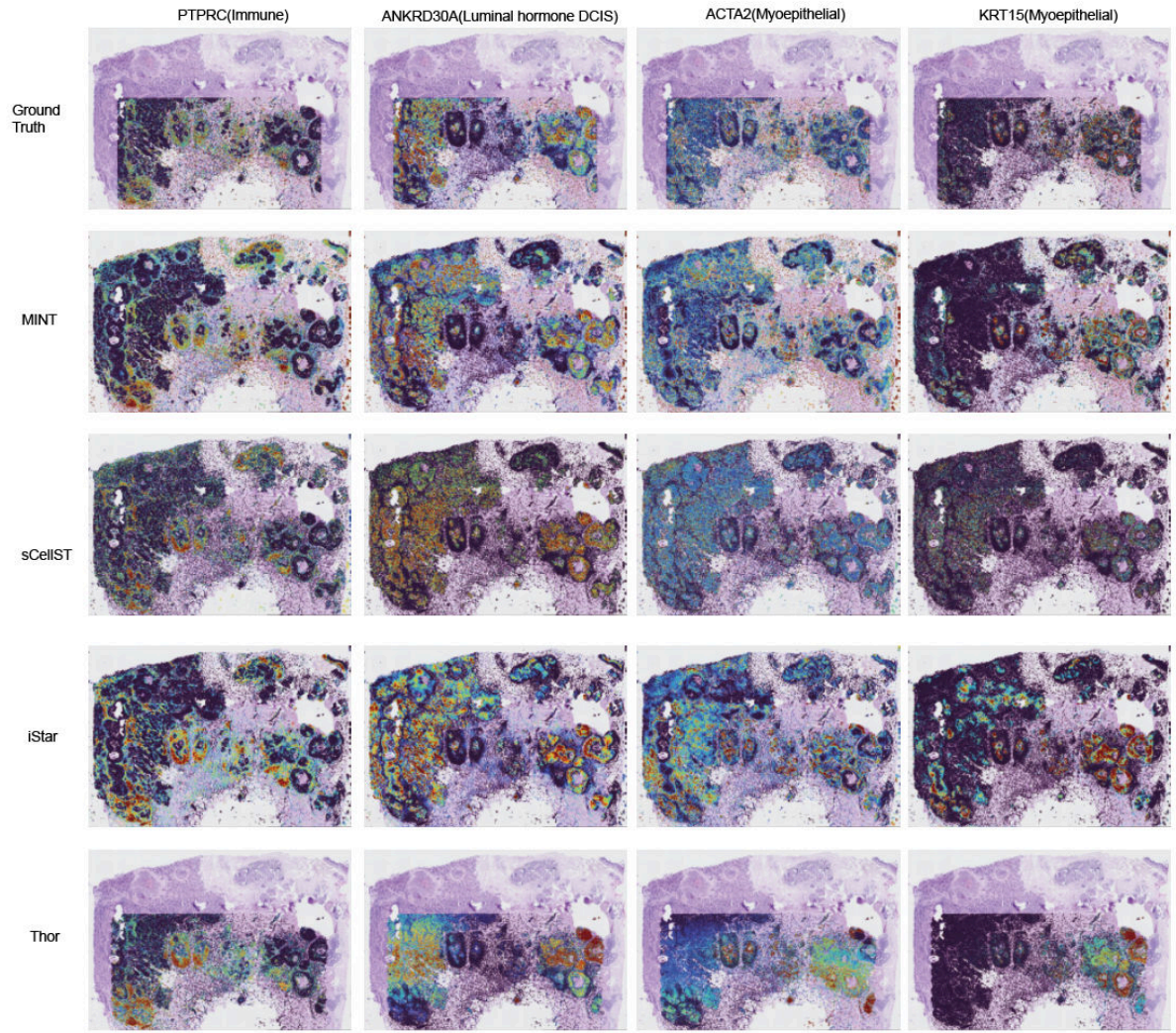

**Supplementary Fig. S9 Marker-gene spatial expression maps: breast cancer (Rep2).** Tissue-wide spatial maps for four lineage markers (*PTPRC*, immune; *ANKRD30A*, luminal; *ACTA2* and *KRT15*, myoepithelial). Columns correspond to markers; rows show Xenium ground truth, MINT, sCellST, iStar and Thor. Per-cell expression is overlaid on the downsampled H&E and coloured by expression level, with the colour scale clipped at the 1st–95th percentiles (Methods).

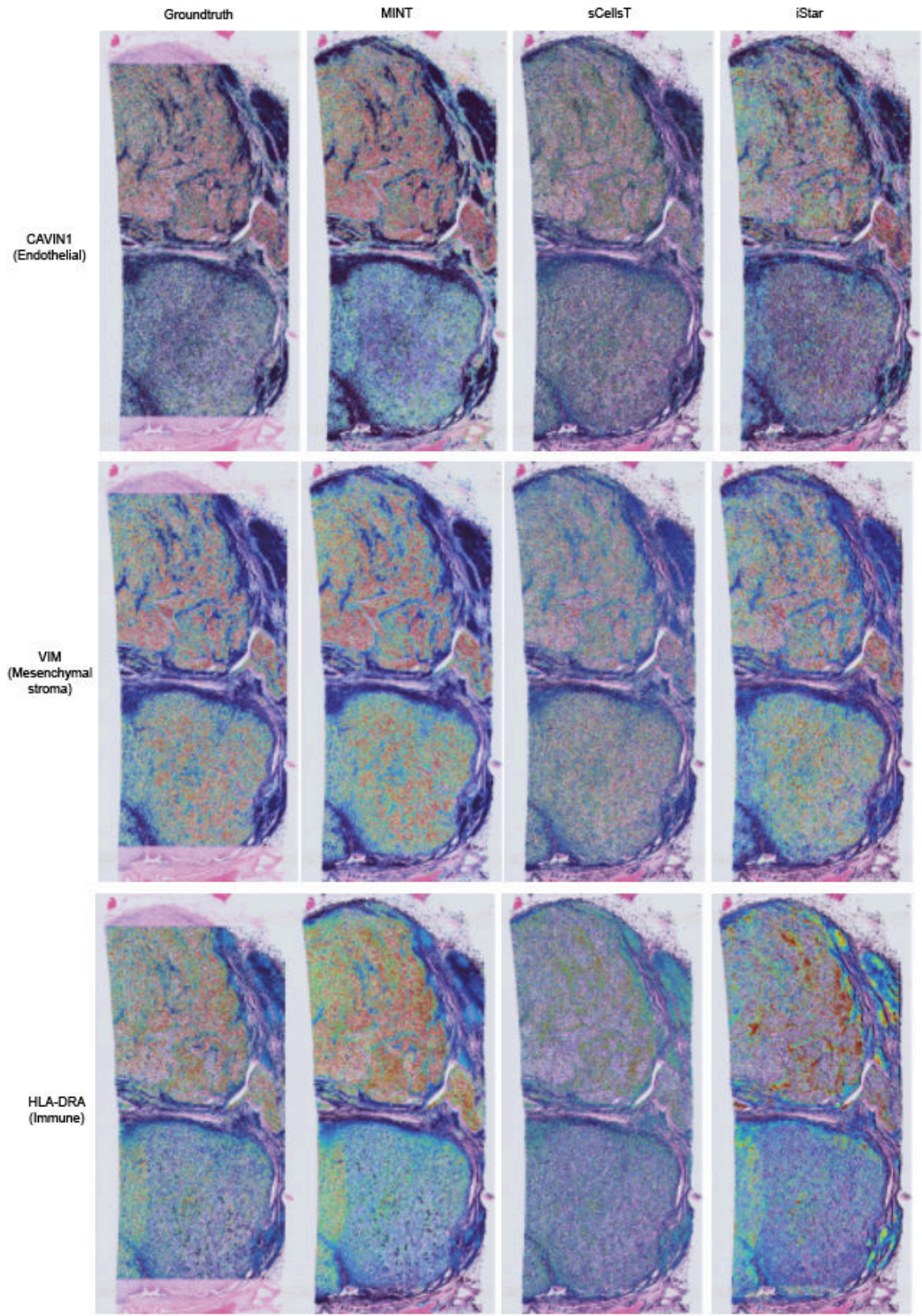

**Supplementary Fig. S10 Marker-gene spatial expression maps: renal carcinoma.** Tissue-wide spatial maps for three lineage markers (*CAVIN1*, endothelial; *VIM*, mesenchymal stroma; *HLA-DRA*, immune). Columns show Xenium ground truth, MINT, sCellST and iStar; Thor is omitted because it did not complete on this section (~388,000 cells; Methods). Per-cell expression is overlaid on the downsampled H&E and coloured by expression level, with the colour scale clipped at the 1st–95th percentiles (Methods).

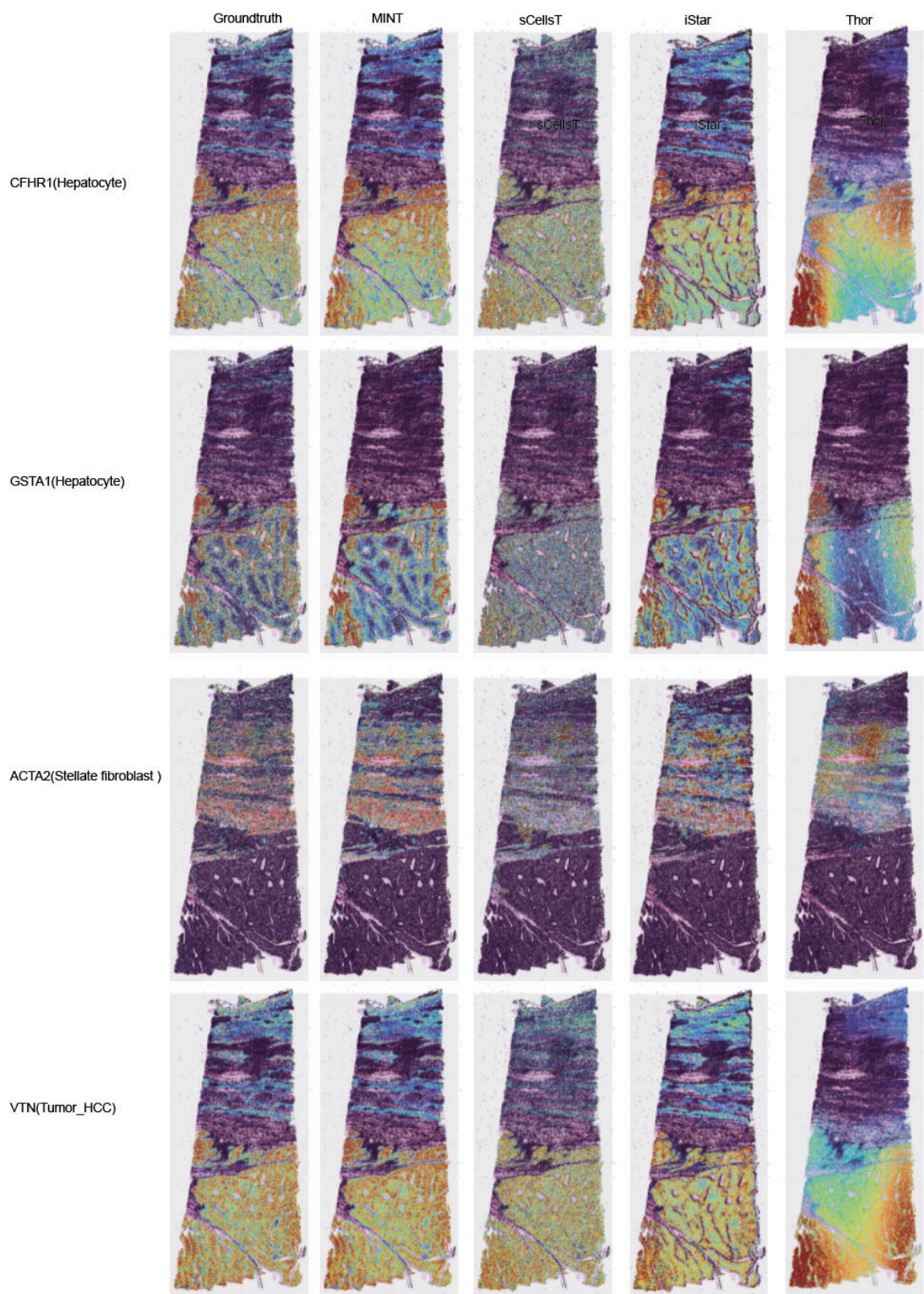

**Supplementary Fig. S11 Marker-gene spatial expression maps: liver cancer.** Tissue-wide spatial maps for four lineage markers (*CFHR1* and *GSTA1*, hepatocyte; *ACTA2*, stellate/fibroblast; *VTN*, tumour). Columns show Xenium ground truth, MINT, sCellST, iStar and Thor. Per-cell expression is overlaid on the downsampled H&E and coloured by expression level, with the colour scale clipped at the 1st–95th percentiles (Methods).

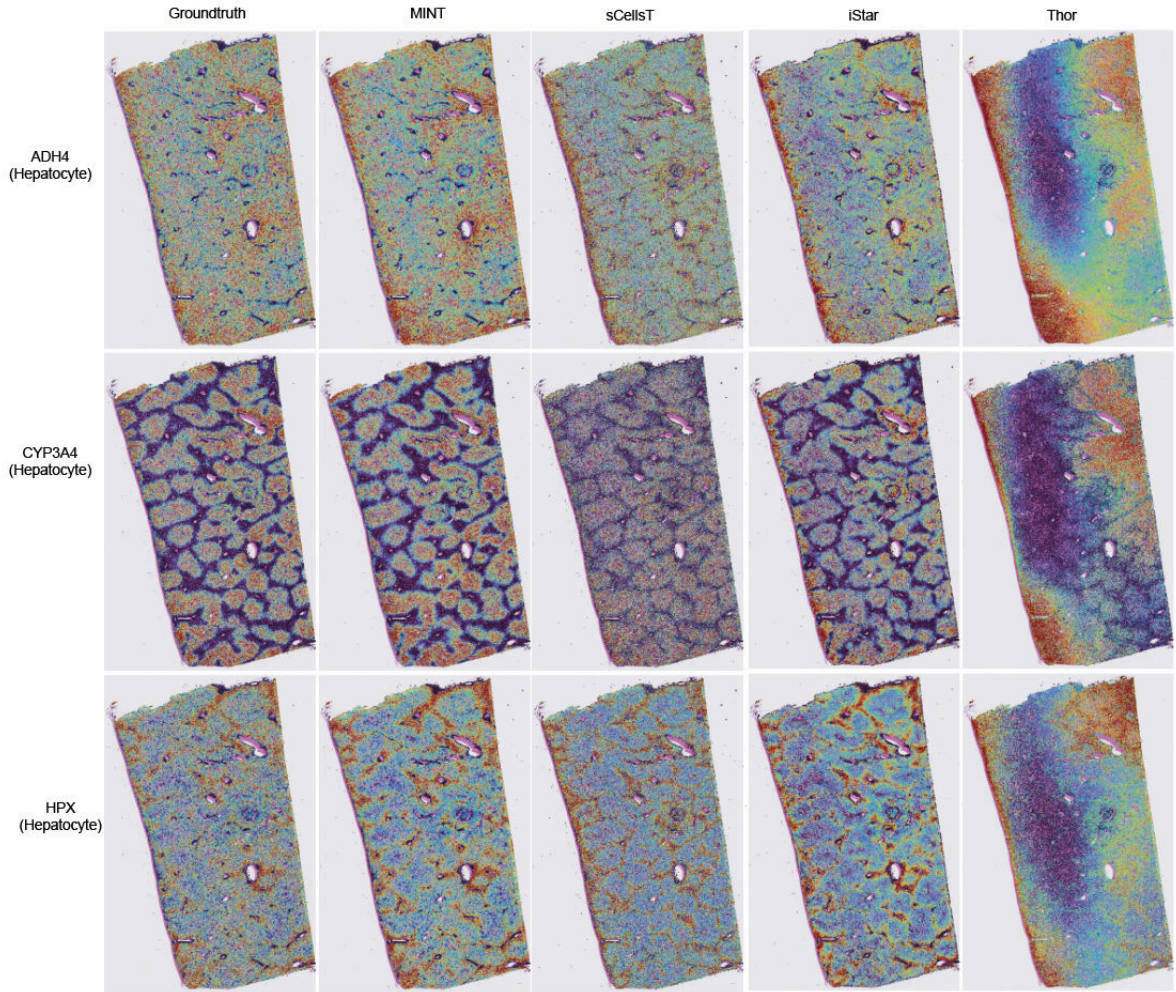

**Supplementary Fig. S12 Marker-gene spatial expression maps: normal liver.** Tissue-wide spatial maps for three hepatocyte markers (*ADH4*, *CYP3A4*, *HPX*). Columns show Xenium ground truth, MINT, sCellST, iStar and Thor. Per-cell expression is overlaid on the downsampled H&E and coloured by expression level, with the colour scale clipped at the 1st–95th percentiles (Methods).

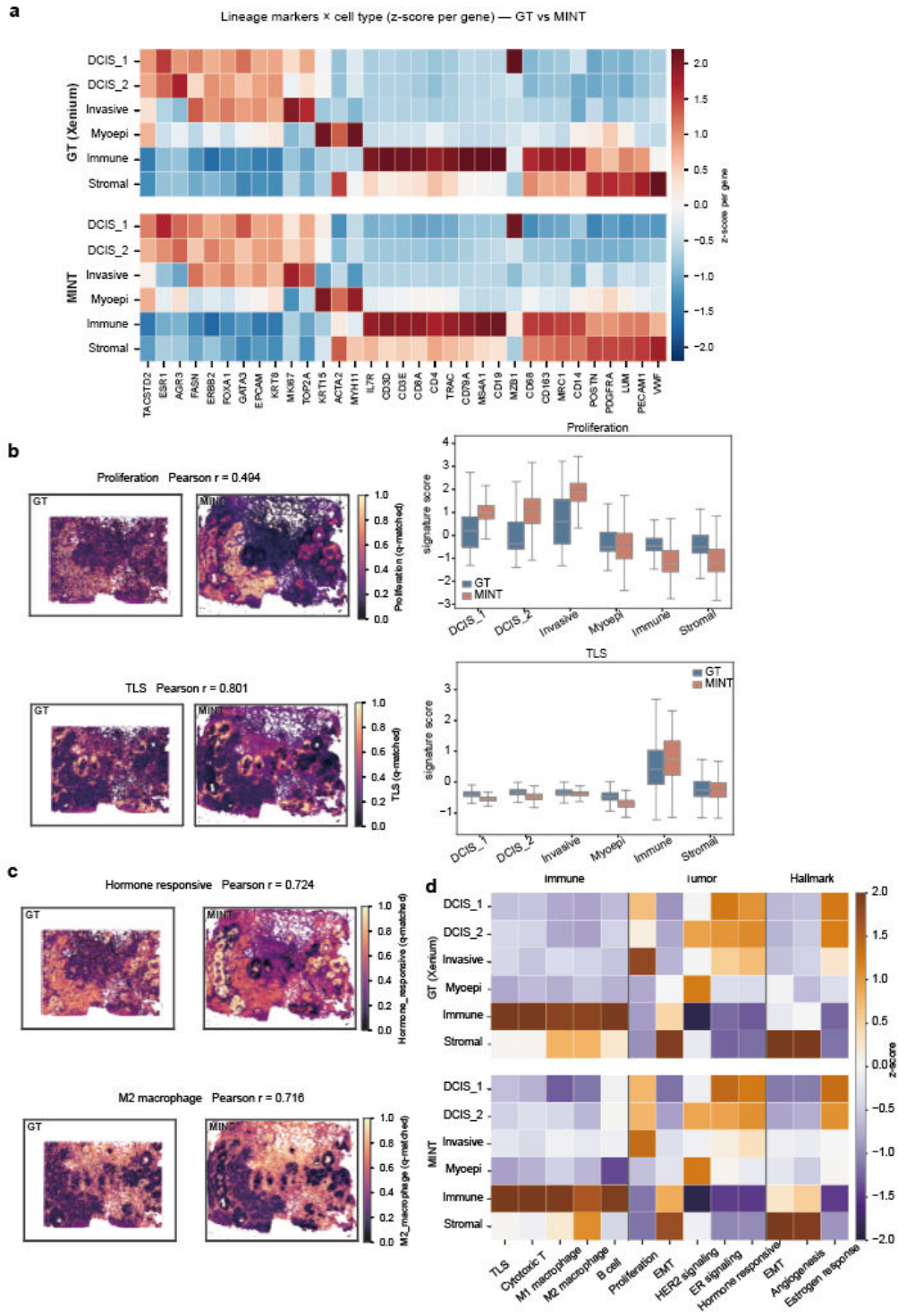

**Supplementary Fig. S13 Downstream single-cell analysis of MINT predictions on human breast cancer (Rep1).** Xenium ground-truth and MINT-predicted cells were each clustered (Leiden) and annotated into six populations: luminal-like and basal-leaning ductal carcinoma in situ, invasive carcinoma, myoepithelial, immune and stromal cells, in a shared CP10k+log1p space. **a**, Lineage-marker by cell-type expression heatmap for ground truth (top) and MINT (bottom); rows are the six populations and columns are canonical lineage markers grouped by epithelial, proliferation, myoepithelial, T-cell, B/plasma, macrophage, fibroblast and endothelial identity, coloured by per-gene z-score. **b**, Spatial maps of the proliferation and tertiary-lymphoid-structure signature scores in ground truth (left) and MINT (right) on shared histology coordinates, with the ground-truth-versus-MINT Pearson correlation annotated on each map and box plots of each signature across the six cell types (centre line, median; box, interquartile range Q1–Q3; whiskers, $1.5 \times$ the interquartile range; outliers not shown). **c**, Spatial maps of the hormone-responsive and M2-macrophage signatures in the same layout. **d**, Signature-by-cell-type z-score heatmap of immune programmes scored across the six cell types (ground truth, top; MINT, bottom).

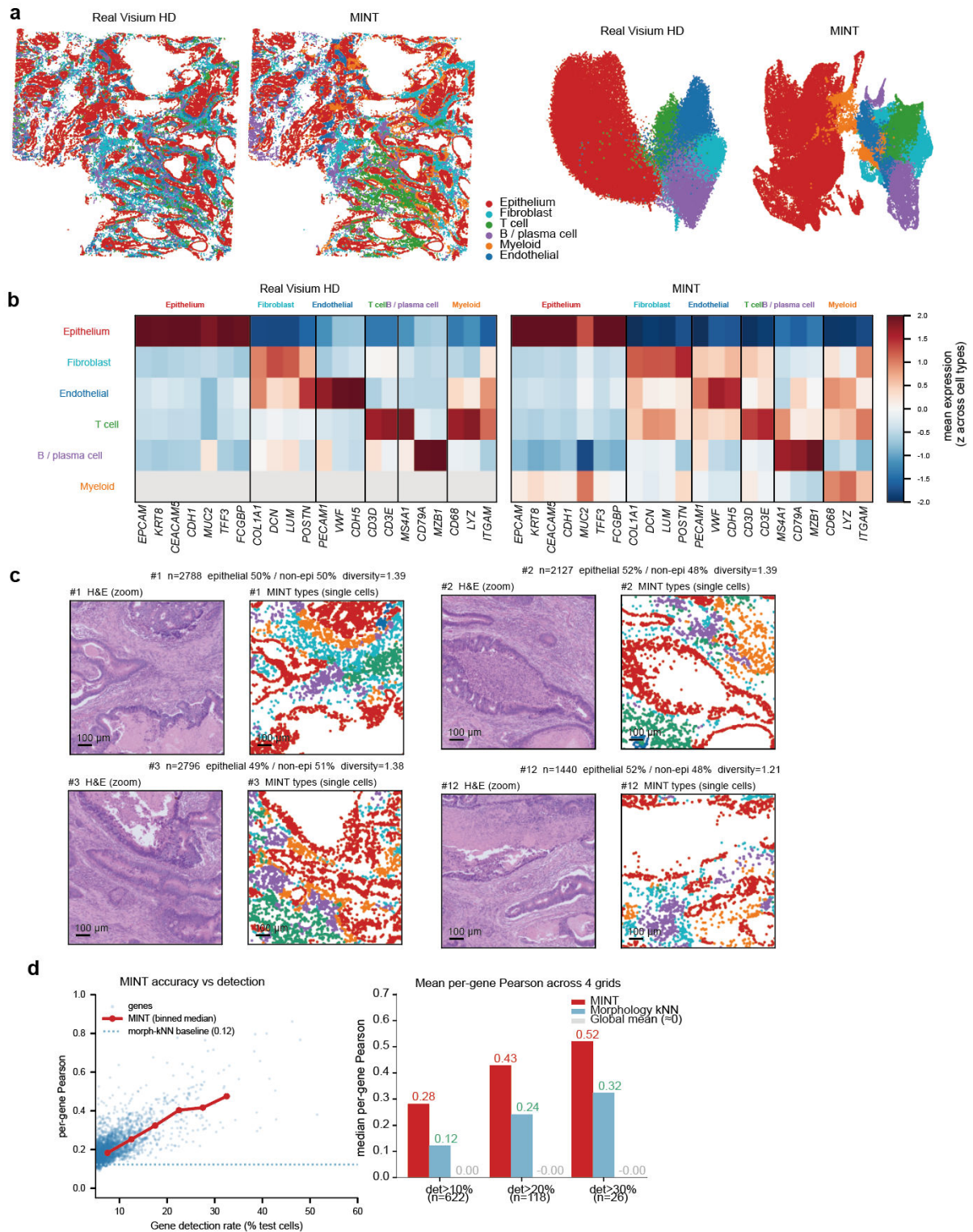

**Supplementary Fig. S14 Visium HD human colon cancer: HD versus MINT clustering, myeloid recovery and per-gene accuracy.** **a**, Spatial cell-type maps and UMAP embeddings for the raw Visium HD measurement and the MINT prediction (six cell types: epithelial, fibroblast, T cell, B/plasma, myeloid and endothelial). **b**, Cell-type marker-signature heatmaps for the raw Visium HD measurement (left) and the MINT prediction (right). **c**, Zoomed regions of interest comparing H&E with MINT single-cell-type maps. **d**, MINT per-gene accuracy against gene detection rate (left; red, MINT binned median; dotted line, morphology nearest-neighbour baseline), and median per-gene Pearson correlation for MINT, the morphology nearest-neighbour baseline and a global-median predictor at three detection thresholds (right).

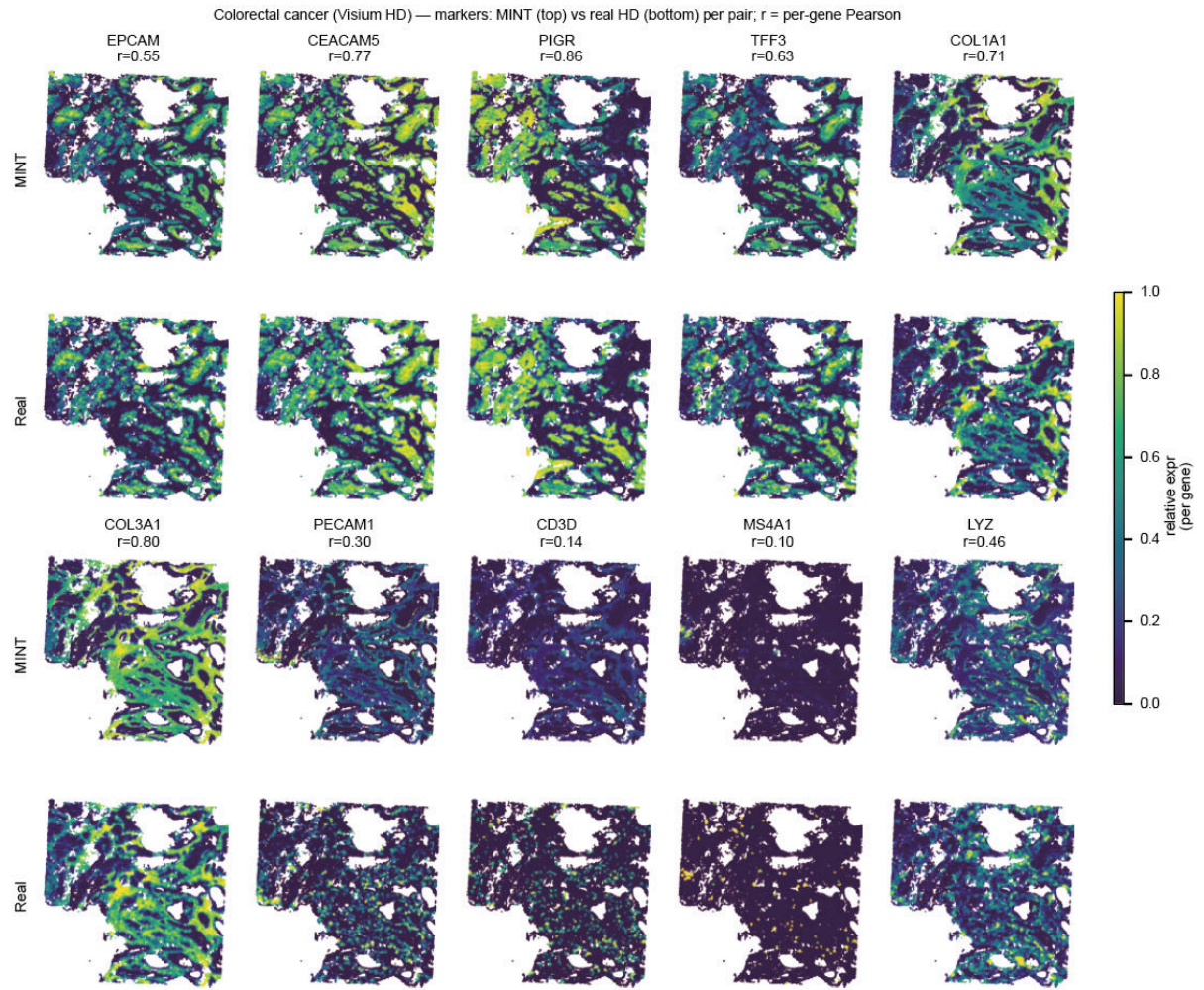

**Supplementary Fig. S15 Visium HD human colon cancer: per-gene marker validation.** Spatial maps of representative markers comparing MINT predictions with HD measurements, with per-gene Pearson correlations annotated (for example, *PIGR* 0.86, *COL3A1* 0.80 and *CEACAM5* 0.77).

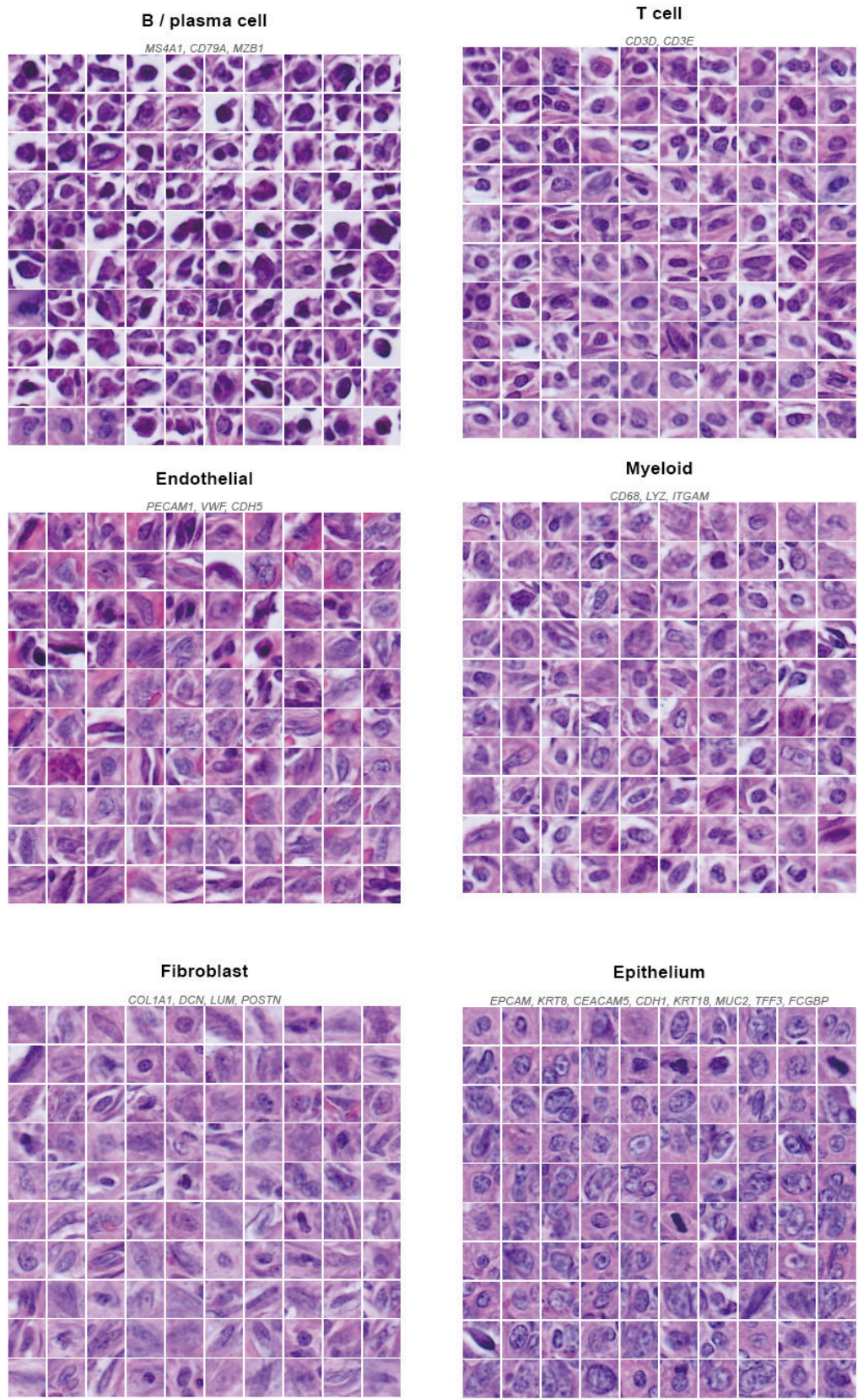

**Supplementary Fig. S16 Visium HD human colon cancer: cell-type morphology gallery.** Nuclear-crop montages of six MINT-predicted cell types (epithelium, fibroblast, T cell, B/plasma, myeloid and endothelial), selected by marker expression.

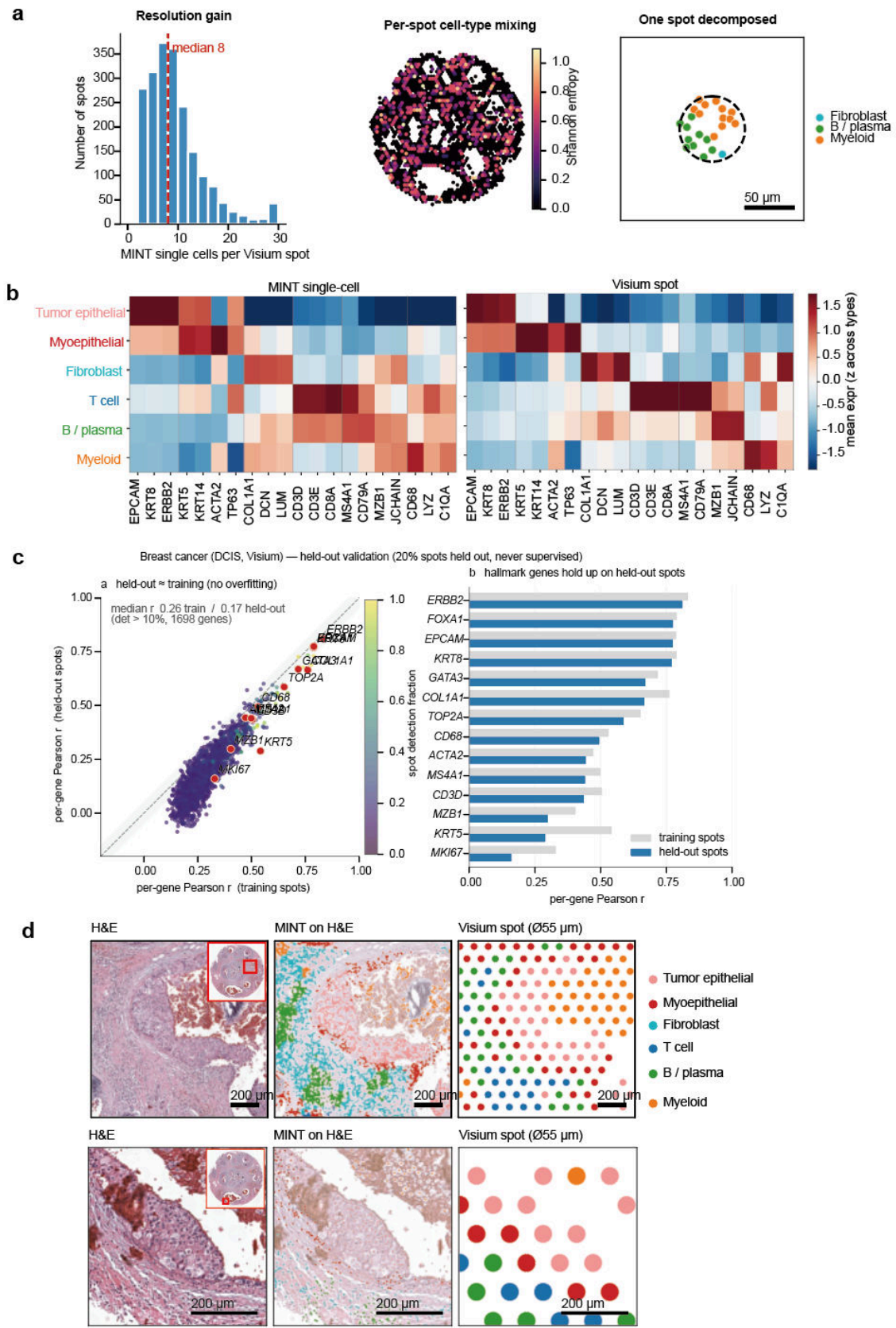

**Supplementary Fig. S17 Real Visium human breast DCIS: resolution gain and held-out validation.** MINT single cells per Visium spot, per-spot cell-type mixing entropy, an example decomposed spot, a cell-type marker heatmap comparing MINT with Visium spots, held-out spot validation, and H&E views comparing MINT single cells with 55-μm spots.

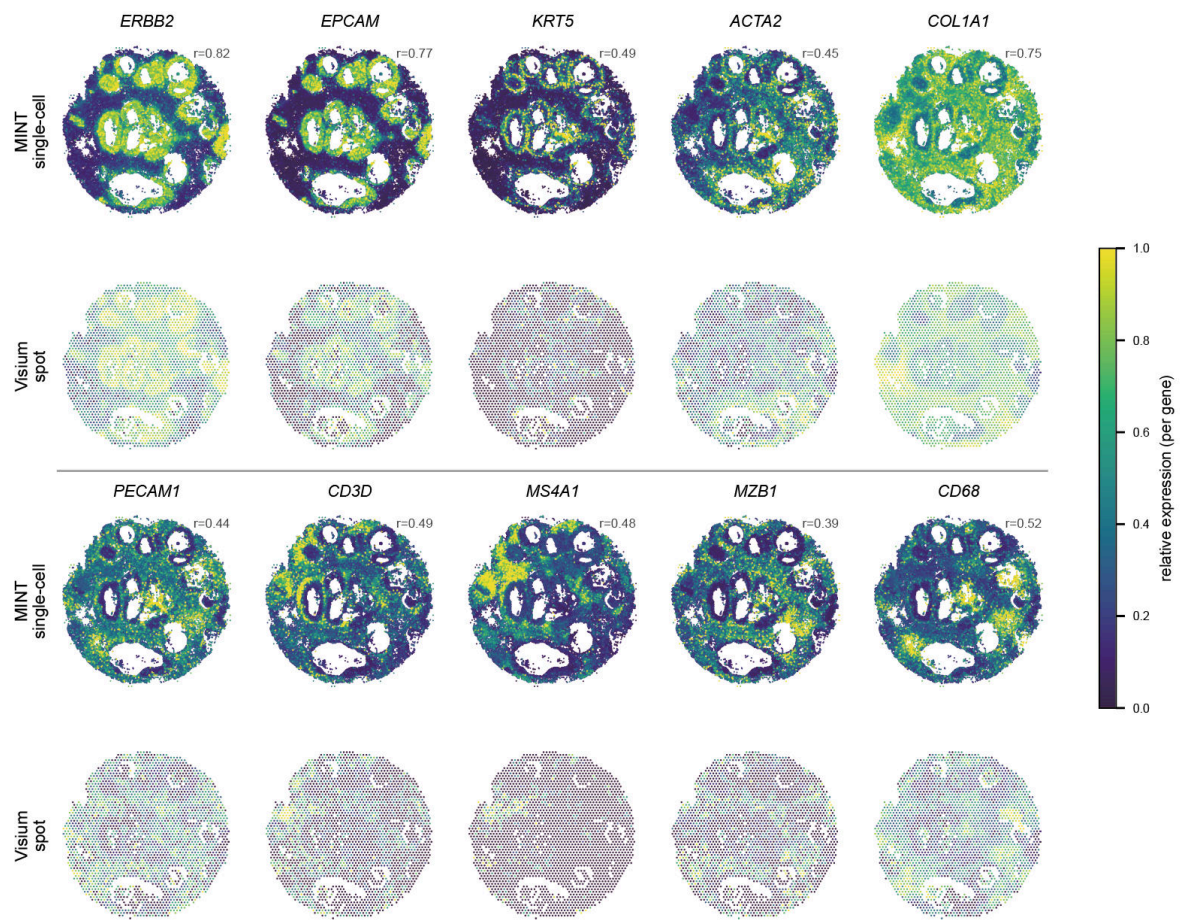

**Supplementary Fig. S18 Real Visium human breast DCIS: per-gene marker maps.** Spatial maps of ten markers comparing MINT single-cell predictions with Visium spots, with per-gene Pearson correlations annotated.

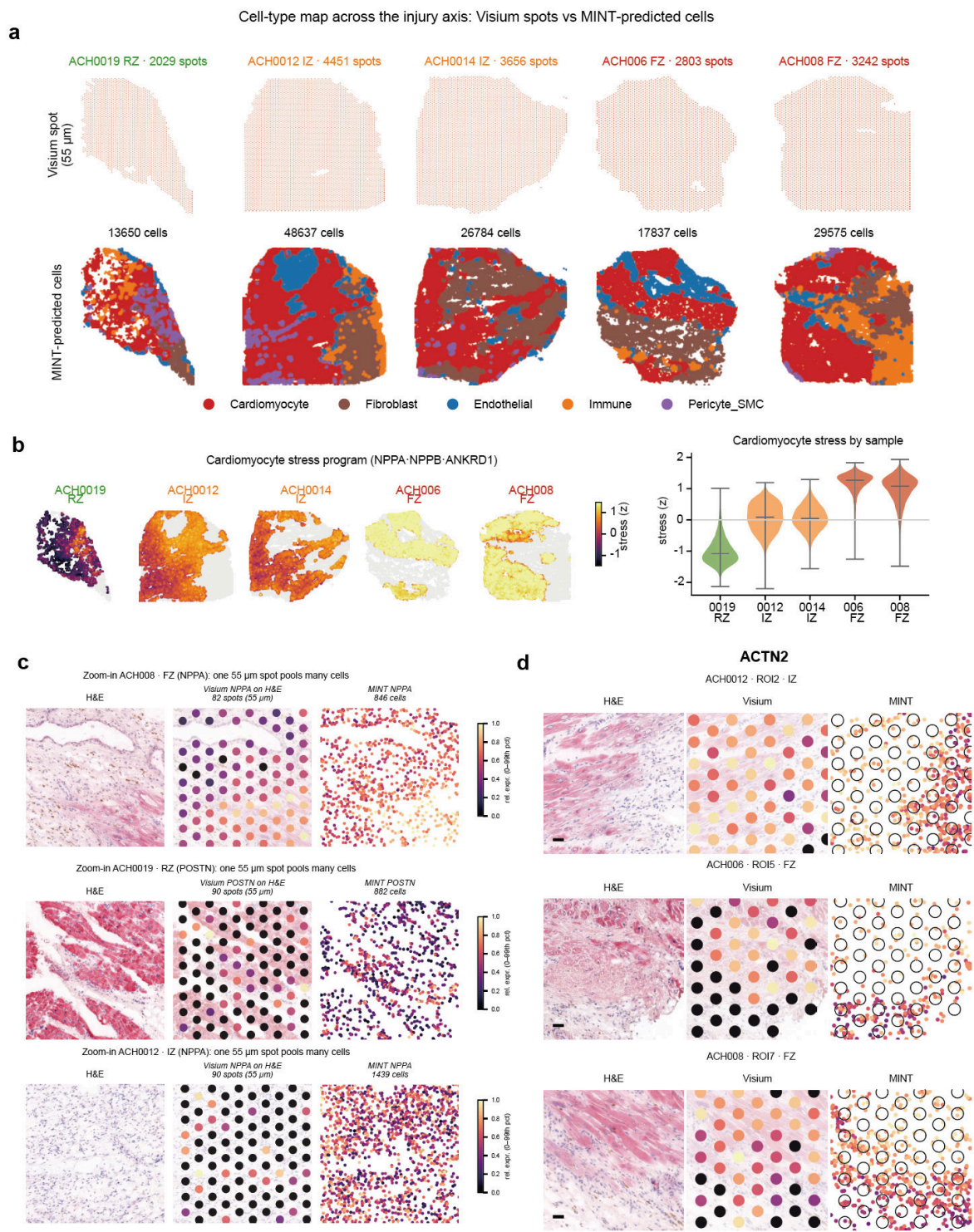

**Supplementary Fig. S19 Real Visium human myocardial infarction.** **a**, Cell-type maps across the injury axis (remote, ischaemic and fibrotic zones; five sections) comparing Visium spots with MINT-predicted cells (cardiomyocyte, fibroblast, endothelial, immune, pericyte/smooth-muscle), with per-section spot and cell counts. **b**, Cardiomyocyte-stress programme (*NPPA*, *NPPB*, *ANKRD1*) as MINT cell-level spatial maps, with the per-section distribution of the programme score (violins; central bar, median). **c**, Zoomed examples in which a single 55-µm spot pools many cells, for the cardiomyocyte-stress marker *NPPA* and the fibroblast-activation marker *POSTN* (H&E, Visium spot and MINT cell-level prediction). **d**, Regions of interest for the cardiomyocyte structural marker *ACTN2* (H&E, Visium spot and MINT cell-level prediction) in three sections.

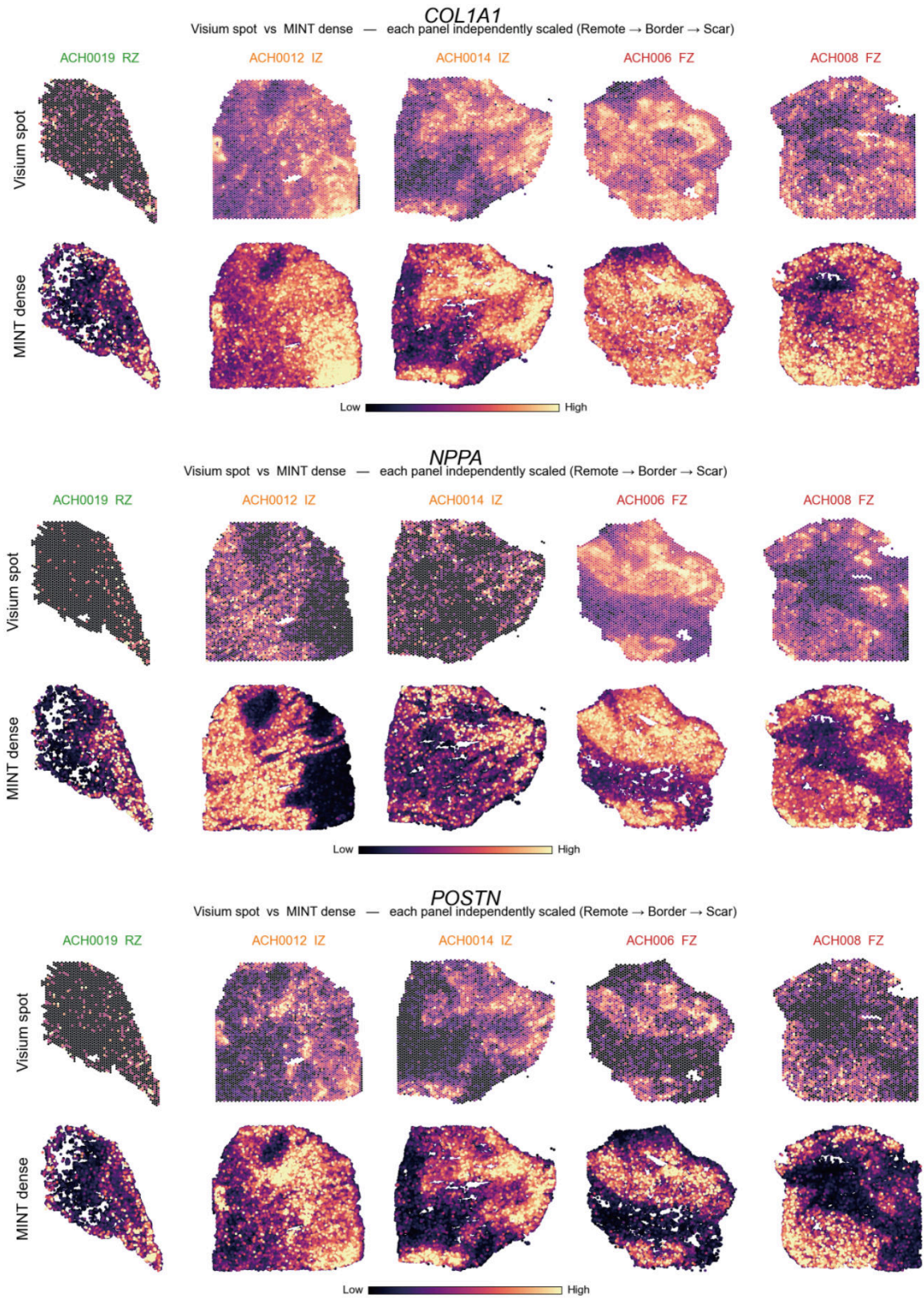

**Supplementary Fig. S20 Real Visium human myocardial infarction: programme-marker maps at spot and cell level.** Spatial maps of the fibroblast-activation markers *COL1A1* and *POSTN* and the cardiomyocyte-stress marker *NPPA* across five sections spanning remote (RZ), ischaemic (IZ) and fibrotic (FZ) zones. Each panel compares Visium spot input with the MINT cell-level prediction and is independently scaled.

### Supplementary Tables

**Supplementary Table S1.** *Datasets used in this study.* Source dataset for every benchmark and validation in the manuscript: platform, tissue, species, gene panel, cell or spot count, accession URL, citation key, and which figure/section consumes it. (File: `supp_table_S1_datasets.csv`.)

**Supplementary Table S2.** *Cross-tissue single-cell benchmark on pseudo-Visium.* Per-tissue, per-method single-cell benchmark of MINT against sCellST, Thor and iStar across the five tissue sections (mean and standard deviation across genes of the per-gene Pearson correlation and RMSE, together with a two-sided Wilcoxon signed-rank test of each method against MINT across genes; one row per tissue and method). Because genes are strongly co-expressed and not independent, the tabulated $p$-values are descriptive paired summaries rather than calibrated inferential significance; the mean accuracy metrics provide the primary comparison. All values are computed on the same H&E-segmented nuclei against Xenium single-cell ground truth, with predicted cell $i$ matching Xenium cell $i$ directly, in a common CP10k+log1p space. Thor did not complete on the renal carcinoma section ($\sim 388,000$ cells), so its renal carcinoma entries are left as dashes (–). (File: `supp_table_S2_benchmark_summary.csv`.)

**Supplementary Table S3.** *Cross-stage prediction and matching metrics on the human skin melanoma dataset.* Per-stage performance ($\pi_0/\pi_1/\pi_2/\text{GT}$) on Xenium Human Skin Melanoma at 80% nuclear loss. Prediction fidelity is summarised on the 50 spatially variable genes held out from alignment (ranks 51–100 by Moran’s $I$, disjoint from the top-50 alignment panel, so these evaluation genes never guided the matching) by per-gene Pearson (reaching $r \approx 0.67$ at $\pi_2$) and per-gene normalised RMSE (RMSE divided by the 1–99 percentile range of log1p ground-truth expression); per-gene Pearson on the full panel is also tabulated but is diluted by the panel’s many low-count genes. Matching quality is given by Top-50 matching accuracy and mean matching distance. GT denotes the ground-truth-alignment reference in which each lost nucleus is matched to its true counterpart. (File: `supp_table_S3_skin_metrics.csv`.)

**Supplementary Table S4.** *Sample-level metadata for the in-house clinical Slide-tags cohort.* Lung adenocarcinoma (P1–P6) and glioblastoma (GL1C, GL1P, GL2C, GL2P, GL3C) sections, each profiled by Slide-tags with an adjacent-section H&E image. Captured nuclei = Slide-tags-sequenced nuclei; segmented nuclei = H&E-segmented nuclei (the MINT reconstruction target); UMI per captured nucleus (median) = per-nucleus sequencing library size. MINT reconstructs the whole transcriptome for every segmented nucleus. The lung cohort’s UMI-per-nucleus median spans $\sim 2,800$–$9,000$ ($\sim 3$-fold across samples), yet the TLS-bearing samples are not the most deeply sequenced (P1 at 4,794 versus the immune-cold P4 at 8,999), consistent with tertiary-lymphoid-structure calls resting on spatial organisation rather than sequencing depth (Methods). For the lung cohort, sex, age, smoking status and CT nodule size are tabulated; pathological stage is given by the diagnosis (adenocarcinoma in situ, Tis; minimally invasive adenocarcinoma, T1mi), all cases were node-negative or benign, and no clinical correlation is drawn from the single section available per patient. The lung sample identifiers in the *Sample* column are the section identifiers deposited under OMIX accession OMIX018434 (Data availability). (File: `supp_table_S4_clinical_cohort.csv`.)

**Supplementary Table S5.** *Distribution-level fidelity and diversity of the generative Slide-tags reconstruction.* For each Slide-tags reconstruction we report the per-gene Kolmogorov–Smirnov distance between MINT’s reconstructed single-cell distribution and a reference distribution (averaged over genes; lower is closer), together with the effective dimensionality (participation ratio of the principal-component variance, a proxy for single-cell diversity) of the reference and of the MINT reconstruction, computed by the identical procedure. For the pseudo-Slide-tags datasets the reference is the held-out true single cells (Xenium): MINT’s reconstruction closely matches the reference distribution ($\text{KS} \leq 0.014$) and recovers a comparable diversity that sits slightly below the ground truth rather than inflating it. For the real Slide-tags datasets no single-cell ground truth exists, so the reference is the sparse measured captured nuclei, which under-sample the tissue’s diversity; the dense reconstruction therefore shows a somewhat higher effective dimensionality than this sparse reference. (File: `supp_table_S5_generative_fidelity.csv`.)

**Supplementary Table S6.** *Cross-slice transfer benchmark (breast cancer replicates).* MINT, sCellST and iStar were trained on one breast cancer replicate and evaluated on the held-out replicate’s Xenium single-cell ground truth, in both directions (Rep1→Rep2 and Rep2→Rep1). MINT used the same spot-level configuration as in Supplementary Table S2, with supervision derived from the source replicate’s pseudo-Visium spot aggregates. For each method and direction we report the mean per-gene Pearson correlation, the mean per-gene RMSE and the mean cell-wise profile Pearson correlation, all computed on the same H&E-segmented nuclei (predicted cell $i$ matched to Xenium cell $i$ directly) in a common CP10k+log1p space. Thor is omitted because it propagates measured expression across a within-section similarity graph and does not learn a mapping that transfers to a new section. MINT attains the highest mean per-gene Pearson and mean cell-wise profile correlation in both directions; its mean per-gene RMSE

differs from sCellST by less than 2% in both directions. Both nucleus-centred methods exceed the grid-based iStar. (File: `supp_table_S6_cross_slice_transfer.csv`.)

**Supplementary Table S7.** *Marker panels and signatures for the clinical immune-microenvironment analysis.* Gene sets underlying the lung Slide-tags cell-type annotation and the tertiary-lymphoid-structure (TLS) metrics. *Cell-type annotation panels:* the per-type marker sets scored in absolute CP10k+log1p units for discrete cell-type calling by marker dominance (B excludes *BANK1*, a noisy probe). *TLS spatial-metric lineage scores:* the compact per-cell lineage means driving B-follicle focality, FDC fold-enrichment and lineage positive fractions. *TLS signatures:* the breast-panel TLS signature (Fig. 5d), scored with `score_genes` on the 7 of 12 genes present in the 313-gene breast panel, and the lung B-follicle/germinal-centre signature (Extended Data Fig. 3b), each gene min-max scaled to [0, 1] and averaged across genes. (File: `supp_table_S7_lung_markers_signatures.csv`.)
